# Mechanosensitive genomic enhancers potentiate the cellular response to matrix stiffness

**DOI:** 10.1101/2024.01.10.574997

**Authors:** Brian D. Cosgrove, Lexi R. Bounds, Carson Key Taylor, Alan L. Su, Anthony J. Rizzo, Alejandro Barrera, Gregory E. Crawford, Brenton D. Hoffman, Charles A. Gersbach

## Abstract

Epigenetic control of cellular transcription and phenotype is influenced by changes in the cellular microenvironment, yet how mechanical cues from these microenvironments precisely influence epigenetic state to regulate transcription remains largely unmapped. Here, we combine genome-wide epigenome profiling, epigenome editing, and phenotypic and single-cell RNA-seq CRISPR screening to identify a new class of genomic enhancers that responds to the mechanical microenvironment. These ‘mechanoenhancers’ could be active on either soft or stiff extracellular matrix contexts, and regulated transcription to influence critical cell functions including apoptosis, mechanotransduction, proliferation, and migration. Epigenetic editing of mechanoenhancers on rigid materials tuned gene expression to levels observed on softer materials, thereby reprogramming the cellular response to the mechanical microenvironment. These editing approaches may enable the precise alteration of mechanically-driven disease states.

---

The cellular microenvironment is a potent regulator of cellular behavior (*1*, *2*). Stimuli from the microenvironment are often classified as chemical or mechanical in nature. Chemical stimuli regulate nearly every fundamental cell process by activating signaling pathways that ultimately converge in the nucleus to alter the epigenetic state of the cell and govern transcriptional networks. Recent studies have applied newly developed CRISPR screening approaches and epigenetic profiling to determine how many aspects of the cellular microenvironment alter the epigenetic landscape, including the investigation of chemical stimuli such as hormones and cytokines (*3*–*6*). Mechanical stimuli from the microenvironment, such as stiffness of the extracellular matrix (ECM) or applied mechanical forces, are also potent regulators of many fundamental cell processes, including growth, death, differentiation, and migration (*7*–*10*). However the mechanisms through which mechanical stimuli are converted into precise patterns of gene expression are not as well-characterized or understood at the genomic level.

The most well-described mechanosensitive mechanisms of gene regulation are associated with either nucleocytoplasmic shuttling of transcription factors and co-activators or epigenetic changes that alter chromatin accessibility to promote downstream transcription (*11*). For instance, mechanical stimuli can inhibit the Hippo signaling pathway to prevent the phosphorylation of the transcriptional co-activator YAP, facilitating its transport to the nucleus where it interacts with TEAD transcription factors to alter gene expression (*12*–*14*). Similarly, mechanosensitive signaling pathways that promote F-actin polymerization release the transcriptional co-activator MRTF from sequestration by G-actin, thereby facilitating its transport to the nucleus where it interacts with SRF transcription factors to regulate genes (*15*–*17*). These mechanical signals can also regulate nuclear transport by stretching the nuclear pore complex (NPC) in a process facilitated by Ran GTPases (*18*) that also leads to translocation of transcriptional activators including YAP (*19*). Mechanical cues can drive epigenetic changes in the nucleus through both indirect and direct regulation. For example, indirect epigenetic signaling occurs via mechanical stretching of cells that leads to epigenetic reinforcement by regulating methyltransferases and/or acetyltransferases and subsequent H3K27me3 deposition that modulates transcription (*20*–*23*). Additionally, exogenous mechanical force may directly influence DNA accessibility, whereby applied force to the cell leads to chromatin deformation that correlates with bursts of transcription (*24*, *25*).

Traditionally, the binding of transcription factors and co-activators at promoter regions has been the classical mechanism through which proteins in the genome are thought to influence transcription of target genes. However, genome annotation efforts over the last two decades have shown that gene regulatory regions occur predominantly outside promoters and are frequently located within non-coding genomic regions (*26*). For instance, the mechanosensitive co-activators YAP/TAZ preferentially bound to distal non-coding genomic regions in cells cultured on rigid tissue culture plastic (*27*, *28*). One particularly well-studied class of distal genomic regulatory elements are enhancers, which act across variable genomic distances to regulate transcription and are often marked by a combination of chromatin accessibility (e.g., DNase-seq, ATAC-seq), presence of active histone marks (e.g., H3K27ac), depletion of repressive histone marks (e.g., H3K9me3), TF binding, and chromatin looping to distal target genes (*29*). The complex logic of gene regulation by these distal elements has been notoriously difficult to dissect, but advances in high-throughput CRISPR screening and single-cell genomics have transformed the capability to classify how and where these *cis*-regulatory elements modulate transcription across the genome (*4*, *5*, *30*, *31*).

Despite these recent developments, we still do not fully understand how and where mechanical stimuli affect the non-coding genome to regulate transcriptional responses that ultimately determine key cell phenotypes. Here, we utilized genome-wide profiling of chromatin accessibility along with epigenetic editing, high-throughput CRISPR screening, and single-cell sequencing tools to characterize how ECM stiffness cues activate *cis*-regulatory elements to regulate gene expression. Through this work we identify and validate a novel set of *cis*-regulatory elements that are responsive to changes in the mechanical microenvironment. For simplicity, we term these regions as ‘mechanoenhancers’, and show they behave as key drivers for downstream mechanically-driven behaviors in human cells.

## Results

### Widespread changes in chromatin accessibility result from short-term exposure to physiologically soft or stiff substrates

We first characterized the response of gene expression and chromatin structure to changes in ECM stiffness cues by culturing primary human neonatal foreskin fibroblasts (HFF cells) on substrate stiffness conditions that represent a wide range of pericellular niches across various tissues in health and disease. Fibroblasts were chosen for these analyses because they have a wide range of available functional genomics data, play a key role in ECM synthesis, and can contribute to disease states in tissue fibrosis (*32*). HFF cells were cultured for 20 hours on either soft (Elastic modulus, E = 1 kPa, mimicking the softest connective tissues) or stiff (E= 50 kPa, mimicking organized musculoskeletal tissues or fibrotic lesions) polyacrylamide hydrogels (*9*, *33*, *34*) as well as on tissue culture plastic (TCP, E= ∼1 GPa). The 20-hour time point minimizes transcriptional feedback that could further complicate understanding the direct influence of ECM stiffness on epigenetic state. Following 20 hours of culture on the soft or stiff hydrogels, HFF cells were harvested to examine both transcriptional changes (RNA-seq) and chromatin accessibility changes (ATAC-seq) in response to these ECM stiffness cues (**Fig. 1A**). We performed all sequencing experiments in at least duplicate per condition, and all RNA-seq and ATAC-seq data were highly reproducible and passed quality control metrics established by the ENCODE Consortium (**Fig. S1, Table S1**). Transcriptomic analysis identified 1,535 significant differentially-expressed genes (**Fig. 1B-C, Table S2**). We found significantly increased expression of genes related to changes in ECM remodeling, apoptotic factors, anti-fibrotic programs, and cell lineage establishing factors on soft hydrogels, while canonical genes associated with YAP/TAZ translocation and pro-fibrotic programs were significantly upregulated on stiffer materials (**Fig. 1C),** similar to previous studies (*35*, *36*).

**Fig. 1.**
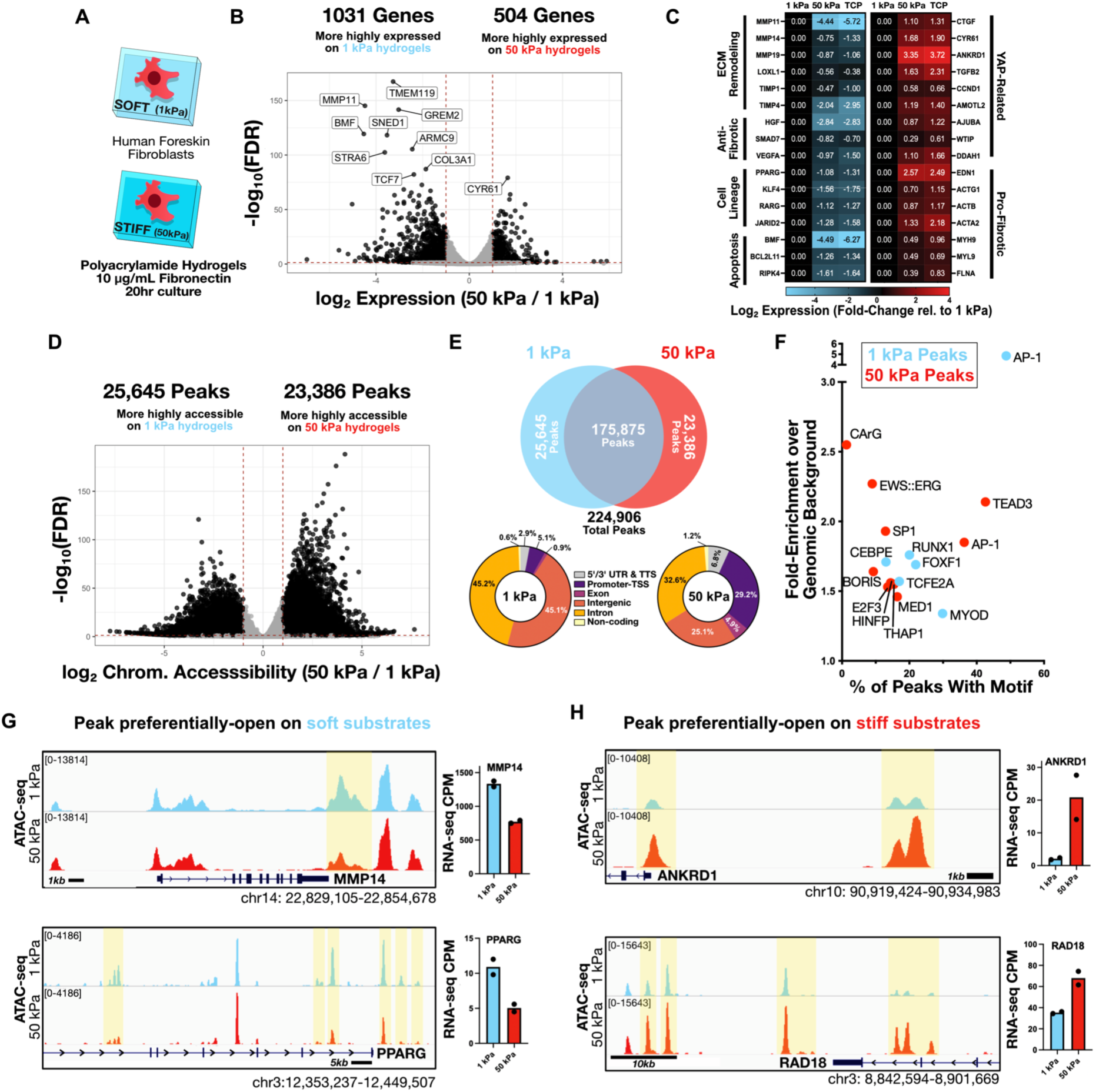
Short-term culture on physiologically-soft materials results in broad changes in gene expression and chromatin structure. **(A)** To examine the influence of physiologically-soft mechanical microenvironments on the cellular epigenetic state, primary human neonatal fibroblasts (HFF cells) were cultured on soft (Elastic modulus, E=1kPa) or stiff (E= 50kPa) fibronectin-coated polyacrylamide hydrogels for 20 hours. **(B)** RNA-seq revealed differentially-expressed genes (FDR < 0.01, abs(Log2 Fold-Change)>1), with clusters of differentially-expressed genes highlighted in **(C). (D)** ATAC-seq revealed differentially-accessible chromatin regions (FDR < 0.01, abs(Log2 Fold-Change)>1) from HFF cells cultured on either soft or stiff hydrogels. **(E)** Overview and annotations of differentially-accessible chromatin regions across HFF cells cultured on either 1 kPa and 50 kPa hydrogels from the Top 5000 most significantly changing peaks over UTR/TTS, promoter-TSS, exon, intergenic, intronic, or non-coding RNA annotations. **(F)** Significantly enriched *de novo* motifs from differentially-accessible regions on 1 kPa or 50 kPa substrates. (**G-H**) ATAC-seq tracks showing representative regions with significantly lower (MMP14, PPARG) or higher (ANKRD1, RAD18) chromatin accessibility on stiff hydrogels.

We next compared changes in chromatin accessibility between the soft and stiff hydrogel conditions by ATAC-seq. Following 20 hours of culture on these materials, we observed widespread changes in chromatin accessibility, with ∼21% of total identified accessible chromatin peaks having significantly differential accessibility between conditions, with an even balance of peaks that were more accessible on soft or stiff materials (**Fig. 1D-E, Table S3**). Sites that were more accessible on the stiff hydrogel had a fivefold increase in the frequency of peaks located in promoter regions as compared to the peaks that were more accessible on soft hydrogels (**Fig. 1E)**, in accordance with previous observations of increased mechanoresponsive TF shuttling into the nucleus (*19*). To further understand which TF signaling modules might be mediating these changes in accessibility, we performed *de novo* TF motif analysis in the entire set of differentially-accessible peaks for both material conditions. The peaks with increased accessibility on soft hydrogels were enriched with AP-1 motifs compared to genomic background. Peaks more accessible on stiff hydrogels were enriched with TEAD, AP-1, SP1, and CaRG motifs (**Fig. 1F**), as has been previously observed for mechanoresponsive pathways (*27*, *37*–*39*). Interestingly, the *de novo* motifs identified for AP-1 were markedly different between soft and stiff materials (**Fig. S2**). Different motif usage can be associated with specific AP-1 complexes across different AP-1 family members (*40*). Our findings of different *de novo* AP-1 motif enrichment in the soft and stiff materials suggest that AP-1 sub-unit formation might differ across these contexts. We found that stiffness-dependent accessibility changes often mapped near canonical mechanosensitive genes (*9*, *27*, *41*, *42*), including increased accessibility on 1 kPa hydrogels near *MMP14* and *PPARG* (**Fig. 1G**), and increased accessibility on 50 kPa hydrogels near the YAP targets *ANKRD1* and *RAD18* (**Fig. 1H**). For all of these examples, increased accessibility correlated with increased expression across the stiffness conditions (**Fig. 1G-H**).

We next explored the short-term reversibility of changes in accessibility following changes in intracellular acto-myosin contractility. Rho-associated protein kinases (ROCKs) are regulators of both acto-myosin contractility and actin organization in the cell, and are a key driver of the cellular sensing of matrix stiffness cues (*9*) that are relevant to cellular growth, migration, and apoptosis. We cultured HFF cells on stiff (50 kPa) hydrogels overnight, and then treated the cells with either DMSO or 10 µM of the ROCK inhibitor Y-27632 (ROCKi) for 1 hour prior to harvest for ATAC-seq directly on the culture substrates. This allowed for the reduction of cell contractility while avoiding other changes associated with cell trypsinization. We observed 2,052 peaks with differential accessibility relative to DMSO-treated cells cultured on the same surface, demonstrating that cell contractility is required for the maintenance of the stiffness-induced changes and that downstream remodeling of chromatin accessibility could happen within an hour of contractile changes (**Fig. S3, Table S4**). This is also consistent with the previous observation that external mechanical stimuli and increases in intracellular contractility drive similar mechanosensitive processes (*43*).

### An intronic mechanoenhancer increases *MYH9* expression on stiff materials

As cell contractility was required for the maintenance of changes in chromatin accessibility observed in stiff hydrogels, we next investigated cis-regulation of the non-muscle myosin genes *MYH9, MYH10, and MYH14*. These genes encode for non-muscle myosin IIA IIB, and IIC respectively, which are the primary drivers of cellular contractility in non-muscle cells (*44*). In primary HFF cells, *MYH9* is the predominantly expressed non-muscle myosin and the only non-muscle myosin that showed ECM stiffness-dependent changes in expression (**Fig. 2A**). Our ATAC-seq analysis identified 14 regions that were differentially-accessible between soft and stiff substrates that mapped within 100kb of the *MYH9* transcriptional start site (TSS). We sought to test if any of these 14 stiffness-dependent peaks near *MYH9* functioned as stiffness-induced modulators of *MYH9* expression. To perturb the epigenetic state at any specific genomic locus we utilized CRISPR interference (CRISPRi) with the dCas9^KRAB^ epigenome editor. dCas9^KRAB^ catalyzes the addition of repressive histone marks at the target site (e.g., H3K9me3) along with the removal of active histone marks (e.g., H3K4me3/H3K27ac) to decrease chromatin accessibility and induce epigenetic silencing (*45*, *46*). We first performed a CRISPRi screen using dCas9^KRAB^ combined with a gRNA library tiling all ATAC-seq peaks in HFF cells (regardless of if they were mechanically-sensitive) within +/- 440 kb of the *MYH9* TSS (114 regions, 5,192 gRNAs) (**Fig. 2B**). To identify regions regulating *MYH9* expression, cells were fixed and stained for MYH9 (NMIIA), sorted into *MYH9*-high and *MYH9*-low expression bins, and compared for their distributions of gRNAs (**Supplementary Text 1, Fig. S4, Tables S5-6**, **Methods**). Across these gRNAs, we identified five putative regulatory elements (pREs) as strong regulators of MYH9 protein expression including two pREs in the *MYH9* promoter/TSS region and three pREs within a ∼5 kb section of intron 3 of *MYH9* (**Fig. 2C-D**). Of the three pREs in intron 3, only the first pRE was differentially accessible between the soft and stiff hydrogel culture conditions (**Fig. 2E**). Further examination of H3K27ac signatures across diverse ENCODE biosamples around the sub-region of differential accessibility showed low signal in suspension or weakly adherent cell lines (e.g., K562 cells), but greater signal across increasingly adherent and contractile cell lines (e.g., HUVEC/HSMM, **Fig. 2F**). Thus, as the stiffness of the culture environment increases, canonical indicators of enhancer activity also increase, suggesting that the activity of the *MYH9* pRE#1 is responsive to mechanical cues across cell types.

**Fig. 2.**
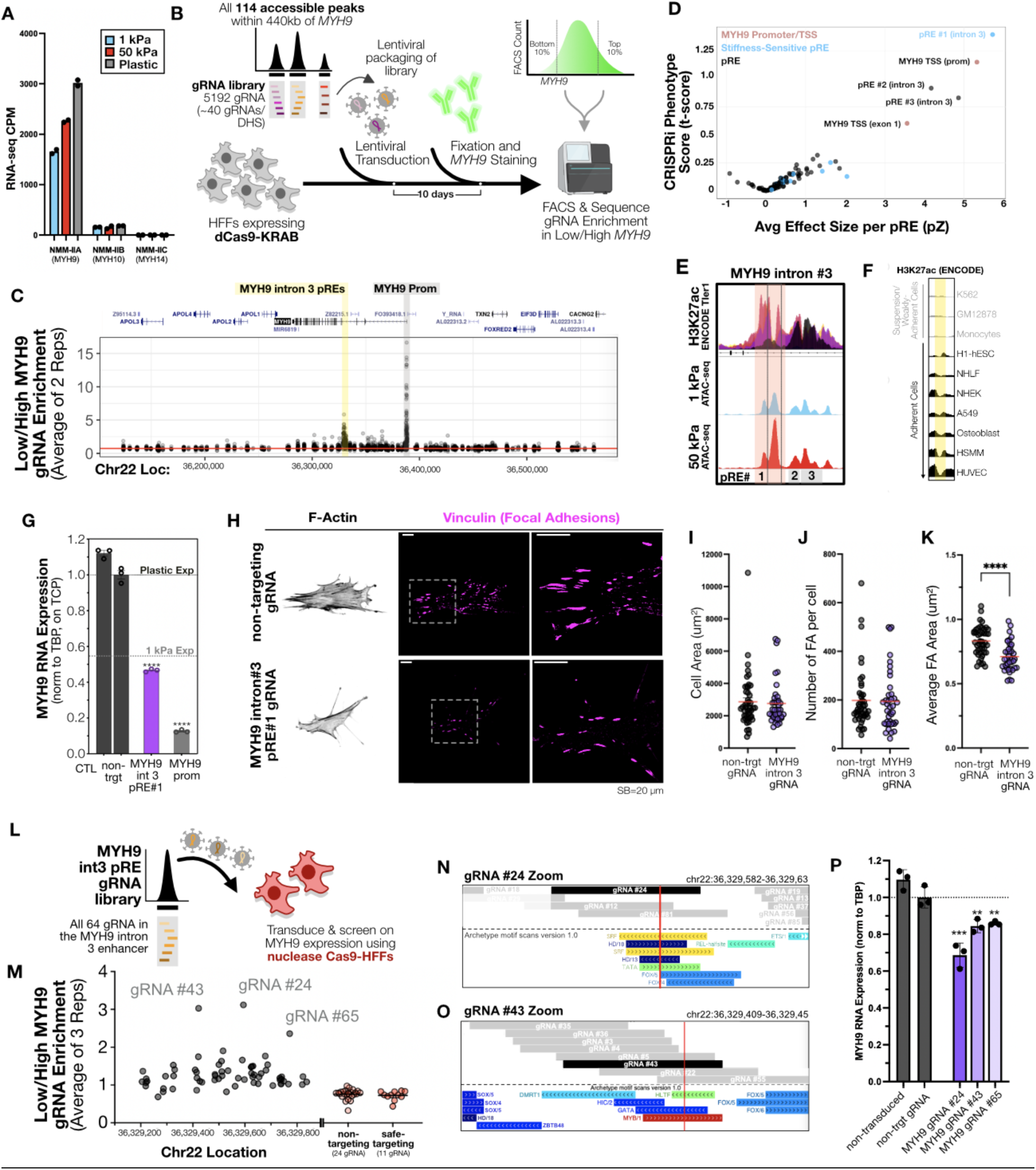
CRISPRi screening reveals a MYH9 intron 3 mechanoenhancer that regulates MYH9 expression and cell contractility. **(A)** Expression of MYH9 on soft 1 kPa hydrogels, 50 kPa hydrogels, and TCP from RNA-seq (N=2 reps/group) **(B)** Schematic of CRISPRi screening procedure for finding genomic regulators of MYH9 protein expression. **(C)** Individual gRNA enrichment in Low/High MYH9 expression bins following the *MYH9 locus* screen averaged across two replicates. **(D)** CRISPRi screening results across the MYH9 locus as shown by MYH9 Repression Phenotype Scores (t-score) and average effect size (z-score) as calculated across each DHS in the screen. Blue points indicate DHS was differentially-accessible in ATAC-seq data between soft/stiff hydrogel conditions across both screen replicates. **(E)** ATAC-seq signal across the MYH9 intron 3 enhancer region, with the yellow highlight denoting the force-sensitive pRE#1 subregion. **(F)** Normalized ENCODE H3K37ac signal around differentially-accessible pRE#1 peak from MYH9 intron 3 region compared for 9 available ENCODE tier 1 cell lines. **(G)** Relative MYH9 RNA expression 10d following lentiviral transduction with dCas9^KRAB^ along with either a non-targeting gRNA, MYH9 intron 3 enhancer-targeting gRNA, or an MYH9 promoter-targeting gRNA. CTL group represents no transduction. **(H)** Representative F-actin and vinculin focal adhesion immunostaining images and corresponding quantifications (**I-K**) of focal adhesion morphologic parameters in HFF cells transduced with either a non-targeting or a MYH9 intron 3 enhancer-targeting gRNA (N=39-45 FA/group, ** = p <0.01, **** = p<0.0001 by Student’s t-test). Red line indicates group means. **(L)** Schematic of Cas9 nuclease saturation indel screening procedure performed in HFF cells across the MYH9 int3 enhancer region. **(M)** Plot showing the results of this Cas9 screening, with the ratio of gRNA enrichment in low MYH9 expression bins as compared to high MYH9 expression bins across the MYH9 intron 3 enhancer as well as across non-targeting gRNAs and ENCODE safe-targeting gRNAs. Dots shown are averages across all three replicates. (**N-O)** gRNA positioning of hit gRNAs relative to the positions of the core SRF CaRG motif (gRNA#24) and HLTF motif (gRNA#43). (**P**) Relative MYH9 expression during singleton validation of the top three gRNA hits from the screen, six days post-transduction (N=3 reps/group).

To validate the screen results, we first delivered single gRNAs targeting the three *MYH9* intron 3 pREs (pREs #1/2/3, two gRNA per pRE) along with dCas9^KRAB^ via lentivirus and analyzed *MYH9* mRNA expression by qPCR, revealing all three of the *MYH9* pREs regulated *MYH9* mRNA expression (**Fig. S5**). For follow-up experiments we used the most effective gRNA targeting the stiffness-sensitive pRE#1. At nine days post-transduction, targeting the MYH9 intron 3 pRE#1 led to ∼54% repression of *MYH9* transcript levels compared to HFF cells that received a non-targeting gRNA and dCas9^KRAB^ (**Fig. 2G**). This degree of change of *MYH9* mRNA levels following epigenetic repression of the *MYH9* intron 3 pRE#1 is consistent with the decreases observed in *MYH9* mRNA expression between cells cultured on TCP and soft 1kPa hydrogels by RNA-seq (**Fig. 2A**, levels marked on **Fig. 2G** graph). When delivering a *MYH9* promoter-targeting gRNA along with dCas9^KRAB^, we saw ∼87% repression of *MYH9* transcript levels (**Fig. 2G**). Following immunostaining of MYH9 and FACS, we also identified similar fold-changes in MYH9 protein levels 15 days after transduction of the gRNA along with dCas9^KRAB^ (**Fig. S6**). Thus, the *MYH9* intron 3 pRE#1 functions as a mechanoenhancer and dictates *MYH9* expression in response to ECM stiffness cues.

To further identify the influence of the *MYH9* mechanoenhancer on cellular contractility, we assessed changes in cell morphology and in key mechanosensitive machinery 9 days after epigenetic repression of the MYH9 promoter or mechanoenhancer on rigid tissue culture plastic. Vinculin-containing focal adhesions (FAs) are a key mechanoresponsive subcellular structures, and their size and shape are strongly dependent on myosin activity (*44*, *47*). Epigenetic repression of the promoter led to substantial changes in cell size and a near complete loss of vinculin-containing focal adhesions (**Fig. S7**). Epigenetic repression of the MYH9 intronic mechanoenhancer (MYH9 intron 3 pRE#1) did not cause large changes in cell morphology, but altered acto-myosin organization (**Fig. 2H**, **Fig. S7**). Immunostaining with vinculin and FA quantification revealed no significant changes in total cell area (**Fig. 2J**) or total number of FAs per cell (**Fig. 2I**). However, we did observe significantly lower area per FA (**Fig. 2K**), suggesting a lower contractile state of these cells compared to cells that received the non-targeting control gRNA. Collectively, this work identifies an ECM stiffness responsive *MYH9* enhancer in intron 3 that behaves as a mechanoenhancer.

Nuclease-active Cas9 and densely tiled saturating gRNA libraries have been used to determine key motifs involved in enhancer function by introducing a variety of disruptive small insertions and deletions through non-homologous end joining (NHEJ)-based DNA repair (*48*). Sequence changes in cells with a loss of enhancer function are then used to identify key motfis. We adapted this Cas9 screening approach to identify motifs in the MYH9 intron 3 mechanoenhancer that control MYH9 expression. We used a stable HFF-Cas9 cell line and introduced all 64 potential gRNAs tiling across the *MYH9* intron 3 mechanoenhancer, and sorted cells based on MYH9 protein expression as the screen endpoint (**Fig. 2L, Table S7**). We identified three gRNAs that substantially decreased MYH9 expression on TCP relative to the other gRNAs across the mechanoenhancer (**Fig. 2M**). We observed that gRNA #24 and gRNA #43 had cut sites directly overlapping an SRF/CaRG motif and an HLTF (helicase like transcription factor) motif, respectively (**Fig. 2N-O**). Upon delivery of these individual gRNAs, *MYH9* mRNA expression was significantly decreased, with a maximum of ∼30% repression by gRNA #24 (**Fig. 2P**). Cytosolic G-actin ratios regulate the mechanically responsive nuclear shuttling of MRTF-A, which then interacts with DNA-bound SRF to further regulate transcription (*49*). HLTF is a key member of the SWI/SNF complex, which has been implicated in actin-based YAP/TAZ release and subsequent DNA binding (*50*). Together, these results suggest that actin-associated mechanosensitive processes drive MYH9 mechanoenhancer activity.

### An intronic mechanoenhancer of *BMF* is more active on soft materials and is a key driver of the ECM stiffness-driven apoptotic response

Low material substrate stiffness, low adhesion states, and restriction of cell spreading have been shown to increase apoptosis or adipogenesis (*7*, *8*, *51*, *52*). Increased apoptosis due to lack of ECM engagement is termed anoikis (*53*, *54*). A key step in cancer progression is developing anoikis-resistance (*55*). From our RNA-seq data we noted that *BMF*, a key transcriptional effector of anoikis, was strongly upregulated on soft substrates (**Fig. 3A**). We also identified a cluster of three ATAC-seq peaks that were significantly more accessible on soft hydrogels located near *BMF* (**Fig. 3B**). We first used a luciferase reporter of enhancer activity to determine whether these differentially-accessible peaks near *BMF* functioned as putative regulatory elements governing *BMF* transcription. Genomic DNA from all three regions was cloned into the luciferase reporter plasmid, reporter plasmids were transfected into HFF cells cultured on TCP, and luciferase activity was measured 24 hours later. Since *BMF* transcription was increased in the low contractility context of soft materials, we hypothesized that the addition of ROCKi Y-27632 should increase luciferase reporter activity. Only BMF pRE#1 in intron 4 demonstrated any basal enhancer reporter activity on TCP. Following treatment with 10 µM Y-27632, pRE#1 enhancer reporter activity was significantly greater than the activity in DMSO-treated cells, while other regions remained at basal levels. This indicates that pRE#1 enhancer activity is increased in lower contractility environments, further supporting the function of this region in increasing *BMF* expression preferentially on soft substrates (**Fig. 3C**).

**Fig. 3.**
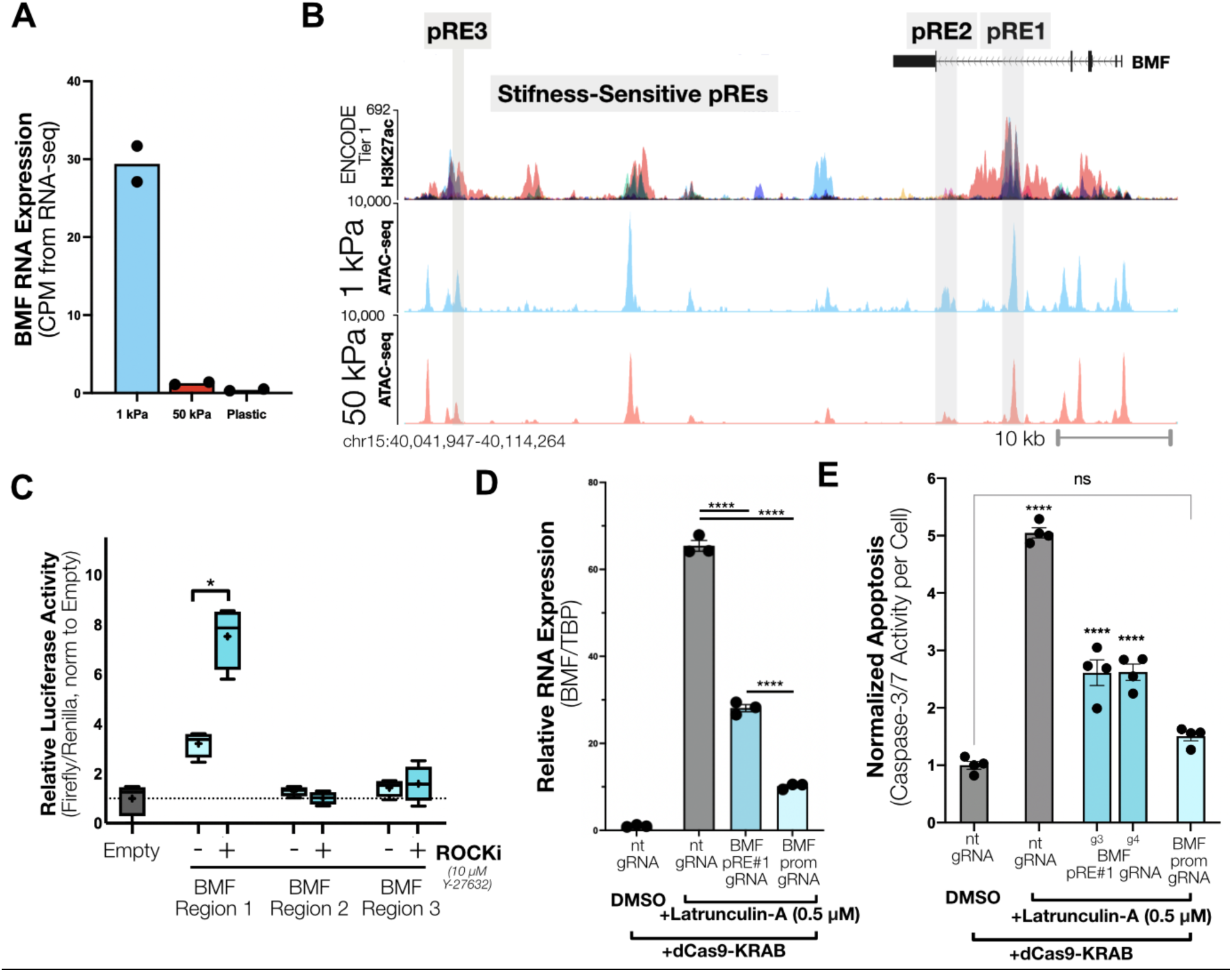
BMF intron #4 mechanoenhancer has increased activity with reduced contractility and is a mediator of anoikis. **(A)** BMF RNA expression from RNA-seq across different stiffness conditions in HFF cells (N=2 reps per stiffness condition). **(B)** ENCODE Tier1 H3K27ac signal and ATAC-seq data between HFF cells cultured on soft and stiff PA hydrogels with regions of differential chromatin accessibility highlighted, with grey highlights indicating regions of differential-accessibility across stiffness contexts. **(C)** Luciferase enhancer reporter readouts from three of the BMF regions with and without 24hr of 10 µM Y-27632 treatment, showing relative firefly luciferase activity controlled by these enhancers normalized to a control co-transfected renilla luciferase reporter. Box and whisker plots show median, plus indicates the group mean, and bars indicate the top/bottom 10% expression range (N=4 reps/group). **(D)** Relative RNA expression (N=3) and **(E)** normalized Caspase-3/7 activity (N=4 reps/group) of HFF cells either untreated, or transduced with various gRNAs, following 0.5 uM LatrunculinA for 24 hours. All data presented as mean +/- SEM and are representative of at least two independent experiments,**** indicates p<0.0001, * indicates p <0.05 by Student’s t-test. Caspase 3-7 activity and luciferase assay stats are shown compared to the DMSO control group, while RNA expression comparisons are shown by overlay bars. “nt gRNA” abbreviates “non-targeting” gRNA.

We next tested the ability of BMF pRE#1 to regulate *BMF* transcription and anoikis. To study this behavior, we utilized a canonical model system for anoikis wherein Latrunculin-A (LatA) treatment is used to depolymerize the actin cytoskeleton to induce loss of FAs and integrin engagement to mimic loss of adhesion to the ECM (*56*). We first transduced HFF cells with dCas9^KRAB^ and either a non-targeting gRNA or a gRNA targeting either *BMF* pRE#1 or the *BMF* promoter. After eight days of culture we evaluated *BMF* mRNA levels. Treatment with LatA increased *BMF* expression ∼60-fold compared to DMSO-treated cells (**Fig. 3D**). Repression of BMF pRE #1 and the *BMF* promoter reduced the LatA-dependent increase in *BMF* expression by ∼60% and ∼85%, respectively (**Fig. 3D**). We then assessed changes in apoptosis by measuring Caspase-3/7 activity using a luciferase reporter system at day 14 post-transduction (8 days +LatA). Repression of BMF pRE #1 reduced LatA-induced cell apoptosis by ∼50% while *BMF* promoter repression completely prevented LatA-induced apoptosis relative to the DMSO-treated control condition (**Fig. 3E**). Collectively, these data show that pRE#1 acts as a mechanoenhancer of *BMF* that is more active on softer ECMs and functions to promote anoikis.

### High-throughput CRISPR screening identifies key mechanoenhancers that modulate cellular growth and migration

To understand which *cis*-regulatory elements contribute most strongly towards mechanosensitive cellular behaviors, we performed high-throughput CRISPRi screening with cellular growth and migration as the phenotypic readouts. We first generated a library of 21,458 gRNAs targeting the top 1000 non-promoter ATAC-seq peaks ranked by increased accessibility on stiff hydrogels (**Tables S3, 8**). We also included gRNAs targeting the promoters of 53 genes that had previously been shown to modulate migration (*57*) as positive controls, and 1000 negative control non-targeting gRNAs. HFF cells were transduced at an MOI of 10.8 to maximize library coverage across a smaller subset of cells and then were assessed for changes in growth or migration (**Fig. 4A**). For the growth screen, genomic DNA was collected on day 8 and day 29/30 (14 population doublings), sequenced, and gRNA enrichment was compared across groups. For the migration screen, at eight days post-transduction we performed a transwell migration assay that allowed cells to migrate overnight. These populations were then separated and used in an additional migration assay the next day. Cell populations that either successfully migrated through the transwell assay twice and those that never migrated through the transwell were collected, genomic DNA was harvested, and gRNA enrichment across populations was determined from sequencing (**Methods**).

**Fig. 4.**
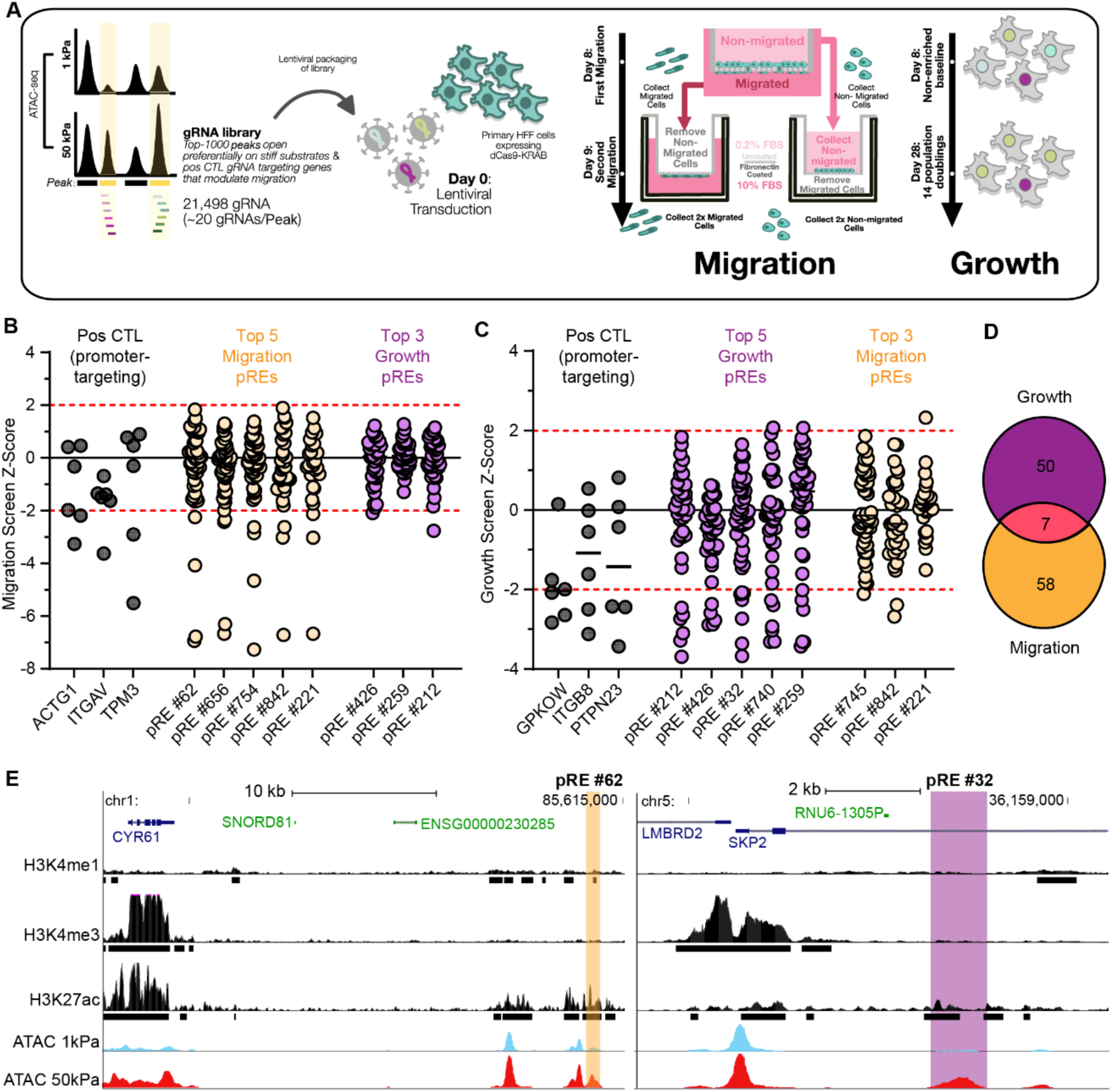
Functional screening of putative mechanosensitive regulatory elements across migration and growth. **(A)** Experimental schema for the paired migration and cellular growth screens in HFF cells transduced with a CRISPRi library of 21,498 gRNA corresponding to the top 1000 differentially-accessible ATAC-seq peaks on stiff substrates. **(B-C)** Migration **(B)** and growth **(C)** phenotype Z-scores for the promoter positive controls, the five pREs with the greatest phenotype Z-scores, and the three representative pREs within the top 5 highest Z-scores for the other phenotype. Each dot represents one gRNA targeting a given pRE. Red dashed line indicates a Z-score threshold of two. **(D)** Venn diagram comparing the pREs that regulated both or only one of the measured phenotypes. **(E)** Intergenic pRE located near CYR61 that regulated migration (left) and pRE located within an intron of SKP2 that regulated growth (right). H3K4me1, H3K4me3, and H3K27ac signal tracks and peak calls in HFF cells are shown. ATAC-seq from HFF cells cultured on soft (1kPa) and stiff (50kPa) are shown below in blue and red, respectively. Note the scaling of H3K4me1, H3K4me3, and H3K27ac, is different from scaling of ATAC-seq.

We observed strong effects from promoter-targeting of the positive control genes. In the growth screen, we found that perturbation of both DepMap essential genes (*GPKOW*, *EIF3E*, *ACTG1*, *CSNK1A1*, *PCYT1A*, *PTPN23*)(*58*, *59*) and genes related to cell growth (*ABL1*, *ITGB8*, *G3BP2*, *OTUD6B*) led to decreases in cell proliferation (*60*, *61*). In the migration screen, we found that the promoter-targeted repression of key genes known to influence cell adhesion and force generation, including *ITGAV*, *ACTG1*, *CDC42,* and *TPM3(57)*, decreased cell migration (**Fig. S8**). We observed that perturbations to mechanically-sensitive test regions led to similar degrees of enrichment as perturbations of positive control genes (**Fig. S8**). When analyzing the distribution of effect sizes for gRNAs across a given peak, we noted strong Z-score enrichment of only a fraction of the gRNAs across both screens (**Fig. 4B-C**), which is consistent with previous reports of epigenetic editing of regulatory elements (*31*).

In total, we identified 58 and 50 pREs that regulated either migration or proliferation, respectively, and 7 regions regulating both phenotypes (**Fig. 4D, Tables S8-9**, **Methods**). Although ECM stiffness is known to influence both cell growth and migration (*62*, *63*), we found no correlation between phenotype scores across pREs regulating either or both phenotypes (**Fig. S9A**). Furthermore, perturbations of pREs that regulated only migration or both phenotypes, had greater effects on migration compared to pREs that regulated only growth. There was no difference between the same groups for the growth phenotype (**Fig. S9B**). We examined the genomic contexts of two strong hit pREs: 1) an intronic pRE located within the gene *SKP2* that was identified in the growth screen (pRE #32) (**Fig. 4C,E**), and 2) an intergenic pRE located near the gene *CYR61* that was identified in the migration screen (pRE #62) (**Fig. 4B,E**). Both *SKP2* and *CYR61* were more highly expressed on 50kPa versus 1kPa surfaces (**Fig. 1B-C**). Upregulation of *SKP2* has been linked to metastasis and inhibition of *SKP2* can suppress cancer cell proliferation (*64*) and previous work identified the region near pRE #62 as a *CYR61* enhancer that is activated in colorectal cancer development (*65*). Our work further links the mechanical microenvironment as a potential factor in disease progression.

We next asked whether any of the pREs from the growth and migration screens were likely to be functional in other cell and tissue types. We quantified the overlap of the pREs with accessible chromatin regions in 95 of the ENCODE biosamples (**Table S10, Supplementary Text 2**). We found these pREs generally clustered across cell types into three main groups: 1) pREs that overlap accessible regions in all or most biosamples (“ubiquitous”), 2) pREs that overlap accessible regions in a majority of biosamples (“prevalent”), and 3) pREs that overlap accessible regions only in biosamples of similar cell or tissue types (“lineage-specific”) (**Fig. S10**). pREs had the greatest overlap with accessible chromatin regions for highly adherent cell types (e.g., endothelial and fibroblast lineages) and the least overlap with suspension cell types (e.g., T-cells and K562 cells), in accordance with our observation that the H3K27ac signal of the *MYH9* mechanoenhancer increased as the adherence of the cell type increased (**Fig. 2F**).

### Single cell CRISPRi screening identifies gene targets regulated by mechanoenhancers that drive cellular growth and migration

To identify the gene targets that were regulated by the pREs identified in the proliferation and migration screens, we made a sub-library of gRNAs targeting 87 of the pREs (10 gRNAs/pRE) with the largest effect sizes in either or both of the previous screens (**Methods**). This library also included positive controls of promoter-targeting and known enhancer-targeting gRNAs, and 100 non-targeting negative control gRNAs, for a total of 1,005 gRNAs in the sub-library (**Table S11**). We transduced HFF cells expressing dCas9^KRAB^ cultured on TCP with this gRNA library at 0.33 MOI, and eight days post-transduction we profiled 103,440 quality single cell transcriptomes (**Fig. 5A**). We recovered a median of 1 gRNA per cell and identified an average of 159 cells containing each gRNA (**Fig. S11A-B**). To identify significant gene linkages to each pRE, we tested cells that had each gRNA versus all cells that did not receive that gRNA, and compared expression for all genes within +/- 1Mb from the targeted pRE (**Fig. 5B, Table S12, Methods**), as previous studies suggest that most cis-regulatory interactions occur within this distance (*30*, *66*–*68*).

**Fig. 5.**
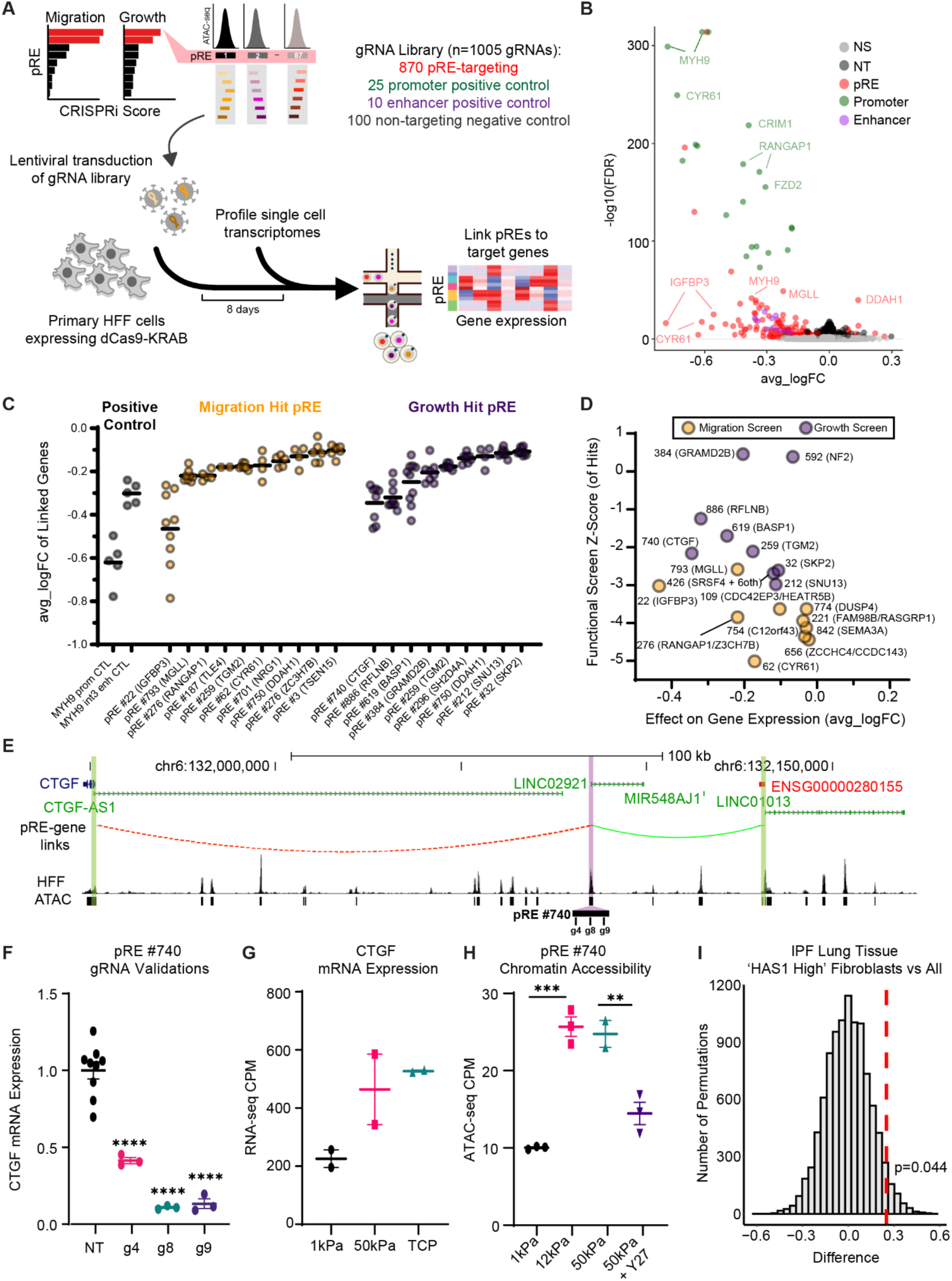
Single cell CRISPRi screen identifies genes regulated by mechanosensitive regulatory elements. **(A)** Overview of single cell CRISPRi screen workflow. Briefly, a gRNA library targeting pREs identified in the migration and growth bulk screens was delivered to CRISPRi HFF cells and single cell transcriptomes were profiled eight days later. **(B)** Volcano plot comparing the change in mRNA expression (avg_logFC) versus the significance (-log10(FDR)) of each gRNA-gene connection. Significant gRNA-gene connections for gRNAs targeting pREs (‘pRE’), positive control gRNAs targeting previously identified enhancers (‘Enhancer’), positive control gRNAs targeting promoter regions (‘Promoter’), and non-targeting gRNAs (‘NT’) are colored as red, purple, green, and black, respectively (FDR < 0.01). Nonsignificant (‘NS’) gRNA-gene connections are colored in light grey. **(C)** Average logFC of gene expression for MYH9 promoter-targeting positive control gRNAs and intron 3 enhancer-targeting positive control gRNA (grey), the ten migration (yellow) and growth (purple) pREs with greatest effects on gene expression following perturbation. Points shown are individual gRNA-gene linkages corresponding to each pRE, and all regions shown display significant reduction of the target gene (FDR < 0.01). **(D)** Z-scores of hit gRNA for each pRE from functional screening versus the average logFC of significant gRNA from the same pRE. Points shown for the top 10 pREs by Z-score from functional screening, along with the greatest absolute fold-change of pRE-gene linkages from scRNA-seq screen. **(E)** Browser track showing pRE-gene connections for a pRE proximal to *CTGF*, with red and green indicating decrease and increase in gene expression, respectively. Purple shading indicates pRE #740 and green shading indicates promoters of differentially expressed genes. ATAC-seq signal tracks and peaks in HFF cells are shown. **(F)** CTGF mRNA expression measured via RT-qPCR for individual gRNA validations (non-targeting (‘NT’), N=6; N=3 for all other gRNAs (‘g4’, ‘g8’, ‘g9’)). **(G)** CPM values for CTGF mRNA expression from bulk RNA-sequencing of HFF cells cultured on 1kPa (N=2), 50kPa (N=2), or TCP (N=2) surfaces. **(H)** CPM values for chromatin accessibility of CCN2 enhancer in HFF cells cultured on 10ka (N=3), 12kPa (N=3), or 50kPa (N=2), surfaces or cultured on 50kPa surface and treated with ROCKi (Y27) (N=3). **(F-H)** Individual points represent biological replicates. Error bars represent mean +/- 1 SEM. **** indicates p-value < 0.0001, *** indicates p-value < 0.001, ** indicates p-value < 0.01. **(I)** Distribution of the difference in effect size for pRE-connected genes versus permuted samples comparing ‘HAS1 High’ Fibroblasts versus all other cell types (‘All’) within IPF lung tissue. One-tailed p-value for permuted samples comparing the mean effect size is shown in the plot. Red dashed line indicates the observed difference between the effect size of the pRE-connected genes and all other genes.

In total, we identified 201 significant pRE-gene connections and connected 65 pREs to at least one gene (74.7%, 65/87), linking a median of 2 genes to every significant pRE and 1 pRE to each gene with at least one connection (**Fig. S11C-E**, **Methods**). We recovered one pRE with >10 gene linkages that was accessible across multiple ENCODE biosamples (“ubiquitous” cluster, **Fig. S12**, **Supplementary Text 3**). For the positive controls, we recovered 92% of expected promoter-targeting and 100% of enhancer-targeting gRNA-gene connections (**Fig. S13A**) that resulted in significant decreases in target gene expression (**Fig. S13B-E**). Notably, perturbation to the *MYH9* intron 3 enhancer (**Fig. 2**) led to reduced *MYH9* expression and also increases in expression of two additional genes, *APOL2* (∼90kb downstream) and *RAC2* (∼900kb upstream). Across the identified pRE-gene linkages, the magnitude of gene expression change correlated positively with the basal gene expression level, showing that more highly expressed genes are more robustly repressed (**Fig. S14A**), and change in gene expression diminished as the distance between pRE and target gene increased (**Fig. S14B-D**), consistent with previous work (*30*).

Prediction methods for linking noncoding regulatory elements to the affected gene(s) often rely on genomic proximity (e.g., nearest gene) or chromatin conformation data. We first computed the number of genes located between the pRE and the target gene. In contrast to predictions that nominate the nearest gene as the target of a *cis*-regulatory element, we found a median of 3 and a mean of 6.5 genes were “skipped” by the pRE to regulate target gene expression (**Methods**). Moreover, 37.1% of pRE-gene links have at least one other gene in between the pRE and the regulated gene, with 21.8% of links skipping at least five genes (**Fig. S15A-B**). This result is consistent with previous work in which 33% of identified enhancer-gene pairs in K562 cells skipped at least one gene (*4*, *5*, *30*, *31*). We next assessed the overlap of the pRE-gene links with high-resolution Micro-C chromatin contact data (*69*) (**Methods**). We observed all targeted pREs and all genes within the tested window have at least one chromatin contact. Only 13.8% of pRE-gene links identified in this study have annotated Micro-C chromatin looping between the pRE and the promoter region of the paired gene. CRISPRi perturbation of the pairs with evidence of looping led to a significantly greater decrease in gene expression than those without looping and those pairs were also located closer in genomic space than those without looping (**Fig. S15C-D**). We observed the same trend at a relaxed pRE-gene link FDR threshold of 0.05 (**Fig. S15E-F**). These results demonstrate that enhancers often may not regulate the nearest gene and also provide functional evidence for potential sub-analysis threshold looping and/or looping-independent *cis*-regulatory interactions.

### Genes linked to mechanically regulated pREs play key roles in diverse cellular functions

Next we examined the downstream target genes for many of the pREs that showed strong functional significance in driving growth or migration during screening. We first assessed the positive control *MYH9* promoter and mechanoenhancer perturbations, and observed that targeting the mechanoenhancer led to ∼50% of the repression as targeting the *MYH9* promoter (**Fig. 5C**), in agreement with the previous singleton gRNA experiments (**Fig. 2G**). We next compared the effect size of the top eight pRE-gene linkages from the growth and migration screens (ranked by phenotype Z-score) to the impact on target gene expression in the single cell screen, with the top two pRE-gene linkages that influenced single cell gene expression for comparison (**Fig. 5D**). pRE#62, which had the strongest combination of effects on gene expression and migration, connected to *CYR61*. Similarly, pRE #740 perturbation had strong effects on both *CTGF* gene expression as well as cellular growth (**Fig. 5E**). Both CYR61 and CTGF are canonical YAP/TAZ target genes. *CYR61* has been strongly linked to migratory phenotypes across many cell-types (*70*), and CTGF is known to play key roles in cellular growth (*71*, *72*).

pREs that we identified as being strong drivers of cell migration were found to change the expression of genes known to function in mechanoresponsive and cell migration-related pathways **(Fig. 5C**). These include *CYR61* (*CCN1*, **Fig. S16**), *DUSP4* (**Fig. S17A-B**), *FAM98B and RASGRP1* (**Fig. S18**), and *RANGAP1* (**Fig. S19**) (**Supplementary Text 4**). Targeting growth pREs led to changes in expression of genes with known functions in cell proliferation, including *CTGF* (*CCN2*, **Fig. 5E**), *NF2* (**Fig. S16C,D**), and *SKP2* (**Fig. S20**) as well as a novel regulator of cell proliferation, *RFLNB* (**Fig. S17C-D**) (**Supplementary Text 5**).

To validate the pRE-gene linkages identified in the scRNA-seq CRISPRi screen, we selected 13 pRE-gene connections for singleton gRNA validations and validated 30 gRNAs across these connections. During validation, we delivered individual gRNAs to the same HFF CRISPRi cell line cultured on TCP and assayed for changes in gene expression via RT-qPCR at eight days post-transduction. Through these validations we confirmed 10/13 pRE-gene connections and identified mechanoenhancers of *CTGF*, *RFLNB*, *SKP2, NF2* from the growth screen and *CYR61*, *DUSP4*, *FAM98B*, *RASGRP1*, *RANGAP1*, and *ZC3H7B* from the migration screen (**Fig. 5C-D, Fig. S16-20, Table S13**). The changes in mRNA expression in the validations were significantly correlated with the gene expression changes in the single cell screen (**Fig. S21**). Culturing HFF cells on increasingly stiffer substrates led to increases in expression of the pRE-linked genes, *CTGF*, *SKP2*, and *RANGAP1*, and treatment of HFF cells with the ROCKi Y-27632, for only one hour, led to changes in chromatin accessibility of the *CTGF*, *SKP2*, and *RANGAP1* mechanoenhancers demonstrating their plasticity and responsiveness to changes in intracellular actomyosin contractility (**Fig. 5E-H**, **Fig. S19D-E, Fig. S20B-C**). Collectively, these data demonstrate that many of the pREs identified in the screen are bona fide mechanoenhancers that regulate the expression of key genes and ultimately alter cell growth and migration in response to ECM stiffness cues.

### Mechanoenhancers regulate genes that play key roles in fibrosis

Since cellular responses to increased tissue stiffness can lead to both tumor growth and fibrosis (*73*), we explored whether the pRE-connected genes functioned in related disease processes. Using the union set of all genes connected to at least one pRE in the single cell screen, we performed gene set over-representation analysis and observed significant overlap with cancer- and fibrosis-related molecular signatures, biological pathways, and TF binding at promoter regions (**Fig. S22A-B, Table S14**). In idiopathic pulmonary fibrosis (IPF), fibroblast populations are characterized by uncontrolled proliferation and survival, and can remodel the ECM to a stiffer pro-fibrotic environment (*74*, *75*). Therefore, we sought to determine if the stiffness-activated pREs may contribute to tissue responses or transformations in IPF. We compared the accessibility of the pREs in diseased IPF versus healthy lung tissue (*76*). We found that accessible chromatin regions that overlapped hit pREs identified in the bulk phenotype screens and pREs connected to a gene in the single cell CRISPRi screen were significantly more accessible in IPF lung tissue versus lung tissue from unaffected controls (**Fig. S23A-B**). Given this difference in chromatin accessibility, we hypothesized that the genes regulated by the pREs would be upregulated in disease-associated cells. We compared the expression profiles of pRE-connected genes in lung tissue from individuals with IPF using a previously published single cell gene expression dataset (*77*) (**Methods**). Compared to random permutations, we observed that the pRE-connected genes were more highly expressed in the pathogenic fibroblast population (‘HAS1 High’) compared to all other cell types (**Fig. 5I**). These results further support a role for mechanoresponsive pREs regulating gene expression in pathogenic cell types in the context of a disease with a dysregulated mechanical microenvironment.

## Conclusions

Using complementary genome-wide epigenetic profiling, epigenetic editing, phenotypic screening, and single cell CRISPRi screening methods, we identified a class of *cis*-regulatory elements that are responsive to changes in material stiffness, and refer to these elements as ‘mechanoenhancers’ for simplicity. Mechanoenhancers could be more active on either soft or stiff substrates. Detection of stiff material cues by cells requires intracellular contractility, and accordingly we found that blocking cell contractility via ROCKi was sufficient to substantially reduce chromatin accessibility of ECM stiffness-sensitive peaks. Functional screening revealed that some pREs could act as key transcriptional drivers influencing a number of fundamental cell processes, including ECM mechanosensing, apoptosis, cellular growth, and migration.

Using single cell CRISPRi screening, we linked downstream target genes for 75% of the pREs identified as hits in the functional growth and migration screens. These mechanoenhancer-gene connections revealed that, unlike promoter-based regulation, mechanoenhancers often regulated multiple downstream gene targets (with a median of 2 recovered linkages per pRE) and regulated transcription across large genomic distances. Epigenetic repression of stiffness-activated mechanoenhancers resulted in marked reductions in target gene expression (with a range of ∼15-90% repression and a median of ∼50% repression), even in the continued presence of strong mechanical stimuli such as rigid tissue culture plastic. This suggests that epigenome editing of mechanoenhancers can be an effective means of decoupling mechanically-driven behaviors from the mechanical stimuli.

On stiff ECM conditions, we found that the activities of multiple canonical mechanosensitive signaling pathways likely combine to play a large role in driving mechanoenhancer activity. In peaks with increased chromatin accessibility on stiff materials, we found enrichment in motifs for many previously identified mechanosensitive transcription factors (including TEAD, AP-1, SP-1, and SRF/CaRG) in addition to motifs for signaling pathways that have not previously been associated with mechanosensing pathways (BORIS, E2F3, MED1)(*27*, *36*)(*37*, *78*). Through unbiased functional screening, we further identified key mechanoenhancers driving downstream gene targets of these mechanosensitive pathways, including mechanoenhancers for the canonical YAP/TAZ target genes *CTGF* (growth screen) and *CYR61* (migration screen)(*79*). A key question for future work will be determining how these mechanosensitive pathways work in concert to regulate changes in gene expression. To this end, we noted that the nuclease-active Cas9 editing screen of the MYH9 mechanoenhancer revealed that both a SRF/CaRG motif as well as a HLTF motif contributed strongly to enhancer function, suggesting that mechanoenhancers may be key integrators of mechanosensitive signaling pathways.

Previous studies examining chromatin accessibility in soft hydrogel conditions with ATAC-seq did not find peaks with increased accessibility on soft materials(*37*, *39*). However, through the use of on-plate ATAC-seq processing that did not require de-adhesion of cells prior to collection, we identified a subset of peaks that were more accessible on soft ECM. Peaks with significantly increased chromatin accessibility in soft ECM environments were ∼5-fold depleted in promoter regions as compared to the peaks found to be increasingly accessible on stiff ECM. De novo motif analysis of these peaks reveal that AP-1 family motifs were by far the most enriched motif (present in over 50% of the peaks), with lower-level enrichment for RUNX1, FOXF1, and CEBPE motifs. AP-1 motifs were also recovered in peaks more open on stiff ECM, though the motifs were markedly different with the stiff ECM AP-1 motif having higher levels of GC content. The lower GC content present in the soft ECM AP-1 motif as compared to the stiff ECM AP-1 motif supports stronger enhancer binding by AP-1 in the soft ECM accessible peaks. Partial assembly of transcription factors or co-activators at non-coding loci can function as an enhancer selector or priming for potential enhancer functionality (*80*). Further, this enhancer selection was driven by AP-1 binding. We note that peaks more accessible on soft ECM do not become further activated with increasing stiffness, suggesting that they could be primed to interact with other microenvironmental stimuli. Chemical and mechanical stimuli have previously been shown to interact in a combinatorial fashion to drive downstream behaviors including differentiation and transformation (*7*, *52*), and we postulate that this AP-1 priming in response to mechanical stimuli may play a key role in this functionality.

In contrast to the well-described mechanosignaling pathways that increase transcription on stiff ECM, there is less known about mechanosensitive signaling pathways that drive increased transcription on soft ECM and/or inhibit transcription on stiff ECM (*11*). Recent work has begun to describe mechanisms of mechanically-induced epigenetic repression. Dynamic mechanical stretch leads to deposition of H3K27me3 resulting in transcriptional repression (*21*–*23*). Furthermore, resistance to apoptosis induced by EGFRi/MEKi drug treatments in NSCLC cells is driven by YAP complexing with the SLUG (SNAI2) transcriptional repressor at genomic loci to drive epigenetic repression of *BMF(81)*. Specifically in NSCLC cells on tissue culture plastic, YAP and SLUG proteins were both complexed with DNA across intergenic and intronic peaks near *BMF* including the stiffness-repressed BMF intron 4 mechanoenhancer identified in our work. A key topic for future investigation will be to better understand which mechanosensitive signaling pathways may combine to promote epigenetic repression of mechanoenhancers on stiff ECM. We posit that one common mechanism of the enhanced activity of mechanoenhancers on soft ECMs could be the lack of stiffness-induced epigenetic repression.

One common characteristic of mechanically-induced disease states, including cancer, atherosclerosis, and fibrosis, is the feed-forward mechanical signaling driving these disease conditions (*82*–*84*). During feed-forward signaling in these disease contexts, small changes in ECM stiffness activate the expression of genes that serve to further reinforce these signaling changes (*85*, *86*). We identified mechanoenhancers that drive the activation of multiple genes that likely contribute feed-forward signaling due to their functionality, including *CTGF*, *CYR61*, *MYH9*, *RFLNB*, *RANGAP1*, *RASGRP1*, and *NF2*. One clear example of a single feed-forward loop is through the activity of the mechanoenhancer for *RANGAP1*, a key factor in cytoplasmic-nuclear shuttling that promotes increased import into the nucleus (*18*, *87*). In this feedforward pathway, mechanical force activates the mechanoenhancer resulting in increased *RANGAP1* expression, which potentially promotes nuclear import and further increases the sensitivity of that cell to subsequent mechanical signals. This is consistent with a proposed biophysical mechanism of mechanical force biasing nuclear transport, and extends that model to suggest that this process may be tuned by transcriptional feedback of key nuclear transport machinery like *RANGAP1 (88)*. Similarly, the *MYH9* mechanoenhancer drives increased expression of *MYH9* in response to ECM stiffness, thereby initiating a possible feed-forward mechanical loop (*89*) wherein enhanced contractility potentially further activates the mechanoenhancer and subsequent changes in gene transcription then serve to reinforce functional states.

We additionally identified evidence that these mechanoenhancers may be active in the idiopathic pulmonary fibrosis disease state. Specifically, we found that hit pREs from the functional screens were significantly more accessible in bulk IPF lung tissue compared to healthy tissue (*76*). We also found that target genes of stiffness-activated mechanoenhancers were significantly and specifically upregulated in the pathogenic ‘Has1’ fibroblast sub-population that contributes to increased collagen deposition in IPF (*75*). This further supports a mechanism by which mechanical cues from the diseased cellular microenvironment may activate mechanoenhancers that further potentiate both intracellular and extracellular feed-forward signaling that contributes to disease progression.

In this work we show that mechanoenhancers act as key downstream mediators of mechanosensitive signaling pathways and can function as strong drivers of cellular behavior. Epigenetic repression of these mechanoenhancers with CRISPRi allowed for the decoupling of these ECM-driven cellular behaviors from their mechanical stimuli. Therapeutic modulation of known gene targets implicated in mechanical disease states like cancer, atherosclerosis, and fibrosis has historically proven to be challenging, as the repression of these genes usually additionally removes the key homeostatic function of these genes. We find that mechanoenhancers regulate several genes known to be central players in these feedforward loops. Since epigenome editing of key mechanoenhancers can function to break the feed-forward signaling loops that dictate mechanical disease states while leaving homeostatic function intact, they may provide novel therapeutic targets in mechanosensitive diseases. Moving forward, epigenome editing of mechanoenhancers will be a powerful tool towards the precise engineering of the cellular response to the mechanical microenvironment, and could have widespread applications in both cell engineering and gene therapy.

## Acknowledgments

We would like to thank the Duke Sequencing Core and the Duke Viral Vector Core for their help and guidance, and members of the Gersbach and Crawford Labs at Duke University who have provided feedback throughout this project. We also thank Purushothama Rao Tata, PhD, Zachary Farino, PhD, for their suggestions. Fig. panels containing images of sequencing machines were created with BioRender.com. Funding was provided by National Institutes of Health grants UM1HG009428, UM1HG012053, U01AI146356, RM1HG011123, National Science Foundation grant EFMA-1830957, an Allen Distinguished Investigator Award from the Paul G. Allen Frontiers Group to CAG, Open Philanthropy, National Science Foundation Graduate Research Fellowships to LRB and CKT (NSF-GRFP DGE - 2139754), and a Regeneration Next Postdoctoral Fellowship to BDC.

## Data and materials availability

Public datasets used in this study are provided in Table S19. All datasets generated in this study will be made available prior to publication under GSE243765. All code used in this manuscript will be made available in an online repository prior to publication. Addgene catalog numbers for plasmids used in this study are noted in the methods. All materials used in this study will be made available upon request.

