## Supplementary Materials for "Mechanosensitive genomic enhancers potentiate the cellular response to matrix stiffness"

**The PDF file includes:**

Materials and Methods
Supplementary Text
Figs. S1 to S24

### Materials and Methods

Cell culture

Primary human neonatal fibroblasts (HFF cells) were acquired from ATCC (CRL-2097) and cultured in DMEM with 10% FBS, 1% AntiAnti, and 1% NEAA (Sigma) on TCP. All work was performed within 30 doublings from the initial passage of the vial.

RNA-Seq and Omni-ATAC-Seq

*Cell culture and soft hydrogel processing*

Polyacrylamide hydrogel 35 and 150 mm PetriSoft EasyCoat dishes (Matrigen) with an Elastic Modulus of 1 kPa ("soft") and 50 kPa ("stiff") were used for all NGS experiments. These dishes were incubated for 5 minutes with sterile PBS, rinsed two more times with sterile PBS, followed by addition of 10 ug/mL fibronectin (Sigma) for 30 minutes at room temperature. Fibronectin was then removed and dishes were rinsed twice with sterile PBS, followed by a 20 minutes incubation with complete growth media while cells were passaged. Media was removed from the dishes and cell suspensions were added and allowed to attach overnight.

*Bulk RNA-Seq*

40k HFF cells were seeded on 50 kPa dishes and TCP dishes, while 70k HFF cells were seeded on 1 kPa dishes to achieve the same effective plating density due to slightly reduced HFF attachment rates (and spreading) on 1kPa hydrogels. 20 hours after seeding, cells were trypsinized spun down at 300g for 5 minutes and then RNA was isolated from the cells using the Norgen Total RNA Purification Kit (#17250) according to the manufacturer's protocol, and samples were run on a RNA TapeStation (Agilent) to verify all samples had a RIN score > 8. cDNA Libraries were built from our RNA inputs using the TruSeq Stranded Library Prep Kit (Illumina #RS-122-2101) according to manufacturer's instructions. Quality control was performed by running the amplified libraries out on a High Sensitivity D1000 Tapestation (Agilent) to confirm expected size, and Qubit dsDNA HS assays were performed to determine a final concentration. Libraries were diluted to 10nM and pooled together in equal volumes, followed by sequencing performed on an Illumina HiSeq 2500 using a 50bp PE RapidRun kit. Resulting reads were subjected to adapter trimming using Trimmomatic v0.32 (*90*), aligned to GRCh38 with the STAR v2.4 aligner (*91*), and counts were retrieved using featureCounts (*92*) from subread version 1.4.6p4 with Gencode v22 gene annotations used as reference. Differential expression analysis was performed using edgeR quasi-likelihood methodology (*93*) and data was visualized using Degust (*94*) and Rstudio. Genes with significant differential expression were determined using a threshold of FDR < 0.05 and absolute value of Log2(FC) > 1.

*Omni ATAC-seq*

Cells were seeded on Matrigen dishes of varying stiffness (1, 12, 50 kPa elastic modulus) at slightly variable densities to account for reduced HFF attachment on softer substrates (70k, 45k, 40k HFF cells seeded per group respectively) and allowed to culture for 20 hours overnight. For Y-27632 ROCKi experiments, the cells were seeded as normal, but 1 hour prior to harvest 10uM Y-27632 ROCKi (StemCell Tech) in growth media was added to the cells. The Omni-ATAC-seq protocol was used to minimize mitochondrial reads from the preps (*95*), however no trypsinization was used and instead on-plate disruption/removal of nuclei (using the digitonin present in the lysis buffer) was used to better preserve nuclear mechanical context and connectivity prior to transposition by the Tn5. Following the final PCR, libraries were cleaned with a 0.5x/1.8x double-sided SPRI clean. Libraries were subjected to quality control by determining the number of cycles required to reach 25% of the peak threshold in the diagnostic PCR, as well as running the amplified libraries out on a High Sensitivity D1000 Tapestation (Agilent) to confirm expected size, and Qubit dsDNA HS assays were performed to determine a final concentration. Libraries were individually diluted to 6nM and then pooled at equal volumes prior to sequencing on an Illumina HiSeq 4000 using a single lane of 50bp single end reads. FastQC (*96*) was used to identify read quality, and adapter reads were trimmed using Trimmomatic v0.32 (*90*) followed by Bowtie (*97*) alignment (v1.0) of the reads to the reference genome using the settings: -v 2 –best –strata -m 1 with duplicate reads removed using Picard MarkDuplicates (v1.13) and ENCODE hg38 blacklist reads removed using bedtools2 v2.25 (*98*). Peak calling was performed using MACS2 with narrowPeak settings and a threshold of FDR < 0.001 (*99*), and a master peak set was generated as the union set of all called peaks across every sample analyzed (224,906 unique regions total). Count matrices were made using featureCounts (*92*) and DEseq2 v1.36 was used for differential accessibility analysis (*100*). Annotation of genomic regions was performed using ChIPSeeker (*101*), interactive visualization of processed data was done using Degust (*94*) and Rstudio along with ggplot2 and tidyverse plugins were used to generate data visualizations. Sequencing-depth normalized ATAC bigWig files were generated using deeptools bamCoverage v3.0.1 (*102*). All motif analysis was performed using the HOMER suite (*103*).

<u>MYH9 Locus Screening</u>

*Library design and cloning*

Using the ATAC-seq data, every open chromatin region that was within 440 kb of the MYH9 TSS was used as input to generate an oligo pool. For each ATAC-seq peak, we included any gRNA that had a GuideScan specificity score of > 0.2, which has previously been shown to increase the quality of non-coding screens (*104*). This resulted in 114 peaks represented in the library, with an average of ~41 gRNA/peak. We also included 500 non-targeting gRNA as negative controls (*105*). This combined gRNA library of 5,192 gRNA was synthesized as an oligo pool by Twist Biosciences with common overhangs for cloning into our lentiviral backbone.

This oligo pool was PCR amplified, and a hU6-driven lentiviral gRNA vector (pBDC119) was then digested with Esp3I, gel purified, and then ligated along with the amplified oligo pool by Gibson assembly. Following a 1x SPRI cleaning, the Gibson assembly was transformed into Endura competent cells (Lucigen) according to the manufacturer's protocol, and cultured overnight before maxi-prepping the gRNA-library plasmid. A PCR amplicon across the gRNA region of the resulting plasmid was sequenced to a depth of ~100k-1M read pairs on an Illumina miSeq in order to verify coverage across the entire gRNA library (**Fig. S24**).

*Lentiviral generation and functional titering of MYH9 locus library*

gRNA library plasmid was co-transfected into ~18M HEK293T cells along with two lentiviral packaging plasmids using Lipofectamine 3000 (ThermoFisher). 20 hours post-transfection, the transfection media was removed and fresh growth media was added. Media containing viral particles was removed one day later at 48 hours post-transfection and stored, replaced with fresh media and collected one day later before being stored at 4C. Combined media containing viral particles was filtered through 0.45 μm low-protein binding filters, and then concentrated using Lenti-X Concentrator (Takara Bio) according to the manufacturer's protocol. Functional titering to determine MOI was performed by transducing HFF cells across a 50x-10,000x dilution range of the viral stock, and then subjecting the cells to FACS-based cell sorting to identify what percent of the population was mCherry+ for each viral stock dilution.

*CRISPRi locus screen*

A stable HFF line was created using a lentiviral dCas9-KRAB construct (pLV-hUbC-dCas9-KRAB-2A-Blast (pJB289)), followed by the gRNA library being transduced at an MOI of ~0.33 and Puro selection for four days at 1 ug/mL. Cells were maintained for an additional four days, prior to trypsinization and fixation at Day 10 post-transduction. Following trypsinization with 0.25% Trypsin-EDTA for 5 minutes at 37C, trypsin was neutralized with 1X volumes of complete growth media following by 300g for 5 minutes centrifugation and aspiration of the supernatant, one rinse with 1X volume PBS followed by another centrifugation and aspiration leaving 200uL of PBS above the pellet. The eBioScience ICC Fixation kit (ThermoFisher) was used to fix/permeabilize cells according to manufacturer's instructions, with both reagents being equilibrated to room temp prior to usage. Fixation was performed through the addition of 500uL eBioSciences Fix/Perm Buffer (ThermoFisher) to the 200uL PBS and pellet, and incubation at room temperature for 20 minutes. At the end of this incubation 1X Permeabilization Buffer was added to 8mL total volume, spun at 600g for 5 minutes, followed by an additional perm buffer rinse. Following this step: HFF cells were counted, and ~2M cells were removed to be used for unsorted controls, and ~500k cells were set aside to be control samples for single channel compensation controls. Immunostaining of MYH9 was performed using a AlexaFluor-488 conjugated Rabbit monoclonal anti-NMMIIA antibody (clone EPR8965, Abcam, #ab204675) at a ratio of 0.5 uL antibody per 300k HFF cells per 100uL of Perm Buffer which was determined to be the ideal staining ratio using an antibody titration series. HFF cells were incubated for 30 minutes at room temperature in the dark on a nutating rocker, a 600g for 5 minutes spin, and two repeats of 3mL 1X Perm Buffer rinse/spin cycles. Following the last spin down, cells were resuspended in FACS Buffer [1X PBS supplemented w/ 1% BSA (Sigma) and 0.5mM EDTA (Sigma)] at density of ~9M cells/mL and sorted. A SH800 Cell Sorter (Sony Biotechnologies) was used to separate out the top/bottom-expressing MYH9 fractions following immunostaining. Compensation panels were set up using single channel expressing cell populations including untreated cells, antibody-only cells, mCherry-only cells. The top 10% and lower 10% of the MYH9 population was sorted off and used for downstream gRNA-enrichment analysis and sequencing.

*gDNA recovery and library preparation*

Cells were counted following sorting to verify enrichment, followed by DNA recovery/extraction from fixed cells using the PicoPure DNA extraction kit (ThermoFisher) according to manufacturer's instructions. Recovery digests were performed for 20 hours at 65C using up to 1.5M HFF cells per reaction volume. All gDNA was split between sample-indexed 100uL Q5 PCR reactions (up to ~340ng max input per 100uL reaction) to amplify out the gRNA protospacer from HFF cells. These PCRs from gDNA were run as follows [ 98C for 30s / 25x: 98C for 10s, 60C for 30s, 72C for 15s / 72C for 2 min] with primers in **Table S15**, followed by individual PCRs being pooled together and subjected to a double-sided 0.65X/1X SPRI clean-up. Quality control was performed by running the amplified libraries out on a High Sensitivity D1000 Tapestation (Agilent) to confirm expected size, and Qubit dsDNA HS assays were performed to determine a final concentration. All libraries were pooled to an effective concentration of 4 nM and combined in equal volumes prior to sequencing on an Illumina MiSeq, using a v2 50 cycle reagent kit with Read1 being 21 cycles (protospacer) and index read 1 being 6 reads (sample barcoding).

*MYH9 locus library analysis*

Resulting FASTQ files were aligned to a custom reference sequence corresponding to the given gRNA library using bowtie2 and all downstream analyses were performed in R. All gRNA were verified to be represented in the baseline untreated library dat Day 8 post-transduction, and counts+1 for each gRNA were taken (to normalize for samples that dropped out in one condition) and normalized by sequencing depth for each library before downstream analysis (in counts per million reads sequenced, 'CPM'). Due to the highly-apparent strand bias in the positive-strand when targeting the MYH9 gene body (**Supplementary Text 1**) , we only included the non-interfering gRNA from the negative strand (2,863 gRNA). A ratio was taken of the CPM for each gRNA of the low MYH9 expression group to high MYH9 expression group to identify whether the gRNA perturbation led to increases in enrichment in either expression bin. Next, for each screen replicate the Z-score was calculated for each gRNA relative to the control non-targeting gRNA population using similar methodologies as previously described in (*6*). First, each sample's ratio was converted to a log2 fold-enrichment, and population statistics for the negative control non-targeting gRNAs (median, standard deviation, gRNA number) were calculated. For each individual gRNA, the median of the negative control fold-enrichment was subtracted from each individual gRNA's log2 fold-enrichment value, and this value was further divided by the standard deviation of the negative control non-targeting gRNA population to get an individual Z-score relative to the negative control population. Raw Z-score values from both replicates were pooled to calculate pRE-wide effects. Phenotype scores (t-score based) were calculated as:

$$Phenotype\ Score\ =\ U\ (pRE)\ -\ U\ CTL\ /\ \sqrt{(\frac{S\ var}{Nexp}+\frac{S\ var}{NCTL})}$$

$$Svar\ =\ Var\ (pRE)\ *\ (N(pRE)-1)\ +\ Var(CTL)*(Nctl\ -1)$$

*Individual gRNA validations*

For gRNA validations of all 5 hit pRE across the MYH9 locus (including the two promoter/exon1 regions). Oligos containing protospacer sequences were synthesized by IDT and cloned into pLV_hU6-sgRNA_hUbC-GFP-P2A-PuroR (Addgene plasmid #162335). Sanger sequencing was used to confirm the identity of the gRNA. Lentivirus was generated as previously described (ref method section). dCas9-KRAB expressing HFF cells were seeded onto TCP and transduced on day 0. 24 hours post-transduction, lentivirus was removed and replaced with fresh growth media. Puromycin selection was applied as described for the bulk screen, and cells were harvested nine days post-transduction. mRNA was isolated using the Norgen Total RNA Purification Kit (#17250) according to the manufacturer's protocol. 100 ng mRNA was used as input for cDNA amplification using the Invitrogen™ SuperScript™ VILO™ cDNA Synthesis Kit. For RT-qPCR, each reaction contained 1 uL cDNA, 7 uL H20, 1 uL Taqman probe for TBP, 1 uL Taqman probe for MYH9, and 10 uL Quantabio PerfeCTa FastMix II. Delta delta Ct analysis was performed in Microsoft Excel. Graphpad Prism was utilized to conduct one-way ANOVA tests followed by Tukey's HSD for post-hoc testing. Significance is reported in Fig.s as follows: *p-value < 0.05, **p-value < 0.01, ***p-value < 0.001. Taqman probe information provided in **Table S17**. A portion of the transduced cells for the MYH9-intron 3 pRE were propagated to day 15 and then subjected to MYH9 immunostaining and flow cytometry as described in the bulk screen section, with gain values held constant across all collections across samples. Populations were plotted to show shifts relative to transduction with the non-targeting gRNA. Noting high values of MYH9 promoter-targeting gRNA, we did a similar transduction and examined RNA expression at day 6 post-transduction and saw markedly lower levels of MYH9 expression, supporting the idea that MYH9 deficiencies in cytokinesis led to a dropout of transduced cells over longer timeframes (**Fig. S7**).

*Actin & vinculin labeling/immunostaining and focal adhesion imaging/analysis*

HFF cells were seeded into 24 well-plates while being transduced with lentiviruses encoding an all-in-one construct that expressed dCas9-KRAB/hU6-gRNA (Addgene plasmid #71236) with the gRNA being either a non-targeting control, an gRNA for the MYH9 intron 3 enhancer, and a gRNA for the MYH9 promoter. Viral media was removed 20 hours later, and replaced with complete growth media. Puromycin selection was started 2 days post-transduction, wherein 1.5 ug/mL Puromycin was added to the growth media for 3 days prior to removal of the antibiotic selection and continued passaging of the cells for expansion. Six days post transduction our transduced HFF cells were seeded at at ~5k cells/well into µ-Slide 8 Well Glass Bottom (ibidi) chamberslides that were coated with 10 ug/mL fibronectin for 45 minutes at room temp and rinsed 1x with PBS prior to seeding. Following an overnight culture, the media was removed on the chamberslide and 200uL of 4% PFA was gently added to each well and cells were fixed at room temp for 15 minutes, rinsed 2x with PBS, and then permeabilized with a permeabilizing solution [PBS supplemented with 0.5% TritonX-100, 10% w/v sucrose, 600uM MgCl2] for 10 minutes at 4C. Permeabilizing solution was then removed from cells, followed by 2x PBs rinses, and blocked with a labeling solution [1% bovine serum albuminutes (Sigma) in PBS] for 30 minutes at room temperature. Fresh labeling solution was added that contained a 1:300 dilution of a Rabbit monoclonal anti-Vinculin antibody (clone EPR8185, Abcam, #ab129002) and incubated in a nutating rocker in the dark overnight at 4C. The next morning the primary antibody was removed, rinsed 2x with labeling solution, and then a secondary solution that contained a 1:200 dilution of AlexaFluor488 Goat anti-Rabbit secondary (Thermo #A-11008), a 1:100 dilution of AlexaFluor647-Phalloidin (Thermo #A22287) and a 1:5000 dilution of DAPI was added for 1 hour at room temperature on a nutating rocker in the dark. Following three PBS rinses, chamber-slide wells were mounted with Vectashield Antifade Mounting Media (Vector Laboratories, H-1000-10). All focal adhesion and actin imaging was performed using a 20x/0.8NA objective on a Zeiss AxioOversver 7 and a quad-bandpass filter. Focal adhesion morphometric characteristics were quantified using vinculin images input to an online web tool, the Focal Adhesion Analysis Server (FAAS) (*106*). For this analysis the minimum adhesion size was set to 0.21 $\mu m^2$ and the stdev_thresh was set to 5.5. Each value is reported as the average across an individual cell within the group, with N=39-45 cells/per group for either the control non-targeting gRNA or the MYH9 intron 3 targeting gRNA.

<u>MYH9 intron 3 saturation mutagenesis screening</u>

*Library design and cloning*

For the MYH9 intron 3 pRE saturation mutagenesis library, we included any gRNA that was within the hit pRE from the MYH9 locus library, which resulted in 64 gRNA across the library. We also included 25 non-targeting gRNA (*105*) and 11 safe-targeting gRNA (*107*) as negative controls. This combined gRNA library of 100 gRNA was synthesized as an oligo pool by Twist Biosciences with common overhangs for cloning into our lentiviral backbone. This oligo pool was PCR amplified, and pLV_hU6-sgRNA_hUbC-GFP-P2A-PuroR (Addgene plasmid #162335) was then digested with Esp3I, gel purified, and then ligated along with the amplified oligo pool by Gibson assembly. Following a 1x SPRI cleaning, the Gibson assembly was transformed into Endura competent cells (Lucigen) according to the manufacturer's protocol, and cultured overnight before maxi-prepping the gRNA-library plasmid. A PCR amplicon across the gRNA region of the resulting plasmid was sequenced to a depth of ~100k-1M read pairs on an Illumina miSeq in order to verify coverage across the entire gRNA library *(***Fig. S24***)*.

*Lentiviral generation and functional titering*

gRNA library plasmid pool was co-transfected into ~7.8M HEK293T cells along with two lentiviral packaging plasmids using Lipofectamine 3000 (ThermoFisher). 20 hours post-transfection, the transfection media was removed and fresh growth media was added. Media containing viral particles was removed one day later at 48 hours post-transfection and stored, replaced with fresh media and collected one day later before being stored at 4C. Combined media containing viral particles was filtered through 0.45 μm low-protein binding filters, and then concentrated using Lenti-X Concentrator (Takara Bio) according to the manufacturer's protocol. Functional titering to determine MOI was performed by transducing HFF cells across a 0.75x-100x dilution range of the viral stock, and then subjecting the cells to FACS-based cell sorting to identify what percent of the population was mCherry+ for each viral stock dilution.

*MYH9 saturation mutagenesis screen*

HFF cells were transduced with a lentiviral SpCas9 construct (FUGW-SpCas9-2A-HygroR (pVG54)), selected with 100ug/mL hygromycin for 4 days with hygromycin in order to make a stable line. Following four passages the cells were frozen and used for subsequent screening experiments and validations. 600k HFF cells were transduced with lentivirus encoding the MYH9 intron 3 saturation pool. For screening, the same protocol was used as described above for the MYH9 CRISPRi locus screen, with 8 days of culture time prior to fixation, MYH9 immunostaining, FACS for the top/bottom 10% of cells, PicoPure gDNA recovery, and gRNA PCR and processing for enrichment across the low and high MYH9 expression bins.

*Individual gRNA validations*

For gRNA validations of all 3 hit gRNA that had significantly altered MYH9 expression and a non-targeting control gRNA, oligos containing protospacer sequences were synthesized by IDT and cloned into pLV_hU6-sgRNA_hUbC-GFP-P2A-PuroR (Addgene plasmid #162335). Sanger sequencing was used to confirm the identity of the gRNA. Lentivirus was generated as previously described above. Cas9 expressing HFF cells were seeded onto TCP and transduced on day 0. 24 hours post-transduction, lentivirus was removed and replaced with fresh growth media, cells were grown for 8 days (with 4 days of 1.5ug/mL puromycin selection). And for harvest cells were split with 500k cells for gDNA harvested following FACS (as detailed below) and RNA was harvested from 500k cells using a Norgen Total RNA Purification Kit (#17250). qPCR for MYH9 expression was performed as described above for the MYH9 locus screen.

*gRNA validation indel enrichment across MYH9 expression bins*

Additionally 500k cells were processed similarly to the screen that included cell fixation, MYH9 immunostaining, FACS for the top/bottom 10% of cells, PicoPure gDNA recovery. A MYH9 intron 3 PCR was performed with an amplicon size of 666 bp. All gDNA was split between sample-indexed 100uL Q5 PCR reactions (up to ~340ng max input per 100uL reaction) to amplify out the gRNA protospacer from HFF cells. These PCRs from gDNA were run as follows [ 98C for 30s / 25x: 98C for 10s, 60C for 30s, 72C for 15s / 72C for 2 min] with primers in **Table S15**, followed by individual PCRs being pooled together and subjected to a double-sided 0.65X/1X SPRI clean-up. Quality control was performed by running the amplified libraries out on a High Sensitivity D1000 Tapestation (Agilent) to confirm expected size, and Qubit dsDNA HS assays were performed to determine a final concentration. All libraries were pooled to an effective concentration of 4 nM and combined in equal volumes prior to sequencing on an Illumina MiSeq, using a v2 50 cycle reagent kit with Read1 being 21 cycles (protospacer) and index read 1 being 6 reads (sample barcoding). FASTQ reads were run through Crispresso2 (*108*) and indel enrichment in the low MYH9 bin was used to examine any overlapping TF motifs on common indel sites.

BMF Enhancer Characterization

*Luciferase Enhancer Reporter Assays*

BMF regions with differential accessibility were identified, and primers were designed to amplify these regions from gDNA isolated from the HFF cell-lines. Briefly, 2x 25uL reactions were run wherein 30ng gDNA was input with 2x KAPA HiFi Hot Start MM and 0.75uL of 10uM PCR primers (**Table S17**) for either Region #1/2/3 with an annealing temp of 63C. Sequences were confirmed via Sanger sequencing. These enhancer fragments were then assembled into an improved STARR-seq enhancer luciferase reporter vector (*109*) via Gibson assembly and clones were sequences via Sanger sequencing to confirm the fragment addition. To perform the luciferase assay, 15k HFF cells were seeded per well into a 24 well-plate one day prior to transfection, and the day of transfection fresh media was added immediately prior to lipofection, with either DMSO only or 10 uM Y-27632 added. Lipofectamine LTX (2.25uL per well) was used to transfect 300ng of plasmid at a mass ratio of 90% experimental firefly luciferase plasmid to 10% Renilla luciferase pRL-CMV control plasmid (Promega) into cells. Cells were harvested 24 hours later and the Promega DualGlo Luciferase Assay was performed according to manufacturer's instructions, with luciferase activity read on a Promega GloMax Discover instrument (0.3s integration time). The average of four blank wells was then set as the background level and subtracted from all experimental values. Firefly luciferase values for each well were normalized to the Renilla luciferase values. Each experiment was further normalized to the performance of an empty luciferase reporter plasmid as baseline.

*Latrunculin A Induction experiment culture*

Oligos containing protospacer sequences were synthesized by IDT and cloned into an all-in-one lentiviral vector expressing dCas9-KRAB-P2A-PuroR from an hUbC promoter and a gRNA from an hU6 promoter (Addgene plasmid #71236). All gRNA were selected as (-) strand gRNA to minimize the strand-bias artifact (**Supplementary Text #1**). Sanger sequencing was used to confirm the identity of the gRNA. Lentivirus was generated as described above for MYH9. HFF cells were transduced and seeded per well in a 24 well-plate on day 0, by adding 25uL of 20x concentrated virus along with 5k cells and growth media. 24 hours post-transduction, lentivirus was removed. Antibiotic selection was applied for four days and cells were grown for eight days post-transduction. At 9 days post-transduction cells were trypsinized, and were re-seeded at 5k HFF cells/well in a 24 well-plate for RNA experiments or 20k HFF cells/well in a 12WP for Caspase 3/7 experiments. To model detachment, on 11 days post-transduction the media was replaced with growth media containing either DMSO or 0.5 uM Latrunculin A. Cells were harvested for RNA or CaspaseGlo 3/7 analysis one day following the addition of Latrunculin A.

*RNA expression and Caspase 3/7-Activity assays*

mRNA was isolated using the Norgen Total RNA Purification Kit (#17250) according to the manufacturer's protocol. 100 ng mRNA was used as input for cDNA amplification using the Invitrogen™ SuperScript™ VILO™ cDNA Synthesis Kit. For RT-qPCR, each reaction contained 1 uL cDNA, 7 uL H20, 1 uL Taqman probe for TBP, 1 uL Taqman probe for BMF, and 10 uL Quantabio PerfeCTa FastMix II. Delta delta Ct analysis was performed in Microsoft Excel. Graphpad Prism was utilized to conduct one-way ANOVA tests followed by Tukey's HSD for post-hoc testing. Significance is reported in Fig.s as follows: *p-value < 0.05, **p-value < 0.01, ***p-value < 0.001. Taqman probe information provided in **Table S16**. For Caspase-3/7 activity assays, HFF cells were subjected to the CaspaseGlo 3/7 Assay (Promega) and Cell TiterGlo Assay (Promega) according to manufacturer's instructions with luciferase values read out on a Promega GloMax Discover instrument (0.3s integration time). The average of two blank wells per assay was then set as the background level and subtracted from all experimental values. CaspaseGlo3/7 values per group were further normalized to cell counts per group determined from the CellTiterGlo data.

<u>Bulk growth and migration functional CRISPRi screens</u>

*Library design and cloning*

The top 1000 regions from the ATAC-seq data that were increasingly-accessible on the stiff 50kPa substrates as compared to the soft 1 kPa substrate were used as input to generate an oligo pool. For each peak, we included any gRNA that had a GuideScan specificity score of > 0.2, which has previously been shown to increase the quality of non-coding screens (*104*). This resulted in 969 peaks represented in the library, with an average of ~20 gRNA/peak. We also included 1000 non-targeting gRNA (*105*), and 249 promoter-targeting gRNA for 83 positive control genes that have previously been shown to be key modulators of transwell migration following RNAi screens (*57*), with 3 gRNA per gene taken from the Dolcetto library (*110*). This combined gRNA library of 21,458 gRNA was synthesized as an oligo pool by Twist Biosciences with common overhangs for cloning into our lentiviral backbone. This oligo pool was PCR amplified, pLV_hU6-sgRNA_hUbC-GFP-P2A-PuroR (Addgene plasmid #162335) was digested with Esp3I and gel purified, and then the oligo pool and digested vector were ligated by Gibson assembly. Following a 1x SPRI cleaning, the Gibson assembly was transformed into Endura competent cells (Lucigen) according to the manufacturer's protocol, and cultured overnight before maxi-prepping the gRNA-library plasmid. A PCR amplicon across the gRNA region of the resulting plasmid was sequenced to a depth of ~100k-1M read pairs on an Illumina miSeq in order to verify coverage across the entire gRNA library (**Fig. S24**).

*Lentiviral generation and functional titering*

Concentrated lentivirus was generated by the Duke Viral Vector Core from this plasmid pool. Functional titering to determine MOI was performed by transducing HFF cells across a 50x-10,000x dilution range of the viral stock, and then subjecting the cells to a qPCR-based titering protocol that has been previously described in detail (*111*).

*Migration/Growth pRE library screen*

To perform screening, 600k HFF cells were transduced with the lentiviral library virus at 10.8 MOI to achieve a coverage of ~279 cells per gRNA. 20 hours after transduction the viral media was removed and replaced with fresh media, and starting 48 hours after transduction HFF cells were selected with 1 ug/mL puromycin for 4 days. Puromycin selection media was then removed and HFF cells were grown out for two additional days until day 8. On day 8, ~11M cells were counted and split between migration and growth screens. Coverage of at least 279 cells/gRNA was maintained for each group throughout the entire experiment.

<u>Migration Screening</u>: On day 8, the bottoms of 8 um transwell inserts for 6WP were coated with 10 ug/mL fibronectin at room temp for 45 minutes and then rinsed 1x with PBS for 30 minutes before use. HFF cells were counted, placed into low serum conditions (0.2% FBS) and seeded at 240k cells per transwell insert across 18 inserts (~4.4M cells total). These inserts were placed into 10% serum and cells were allowed to migrate for 24 hours. Following this first day of migration, each side of the membrane was separately trypsinized and counted, where 27% of the initial cells were recovered as migratory cells (~1.2M cells) and non-migrated cells were recovered from the top of the insert. These migratory and non-migratory populations were re-seeded (separately by group) in the same way on new fibronectin-coated transwell inserts, with 4-5 inserts seeded at 240k cells/insert and allowed to migrate overnight. Following these two rounds of migration we trypsinized and collected the cells that either migrated twice or did not migrate twice (with a similar number of cells, 24%, being found to have migrated during this second round) and isolated gDNA using DNeasy kits (Qiagen).

<u>Growth Screening</u>: HFF cells were counted on day 8 post-transduction, and gDNA from 2M HFF cells were harvested as the "Day 0" reference population using a DNeasy Blood and Tissue Kit (Qiagen). Around 1M HFF cells were reseeded into 15 cm dishes for ongoing culture, and then serially-passaged as normal for 14 doublings (either 21 days post-"Day0" for replicate 1 or 22 days post-"Day0" for replicate 2) while maintaining at least 1M cells per dish during each passaging, prior to the final gDNA harvest using a DNeasy Blood and Tissue Kit (Qiagen).

*Library preparation and sequencing*

All gDNA was split between sample-indexed 100uL Q5 PCR reactions (up to ~340ng max input per 100uL reaction) to amplify out the gRNA protospacer from HFF cells. These PCRs from gDNA were run as follows [ 98C for 30s / 25x: 98C for 10s, 60C for 30s, 72C for 15s / 72C for 2 min] with primers in **Table S15**, followed by individual PCRs being pooled together and subjected to a double-sided 0.65X/1X SPRI clean-up. Quality control was performed by running the amplified libraries out on a High Sensitivity D1000 Tapestation (Agilent) to confirm expected size, and Qubit dsDNA HS assays were performed to determine a final concentration. All libraries were pooled to an effective concentration of 4 nM and combined in equal volumes prior to sequencing on an Illumina MiSeq, using a v2 50 cycle reagent kit with Read1 being 21 cycles (protospacer) and index read 1 being 6 reads (sample barcoding).

*Screen analysis*

Resulting FASTQ files were aligned to a custom reference sequence corresponding to the given gRNA library using bowtie2 and all downstream analyses were performed in R. All gRNA were verified to be represented in the baseline untreated library at day 8 post-transduction, and counts+1 for each gRNA were taken (to normalize for samples that dropped out in one condition) and normalized by sequencing depth for each library before downstream analysis (in counts per million reads sequenced, 'CPM'). For migration screens: A ratio was taken of the CPM for each gRNA of the 2x migrated group to the 2x non-migrated group to identify migratory or non-migratory enrichment. For growth screens: A ratio was taken of the CPM of the Day 0 population relative to the final Day 21/22 population for each replicate. Next, for each screen replicate the Z-score was calculated for each gRNA relative to the control non-targeting gRNA population using similar methodologies as previously described(*6*). First, each sample's ratio was converted to a log2 fold-enrichment, and population statistics for the negative control non-targeting gRNAs (median, standard deviation, gRNA number) were calculated. For each individual gRNA, the median of the negative control fold-enrichment was subtracted from each individual gRNA's log2 fold-enrichment value, and this value was further divided by the standard deviation of the negative control non-targeting gRNA population to get an individual Z-score relative to the negative control population. Raw Z-score values from both replicates were pooled to calculate pRE-level effects. An individual gRNA was called as a "hit" if the Z-score was above 2 or below -2. pRE-level stats were generated by performing a Fisher's exact text relative to the non-targeting gRNA population, and a pRE-level was labeled significant for follow-up if the pval was less than 0.1. To select pREs for validation in single cell RNA-seq, we further selected the pRE hits that had more than one gRNA as a "hit" and had at least 10 gRNA/DHS in order to enable higher-powered analysis of the downstream data.

*Comparison of phenotype scores between regions regulating growth, migration, or both phenotypes*

Each significant region was labeled for the phenotype it regulated (one of growth, migration, or both). The phenotype (pZ) scores were compared between the three groups using a One-way ANOVA test followed by Tukey's post-hoc tests using the aov and TukeyHSD functions in R.

*Chromatin accessibility of significant screen regions in IPF vs unaffected control tissue*
We obtained ATAC-seq peak calls from GSE180242 (*76*). We intersected the pREs that were significant in either bulk screen with the peak calls using bedtools intersect. We then performed a Student's t-test comparing the fold change in chromatin accessibility for all overlapping peaks in IPF lung tissue vs unaffected control lung tissue using the t.test function in R.

*Analysis of chromatin accessibility across ENCODE biosamples*
We obtained the union set of DNase peak calls across 95 ENCODE biosamples using the 'Table Browser' utility on the UCSC Genome Browser (downloaded February 2023; 'wgEncodeRegDnaseClustered'). The union DNase peak calls were intersected with all regions included in the bulk screen library using bedtools intersect. Next, each region significant in at least one of two screens was labeled as '1' or '0' if the region did or did not overlap an accessible region in at least one biosample, respectively. The region X biosample visualization was generated using the pheatmap package in R with the following parameters: scale = "none", cluster_cols = TRUE, cluster_rows = TRUE. To extract the clusters, we used the cutree_col function specifying h=8. We then compared the phenotype scores between each cluster by performing One-way ANOVA tests followed by Tukey's post-hoc tests using the aov and TukeyHSD functions in R.

To determine if significant screen regions were enriched or depleted from accessible regions in specific biosamples, we performed Fisher's exact tests separately for each biosample comparing the number of significant and nonsignificant screen regions that overlapped or did not overlap an accessible region using the fisher.test function in R.

<u>Single cell RNA-seq screen</u>

*gRNA library design and cloning*
Following hit identification from the combined migration and growth screens (as described above), a library was designed that included the top 10 gRNA by pZ value across either screen for the 87 hit pRE (870 gRNA total). 100 non-targeting control gRNA with similar sequence composition to the targeting gRNAs were included in the library, and 25 gRNA targeting the promoters of contractile genes including *MYH9*, *RANGAP1*, and *CRIM1* were included, as well as the top gRNA from the MYH9 intron 3 enhancer as positive controls. In total our library contained 1005 gRNA sequences, which were synthesized as an oligo pool by Twist Biosciences with common overhangs for cloning into our lentiviral backbone. This oligo pool was PCR amplified, and a hU6-driven lentiviral gRNA CROP-seq vector (pLRB104) was then digested with Esp3I, gel purified, and then ligated along with the amplified oligo pool by Gibson assembly. Following a 1x SPRI cleaning, the Gibson assembly was transformed into Endura competent cells (Lucigen) according to the manufacturer's protocol, and cultured overnight before maxi-prepping the gRNA-library plasmid. A PCR amplicon across the gRNA region of the resulting plasmid was sequenced to a depth of ~100k-1M read pairs on an Illumina miSeq in order to verify coverage across the entire gRNA library (**Fig. S24***)*.

*Lentiviral generation and functional titering*

gRNA library plasmid was co-transfected into ~18M HEK293T cells along with two lentiviral packaging plasmids using Lipofectamine 3000 (ThermoFisher). 20 hours post-transfection, the growth media was removed and fresh growth media was added. Media containing viral particles was removed at 48 hours, replaced, and removed at 72 hours post-lipofection before being stored at 4C. Combined media containing viral particles was filtered through 0.45 µm low-protein binding filters, and then concentrated using Lenti-X Concentrator (Takara Bio) according to the manufacturer's protocol. Functional titering to determine MOI was performed by transducing HFF cells across a 50x-10,000x dilution range of the viral stock, and then subjecting the cells to a qPCR-based titering protocol that has been previously described in detail (*111*).

*Single cell CRISPRi screen*

To perform screening, 775k HFF cells stably expressing dCas9-KRAB were transduced at 0.33MOI with the CROP-seq lentivirus to maintain a coverage of at least 150 cells/gRNA. Following 20 hours, viral media was removed and replaced with regular growth media, and 48 hours post-transduction the cells selected with puromycin (1.5 ug/mL) for 4 days. Following puromycin selection, HFF cells were maintained until day 8, at which point cells were trypsinized and 150k cells were moved on to library prep.

*Single cell RNA-seq library preparation*

Cells were washed 3x with PBS and then resuspended to a final concentration of 1000 cells/uL. Approximately 20,000 cells were loaded onto each channel of a 10X Genomics' 3' Gene Expression (GEX) v3.1 assay chip. Downstream processing was performed according to the manufacturer's protocol. To recover the protospacer sequences (gRNA libraries), a tri-nested PCR was performed separately for each GEX library using 10% of the purified cDNA as input to reaction 1 as previously described (*30*). Briefly, 4ng cDNA was input into a 50 uL reaction with KAPA HiFi and PCR primers prLRB470 and prLRB471 (**Table S18**). The reaction was amplified for 12 cycles and then purified using 25 uL of AMPure XP DNA beads and eluted in 25 uL H20. 1 uL of the purified sample was input into reaction 2 using PCR primers prLRB472 and prLRB473 (**Table S18**). The reaction was amplified for 14 cycles and purified as described above. 1 uL of the purified sample was input into reaction 3 using PCR primers prLRB473 and prLRB289-302 (**Table S18**), amplifying each sample with a unique i7 sequencing index. The reaction was amplified for 7 cycles, purified using 25 uL of AMPure XP DNA beads (Beckman Coulter #A63881), and eluted in 25 uL Buffer EB (Qiagen #19086). Quality control of final libraries was performed prior to sequencing using the Agilent 2200 TapeStation with High Sensitivity DNA 5000 reagents, Qubit High Sensitivity dsDNA reagents, and KAPA Library Quantification Kit for Illumina platforms.

*Sequencing*
GEX libraries were pooled and sequenced on a NovaSeq 6000 S4 flow cell using the parameters: 28x10x10x90. gRNA libraries were pooled and sequenced on a NovaSeq 6000 S1 flow cell using the parameters: 28x10x10x90.

*Data processing*
Cell Ranger: All data processing steps were performed using CellRanger v6.0.1 and the human reference genome ('refdata-gex-GRCh38-2020-A') was downloaded from 10X Genomics' software downloads webpage. Fastq files for each flow cell lane and sequencing run were generated from .bcl files using the cellranger mkfastq pipeline. The corresponding fastq files for each sample were then merged. The merged fastqs were then processed using the cellranger count pipeline with the number of expected cells specified (--expect-cells = 15000). The gene expression libraries were then aggregated using the cellranger aggr pipeline. The gRNA libraries were aligned to a custom bowtie index containing all protospacer sequences included in the pooled gRNA library and the UMI counts corresponding to each gRNA-cell pair were obtained.

Seurat: The gene expression and gRNA UMI count data was imported into Seurat v3.1. A gRNA was defined as 'observed in a cell' if the gRNA had at least 5 UMI counts and comprised at least 0.5% of the total gRNA UMI counts in that cell. We then calculated the total percent of mitochondrial reads per cell and filtered for quality cells as follows:

```
cells[["percent.mt"]] <- PercentageFeatureSet(cells, pattern = "^MT-")
cells <- subset(cells, subset = nCount_RNA > 10000 & percent.mt < 20)
```

*Differential expression analysis*
Using the gRNA-cell assignments, differential expression testing was performed using the MAST framework (*112*) within Seurat v3.1 (*113*), comparing cells in which a given gRNA was observed versus all other cells with at least one gRNA observed excluding the given gRNA and testing all genes within +/- 1Mb of the midpoint of the pRE in which the gRNA is located. Gene coordinates were obtained from the Ensembl Human Gene v104 reference file. P-values were then FDR-corrected on an individual gRNA-level for all tests. All genes within +/- 1Mb of any targeting gRNA were used as input features for NT gRNA tests (N=1,313). Significant gRNA-gene and corresponding pRE-gene pairs are defined as FDR < 0.01.

*Calculation of interaction distance (ep_length)*
The distance between the gRNA and the paired gene was calculated as follows: 1) the gRNA midpoint (gRNA_mid) was defined as (gRNA_start + gRNA_end)/2, the gene start coordinate (gene_start) was defined as the start coordinate for genes on the '+' plus strand, and end coordinate for genes on the '-' strand. 'ep_length' was calculated as gRNA_mid - gene_start.

*Effect size comparison between targeting and control gRNAs*
For each gene with a TSS- or validated enhancer-targeting gRNA, we compared the avg_logFC of expression for the respective gene between TSS-control, enhancer-control, pRE-targeting, and NT-control gRNAs (FDR < 0.01), using a one-way ANOVA followed by Tukey's HSD with Bonferroni correction (adj. p-value). Significant differences in the change in gene expression were defined as adj. p-value < 0.05.

*Interaction distance versus effect size*
Using all significant pRE-targeting gRNA gene pairs, the avg_logFC and effect size (avg_logFC*(1-FDR)) were plotted versus the log10-transformed ep_length (log10(abs(ep_length+1))). Spearman correlation $R^2$ values were calculated using the 'stat_cor' function from the 'ggpubr' R package.

*Nearest gene prediction analysis*
For each potential pRE-gene pair, we calculated the number of genes "skipped" by the element to regulate the gene as follows. First, for the significant pRE-gene connections, we defined the start and end coordinates for a given element and the start and end coordinates, and strand, for the paired gene. Next, we counted the number of genes detected in the gene expression dataset for which the entire gene body was contained within the region between the element and the connected gene. We repeated this for all significant pRE-gene connections.

*Comparison to microC looping*
We obtained chromatin contact data (**Table S19**) and intersected all targeted pREs, TSS regions (+/- 1kb) of every gene and all genes for which a differential expression test was performed, separately extended by +/- 500bp with anchor 1 and anchor 2, using bedtools window -w 500. We then quantified the number of pREs, TSSs, and genes with at least one chromatin contact, defined as at least one intersection with anchor 1 or anchor 2. Next, for all regions that intersected a region in the anchor 1 set, we quantified the number of pRE-gene pairs for which the corresponding contact in the anchor 2 set overlapped either the same TSS/gene or a different TSS/gene. We repeated this for pREs intersecting the anchor 2 set with comparison of contacts for TSSs/genes in the anchor 1 set.

*Gene overrepresentation and transcription factor enrichment tests*
The union set of all genes with at least one significant pRE link (FDR < 0.01, N=196) were queried using the 'enrichr' function from the 'enrichR' R package with default parameters and the following databases: 'DisGeNET', 'ENCODE_and_ChEA_Consensus_TFs_from_ChIP-X', 'ClinVar_2019', 'MSigDB_Hallmark_2020', and 'OMIM_Disease'. Over-represented pathways were defined as adj. p-value < 0.05.

*Chromatin accessibility of significant screen regions in IPF vs unaffected control tissue*
We obtained ATAC-seq peak calls from GSE180242 (*76*). We intersected the pREs connected to at least one gene with the peak calls using bedtools intersect. We then performed a Student's t-test comparing the fold change in chromatin accessibility for all overlapping peaks in IPF lung tissue vs unaffected control lung tissue using the t.test function in R.

*Single cell screen versus individual gRNA validations*
For the 10 pRE-gene connections with at least one significant individual gRNA validation, we calculated the Spearman correlation ($R^2$) and p-value between the change in mRNA expression measured via RT-qPCR (DDCt) versus the gene expression change observed in the single cell screen (avg_logFC of the most significant gRNA-gene connection per DHS) using the 'stat_cor' function from the 'ggpubr' R package.

*Differential expression of linked genes in primary lung tissue single cell RNA-sequencing data*
We obtained differential gene expression results from single cell profiling of healthy lung tissue and of lung tissue from individuals with IPF (GEO accession: GSE135893) (*77*). For each cell type and/or disease context, we compared the effect size (avg_logFC*(1-pval_adj) between pRE-connected genes and other genes using a permutation test framework. First, we calculated the difference in effect size for pRE-connected genes versus all other genes in the dataset. Then, we randomly permuted the genes in each group and calculated the difference in the mean effect size. We repeated this 10,000 times and calculated a one-tailed p-value as the number of times the permuted difference was greater than the observed difference divided by the number of permutations.

*Individual gRNA validations*
Oligos containing protospacer sequences were synthesized by IDT and cloned into pLV_hU6-sgRNA_hUbC-GFP-P2A-PuroR (Addgene plasmid #162335). Sanger sequencing was used to confirm the identity of the gRNA. Lentivirus was generated as described above. dCas9-KRAB expressing HFF cells were seeded onto TCP and transduced on day 0. 24 hours post-transduction, lentivirus was removed. Antibiotic selection was applied and cells were harvested eight days post-transduction. mRNA was isolated using the Norgen Total RNA Purification Kit (#17250) according to the manufacturer's protocol. 100 ng mRNA was used as input for cDNA amplification using the Invitrogen™ SuperScript™ VILO™ cDNA Synthesis Kit. For RT-qPCR, each reaction contained 1 uL cDNA, 7 uL H20, 1 uL Taqman probe for TBP, 1 uL Taqman probe for gene of interest, and 10 uL Quantabio PerfeCTa FastMix II. Delta delta Ct analysis was performed in Microsoft Excel. Graphpad Prism was utilized to conduct one-way ANOVA tests followed by Tukey's HSD for post-hoc testing. Significance is reported in Fig.s as follows: *p-value < 0.05, **p-value < 0.01, ***p-value < 0.001. Taqman probe information is provided in **Table S16**.

**Supplementary Text**

Supplementary Text 1. Strand bias of gRNAs in the MYH9 screen.

In the MYH9 CRISPRi screen, we noted strong enrichment of Z-scores in a strand-dependent fashion, wherein nearly any gRNA that fell in the *MYH9* gene body on the non-template (+) strand showed a strongly repressive phenotype (**Fig. S4**). This intragenic strand bias is likely related to steric hindrance of dCas9 interfering with RNA polymerase, as recently described(*114*)(*107*). For subsequent analyses we only utilized gRNAs targeting the coding strand, which do not lead to the non-specific steric effects of dCas9 on MYH9 transcription (2,863 gRNA total).

Supplementary Text 2. Accessibility of pREs regulating growth and migration phenotypes varies across diverse cell types.

We intersected the pREs with accessible chromatin regions in ENCODE biosamples (N=95) and observed three clusters of pREs: 1) pREs that overlap accessible regions in all or most biosamples ("ubiquitous"; N=79 regions), 2) pREs that overlap accessible regions in a majority of biosamples ("prevalent", N=23 regions), and 3) pREs that overlap accessible regions only in biosamples of similar cell or tissue types ("lineage-specific"; N=13) (**Fig. S10**; **Methods**). Additionally, we observed similar proportions of pREs that also overlap an accessible region for cell types from similar lineages and/or with similar culture conditions (e.g., suspension versus adherent) (**Table S10**). For example, 99.1% (N=114/115) and 96.5% (N=111/115) of pREs overlap accessible chromatin regions in fibroblasts from thigh (AG04449) and in human skeletal muscle myoblasts (HSMM), respectively, which are highly adherent cell types. H7 human embryonic stem cells (H7-hESC) are cultured on soft substrates and 49.6% (N=57/115) of pREs are accessible in that cell type, consistent with the intermediate level of H3K27ac signal of the MYH9 intronic mechanoenhancer. In contrast, only 19.1% (N=22/115) of pREs are accessible in K562 cells and Jurkat cells, with similar overlap observed in other blood lineage cell types (e.g., monocytes, GM12878).

Supplementary Text 3. pRE accessible across various cell- and tissue-types regulates many genes.

Of the "ubiquitous" pREs we identified in Fig. S10, chr1:28648521-28649567 was linked to >10 genes in the scRNA-seq screen (**Fig. S12**). After thresholding to remove small effect sizes, we considered 10 pRE-gene links, with 5/10 links having at least two gRNAs for each connection supporting the change in expression (**Fig. S12A-B, Methods**). Notably, the pRE also overlaps five non-promoter FANTOM5 TSS peaks, all of which are annotated on the plus strand and could potentially indicate the presence of enhancer RNAs (**Fig. S12A**). We also observed significant enrichment of eight TFs in the promoters of the genes, including ZMIZ1 and ZNF384 (**Fig. S12C**, **Methods**). ZMIZ1 has previously been shown to regulate p53 signaling (*115*) and ZNF384 regulates key ECM components and regulators including *MMP1*, *MMP3*, *MMP7*, and *COL1A1* (*116–119*).

Supplementary Text 4. Example pREs that regulate cell migration.

Validated pRE-gene linkages include intronic and intergenic regions of many genes known to regulate cell migration (*27*, *79*, *120–122*). For example, a region ~30kb downstream of the gene *CYR61*/*CCN1*, a canonical YAP/TAZ transcriptional target (*79*) has been strongly linked to migratory phenotypes (*72*) (**Fig. 4G**, **Fig. S16**). Epigenetic repression of this region led to ~50% reduction in *CYR61* expression. Notably, this pRE overlaps an annotated TEAD1/3 binding site, which may suggest that YAP/TAZ binding may facilitate the gene expression and phenotypic changes. Additionally, perturbation of an intronic region of *RASGRP1* led to decreased cell migration, decreased expression of *RASGRP1* and *SPRED1*, and increased expression of *FAM98B* (**Fig. S18**). *RASGRP1* is a Ras GEF that activates the ERK/MAPK cascade and can tune migration, and SPRED1 also regulates ERK/MAPK cascade activation but has also been implicated in Neurofibromatosis Type 1-Like Syndrome and Noonan Syndrome, and was previously shown to reduce formation of F-actin stress fibers by inhibiting TESK1 (*120*). *FAM98B* is a positive regulator of cell proliferation and has been implicated in colorectal cancer progression (*121*). Notably, the second intron of RASGRP1 contains functionally-validated single nucleotide variants implicated in systemic lupus erythematosus (SLE) (*123*, *124*). Likewise, a region ~4.6kb upstream of *RANGAP1* and ~10.7kb upstream of *ZC3H7B* regulated the expression of both genes (**Fig. S19**). Importantly, *RANGAP1* is a crucial cytoplasmic-nuclear shuttling mediator, and it's mechanoactivation further suggests a feedback loop wherein increased ECM stiffness can lead to enhancer activation, increased *RANGAP1* expression, and increase nuclear to cytoplasmic shuttling, thereby reinforcing and amplifying mechanical signals. In contrast, a region ~200kb upstream was found to regulate DUSP4, which inhibits the ERK/MAPK cascade by negatively regulating ERK & JNK kinases (*122*, *125*) (**Fig. S17**).

Supplementary Text 5. Example pREs that regulate cell proliferation.

Similar to pREs that regulated migration, we also identified the target genes for pREs regulating cell proliferation. For example, one particular pRE, which when perturbed with $dCas9^{KRAB}$ significantly regulated cell proliferation and is located ~135kb away from *CTGF*/*CCN2*, was found to function as a mechanoenhancer of *CTGF*. Repression of this pRE led to ~90% reduction in *CTGF* expression (**Fig. 5C-D**). Interestingly, this pRE is near the annotated promoter of a lncRNA but overlaps an ENCODE-predicted distal regulatory element, demonstrates physical interaction with the *CTGF* promoter (ChIA-PET), and overlaps both H3K4me1 and H3K27ac peaks. When HFF cells are cultured on increasingly stiff surfaces, the pRE becomes more accessible and *CTGF* expression increases, supporting a gene expression response to the mechanical stimulus (**Fig. 5E-F**). *CTGF* is a key downstream transcriptional target of YAP/TAZ following mechanical activation (*13*), and knockdown of *CTGF* blocks the YAP-dependent growth phenotype. Our results further suggest one important genomic mediator of this stiffness-dependent change is a distal mechanoenhancer that helps drive increased *CTGF* expression.

Likewise, we validated other novel mechanoresponsive pREs that regulate genes known to be key drivers of cellular growth, including *NF2* (**Fig. S16C-D**), *SKP2* (**Fig. 4F**, **Fig. S20**), and *RFLNB* (**Fig. S17C-D**). *RFLNB* has previously been implicated in perinuclear actin organization, *SKP2* is a potent cell cycle regulator driven by YAP/TAZ activity (*126*), and NF2 is a potent tumor suppressor. Accordingly, we identified that epigenetic repression of the pREs linked to these genes led to either increased cell growth (*NF2*) or decreased cell growth (*RFLNB*, *SKP2*) in the bulk screen and decrease in expression of the target genes by both scRNA-seq and qRT-PCR. Additionally, these genes showed differential expression and the pREs were differentially accessible following exposure to varying culture substrate stiffness (**Fig. 1**).

**Figs. S1-S24**

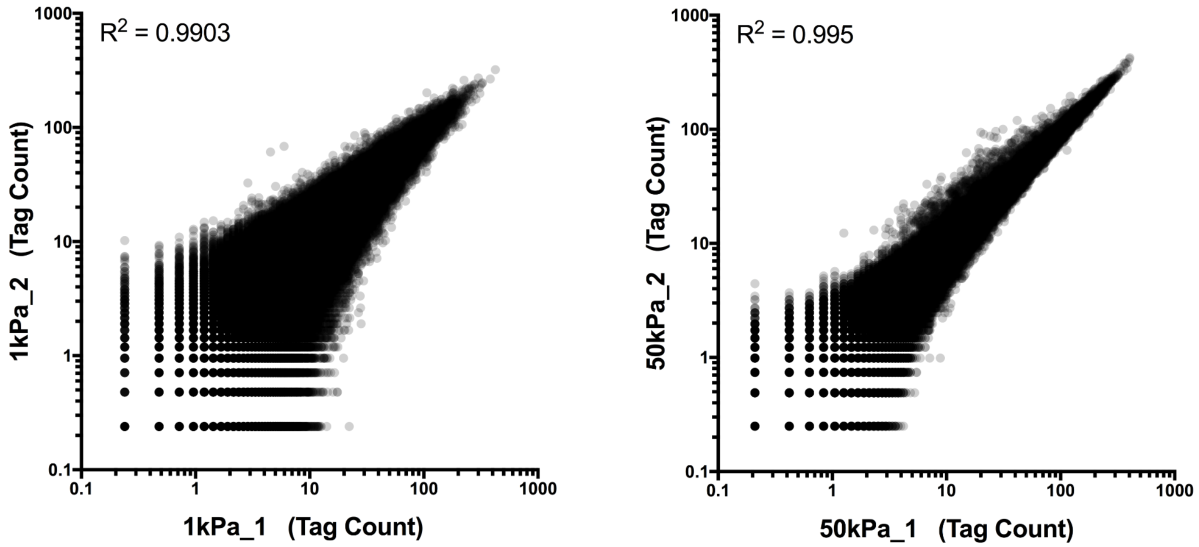


**Fig. S1. Correlation of biological replicates for ATAC-seq data.** Correlation of ATAC-seq tag counts from biological replicates for soft (1 kPa, left) and stiff (50 kPa, right) conditions. Each point corresponds to a peak call from the merged peak set.

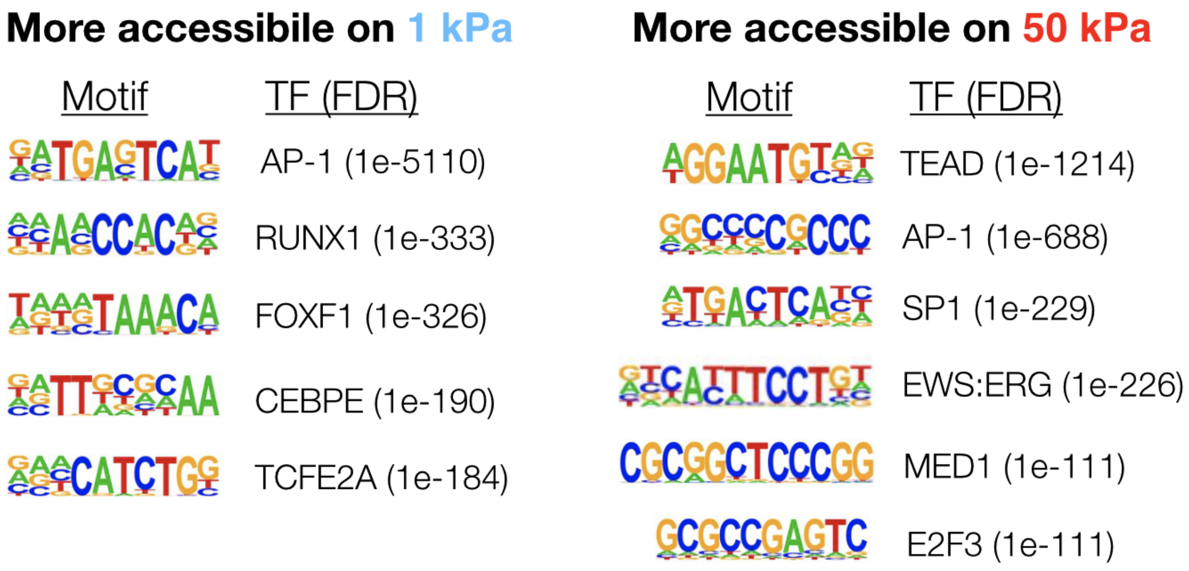


**Fig. S2. Motifs from de novo motif analysis.** Individual de novo motifs that were enriched in 1 kPa or 50 kPa peaksets compared to sequence content matched genomic background (as assayed by HOMER). FDR values from HOMER for a given de novo MOTIF are listed in brackets.

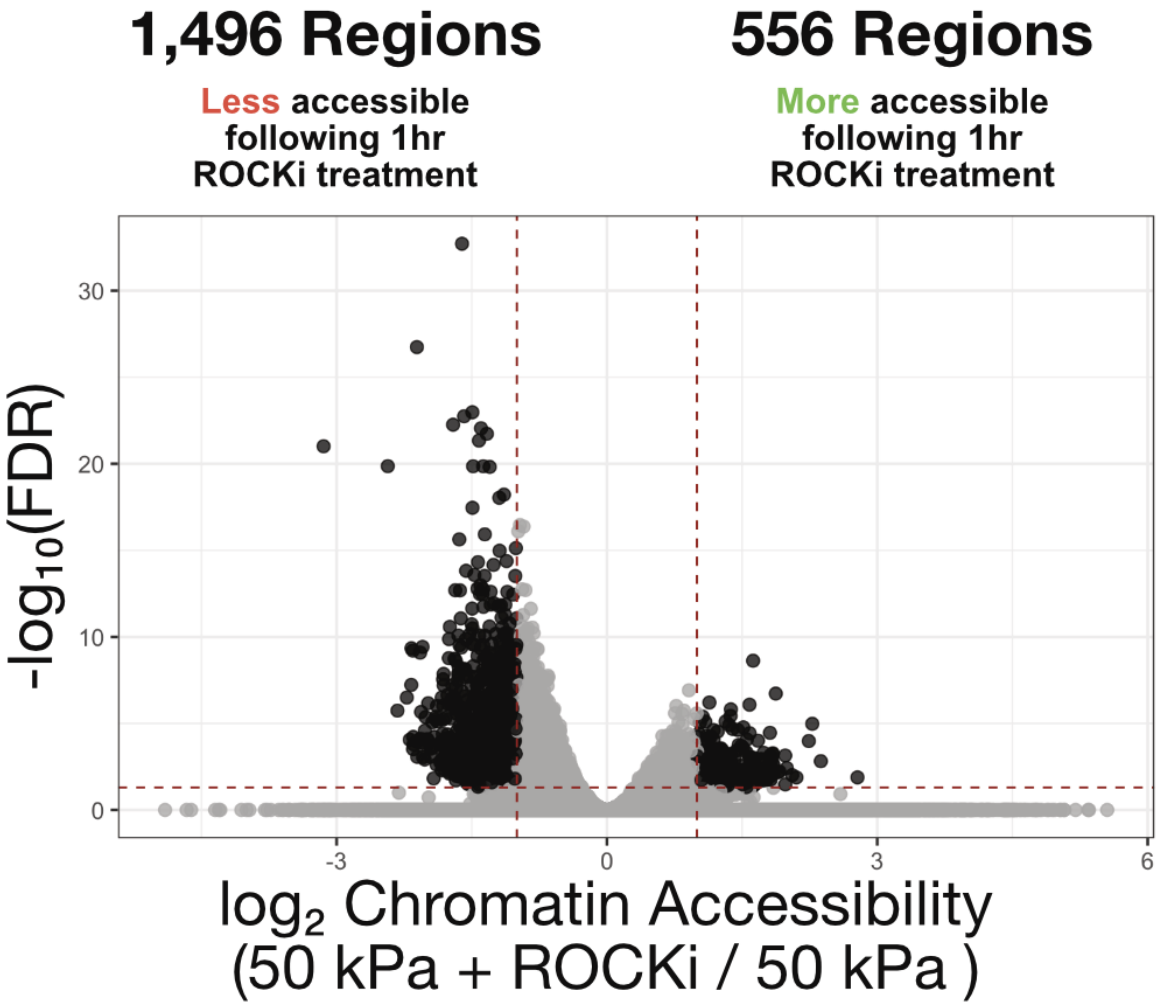


**Fig. S3. Changes in chromatin accessibility following 10 µM Y-27632 ROCKi treatment for 1 hour prior to harvest.** HFF cells were cultured for 20 hours on 50 kPa polyacrylamide hydrogels, with DMSO or 10 µM Y-27632 added for the last hour of culture. ATAC-seq was then performed and each open chromatin region is represented as an individual point.

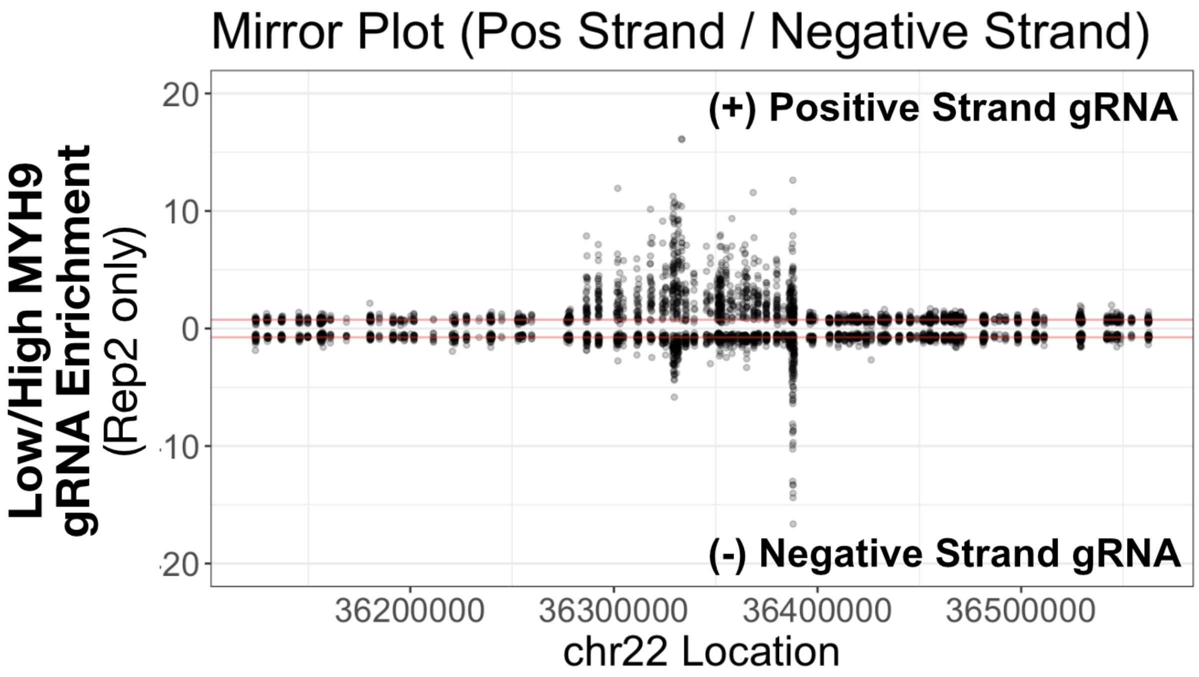


**Fig. S4. CRISPR interference leads to strand bias in screening positive strand protospacer gRNA.** Mirror plot of the average gRNA enrichment in low/high MYH9 expressing cells from the *MYH9* screen across the genomic location, where gRNA points plotted upwards are (+) stranded, and gRNA points plotted downward are (-) stranded.

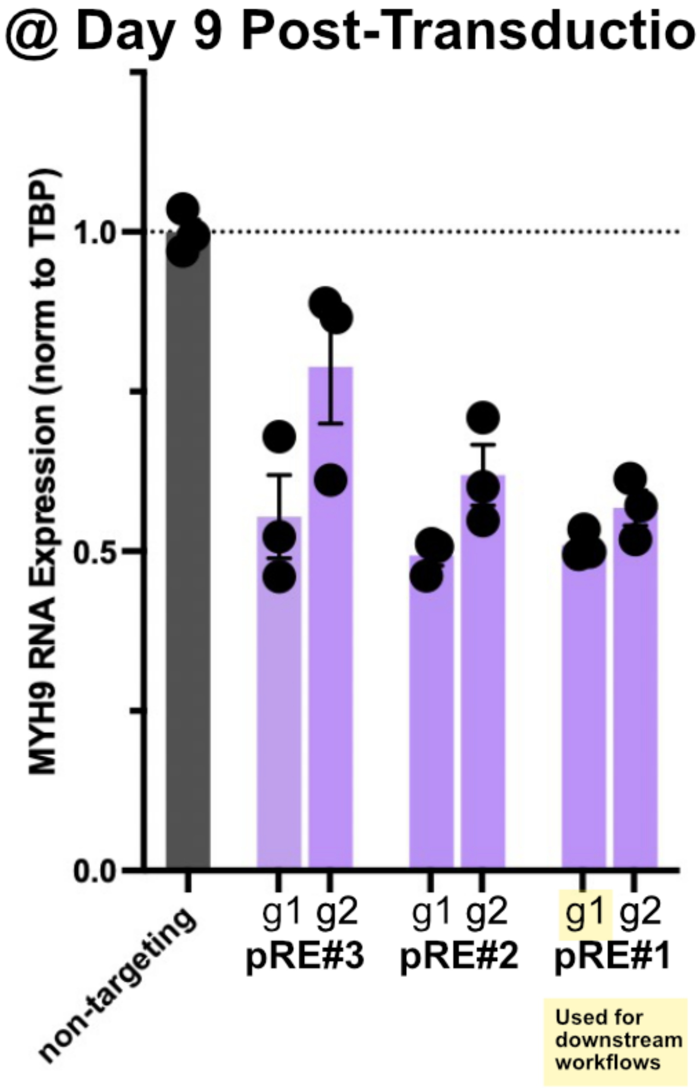


**Fig. S5. Singleton validation for CRISPRi MYH9 locus screen intron 3 hits.** Singleton validations of MYH9 repression found across the three hit pRE from the MYH9 screen at Day 9 timepoints (N=3 reps/group).

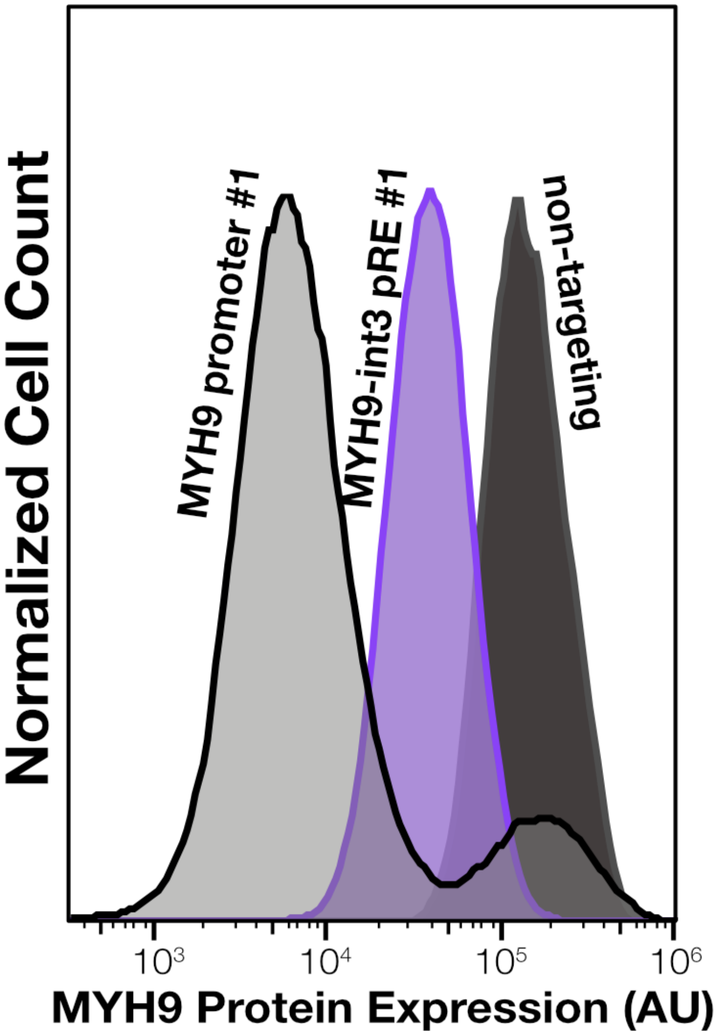


**Fig. S6. MYH9 protein levels in HFFs 10d after CRISPRi perturbation of promoter and intronic mechanoenhancer.** HFFs were transduced with dCas9-KRAB and either a non-targeting gRNA, a MYH9-intron 3 pRE#1 gRNA, or a MYH9 prom gRNA. Cells were fixed, immunostained for MYH9, and then subjected to flow cytometry. 42,602 cells were counted for the non-targerting gRNA, 187,733 cells for the MYH9 intron 3 pRE#1 gRNA (~28% of mean non-targeting expression), and 109,402 cells for the MYH9 promoter gRNA (~18% of mean non-targeting expression, though with a biphasic population).

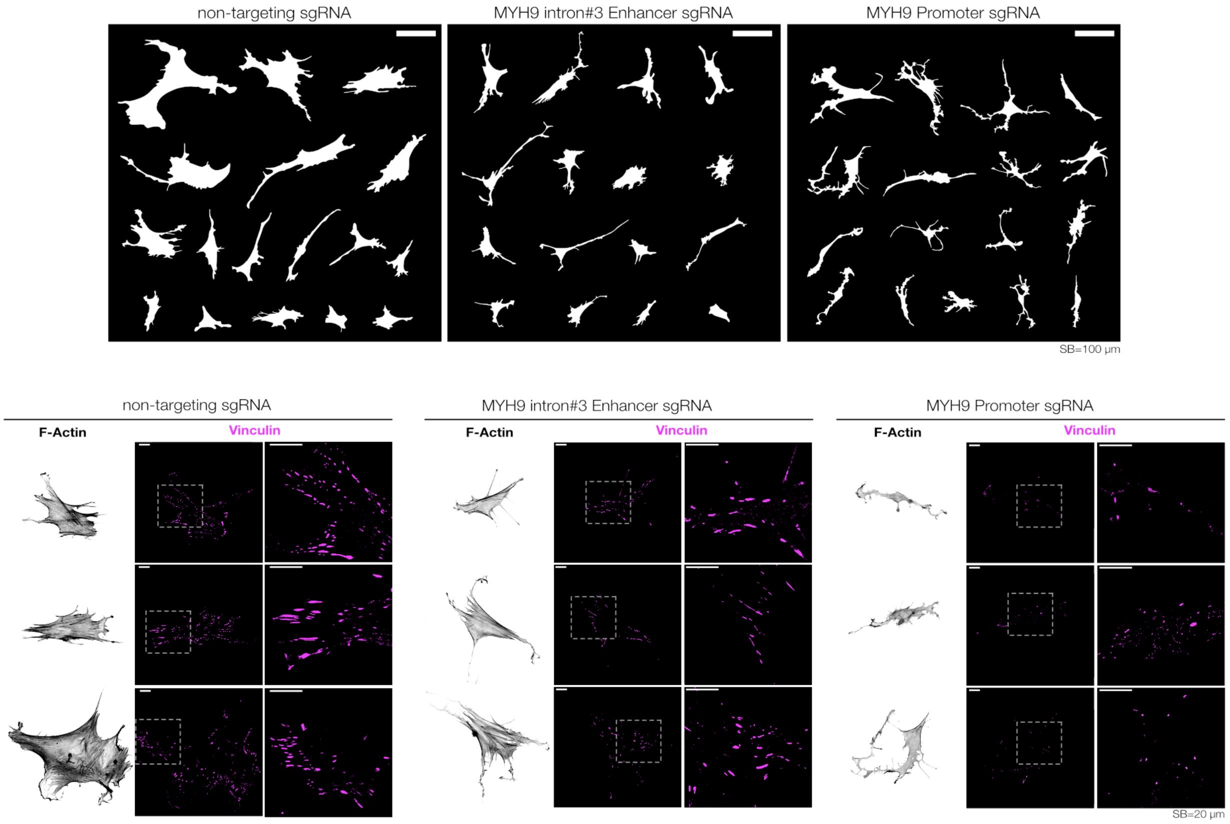


**Fig. S7. Representative morphometric and focal adhesion images of HFF cells following MYH9-intron 3 enhancer repression or MYH9 promoter repression with CRISPRi.** Representative outlines of cell area and vinculin immunostaining for HFF cells following 9 days post-transduction of dCas9-KRAB constructs and either a non-targeting gRNA, a MYH9-intron 3 enhancer targeting gRNA, or a MYH9 promoter targeting gRNA.

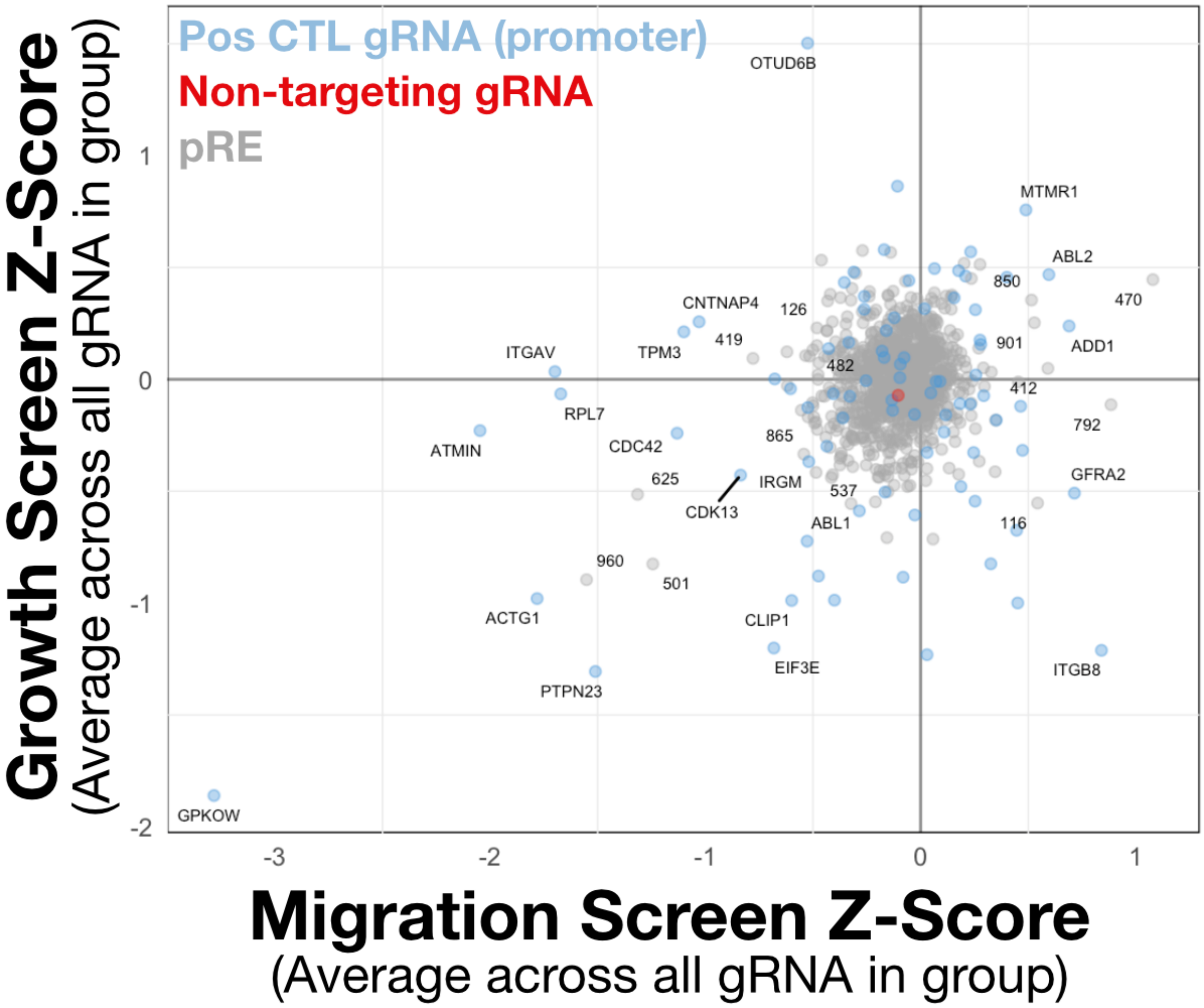


**Fig. S8. Screen effect size (Z-score) for each pRE across both migration screens and growth screens.** Non-targeting gRNA and positive control gRNA of genes changing migration are shown in red and blue respectively. Points shown are average Z-score across all gRNA in each region/target.

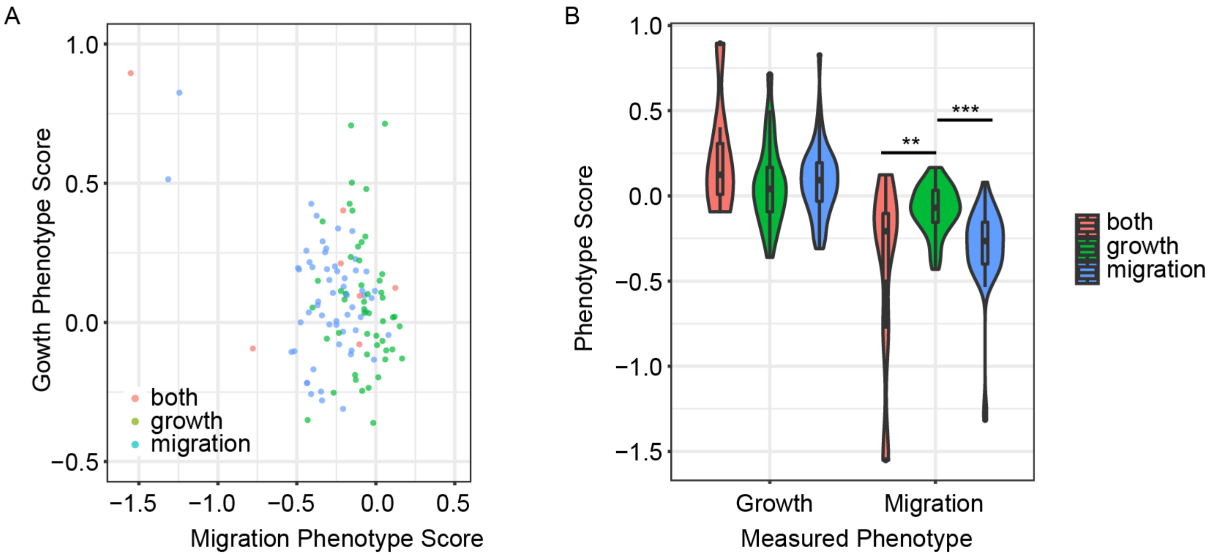


**Fig. S9. Comparison of growth and migration phenotype scores. (A)** Scatterplot comparing the phenotype scores between growth and migration phenotypes of all perturbed pREs. pREs significant in both screens, only the growth screen, only the migration screen, are colored in red, green, and blue, respectively. The phenotype scores between the screens were not correlated for pREs regulating only migration ($R^2$ = 0.0098, p-value = 0.46), both phenotypes ($R^2$ = 0.082, p-value = 0.56), and only regulating growth ($R^2$ = 0.049, p-value = 0.12). **(B)** Comparison of the phenotype scores between regions significant in only the growth, only the migration, or in both screens, colored in green, blue, and red, respectively (Migration phenotype: One-way ANOVA, p-value = 3.1e-06; Tukey's post-hoc tests: both vs migration, adj. p-value = 0.485; both vs growth, adj. p-value = 0.002; migration vs growth, adj. p-value = 0.00001; Growth phenotype: One-way ANOVA, p-value = 0.264). For box plots within violin plots, boxes show the quartiles with a line at the median. Lines extend to 1.5 times the interquartile range, and dots show outliers. Sample sizes for each group are as follows: both, N=7; growth, N=50; migration, N=58. For clarity, significance indicated in plot only for comparison with adj. p-value < 0.05. ** indicates p < 0.01 and *** indicates p < 0.001.

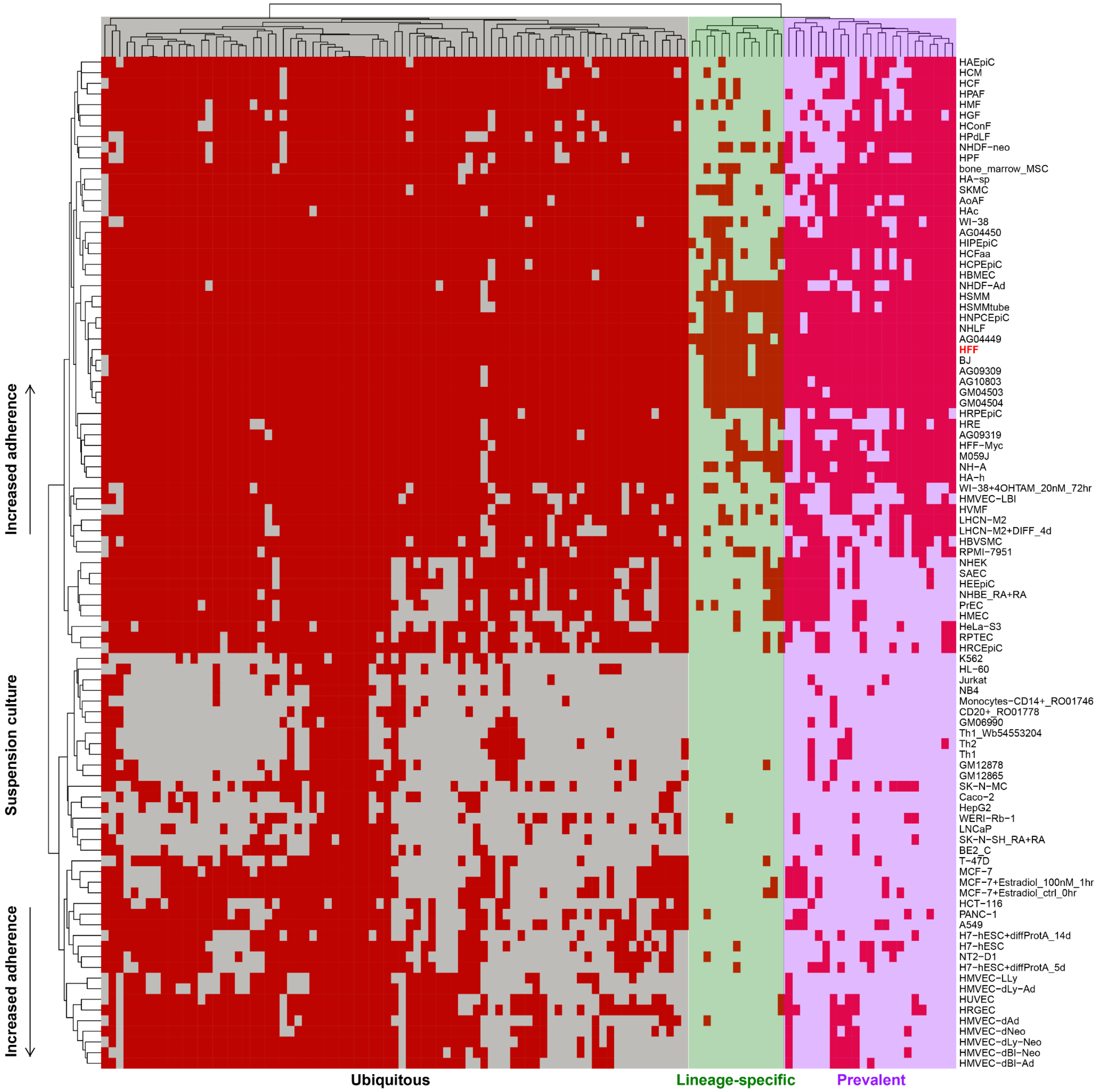


**Fig. S10. Overlap of mechanosensitive regulatory elements and accessible chromatin regions in ENCODE biosamples.** Heatmap of significant screen pREs that regulate migration and/or proliferation (columns) by ENCODE biosamples (rows) with coloring of red or white indicating the screen region overlaps or does not overlap an accessible chromatin region in that biosample, respectively, with data clustered by rows and by columns. “Lineage-specific”, “Prevalent”, and “Ubiquitous” regions indicated by green, purple, and grey, respectively. The biosample corresponding to the same cell type queried in the screens (‘HFF’) is denoted in bold red text.

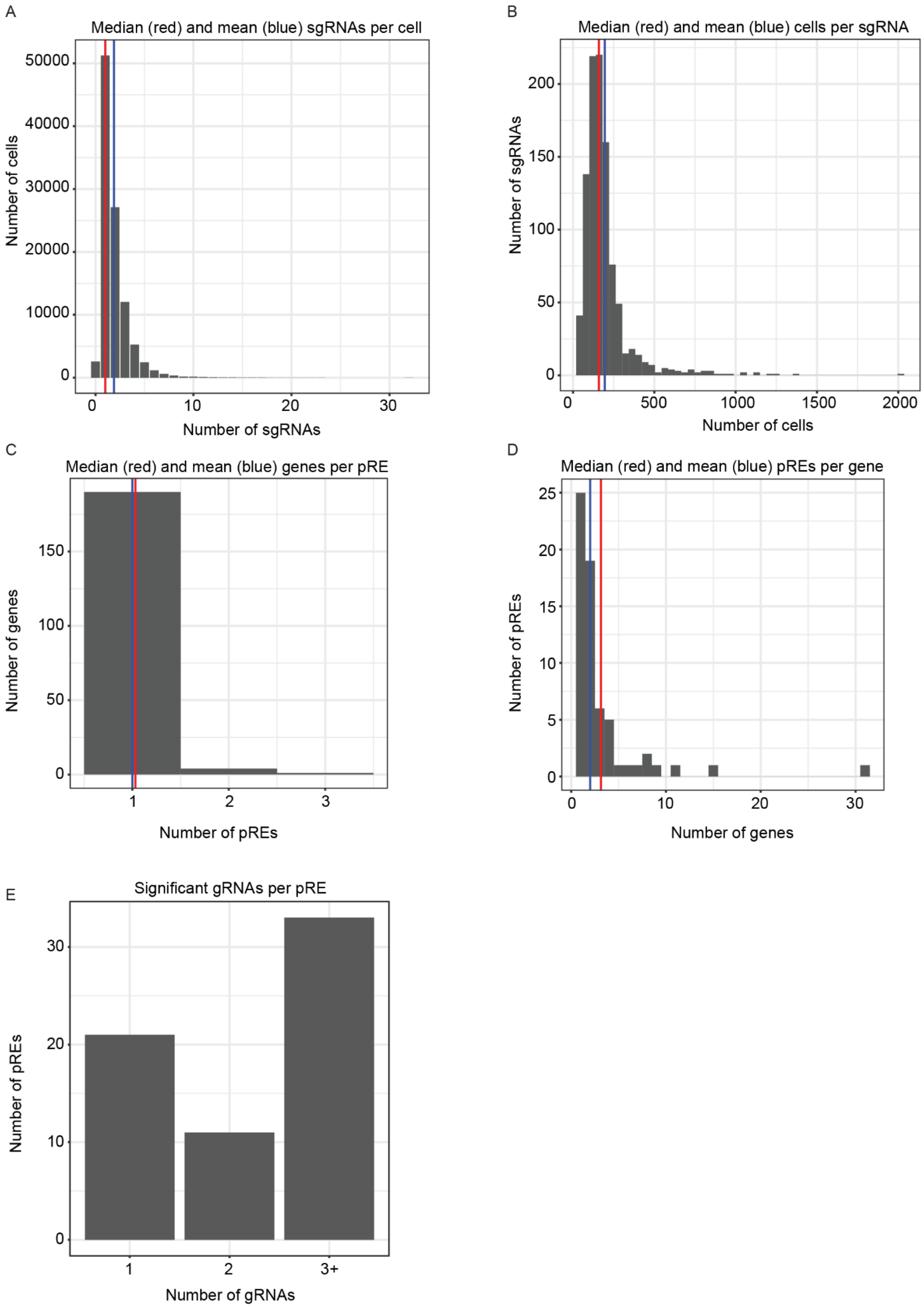


**Fig. S11. Distribution of coverage, MOI, pREs per gene, genes connected to each pRE, and significant gRNAs per pRE. (A)** Histogram of number of gRNAs observed in a given cell (MOI; mean = 1.90, median = 1) **(B)** Histogram of cells in which a given gRNA was observed (coverage; mean = 195, median = 159). **(C)** Distribution of the number of genes that significantly change in expression upon perturbation of each pRE (FDR < 0.01, mean = 1.03, median = 1). **(D)** Distribution of the number of pREs that when perturbed led to a significant change in expression of a given gene (FDR < 0.01, mean = 3.11, median = 2). **(A-D)** Mean and median MOI indicated by vertical blue and red lines, respectively. **(E)** Number of pREs with 1, 2, or 3 or more ('3+') significant gRNAs (N_1 = 21, N_2 = 11, N_'3+' = 33).

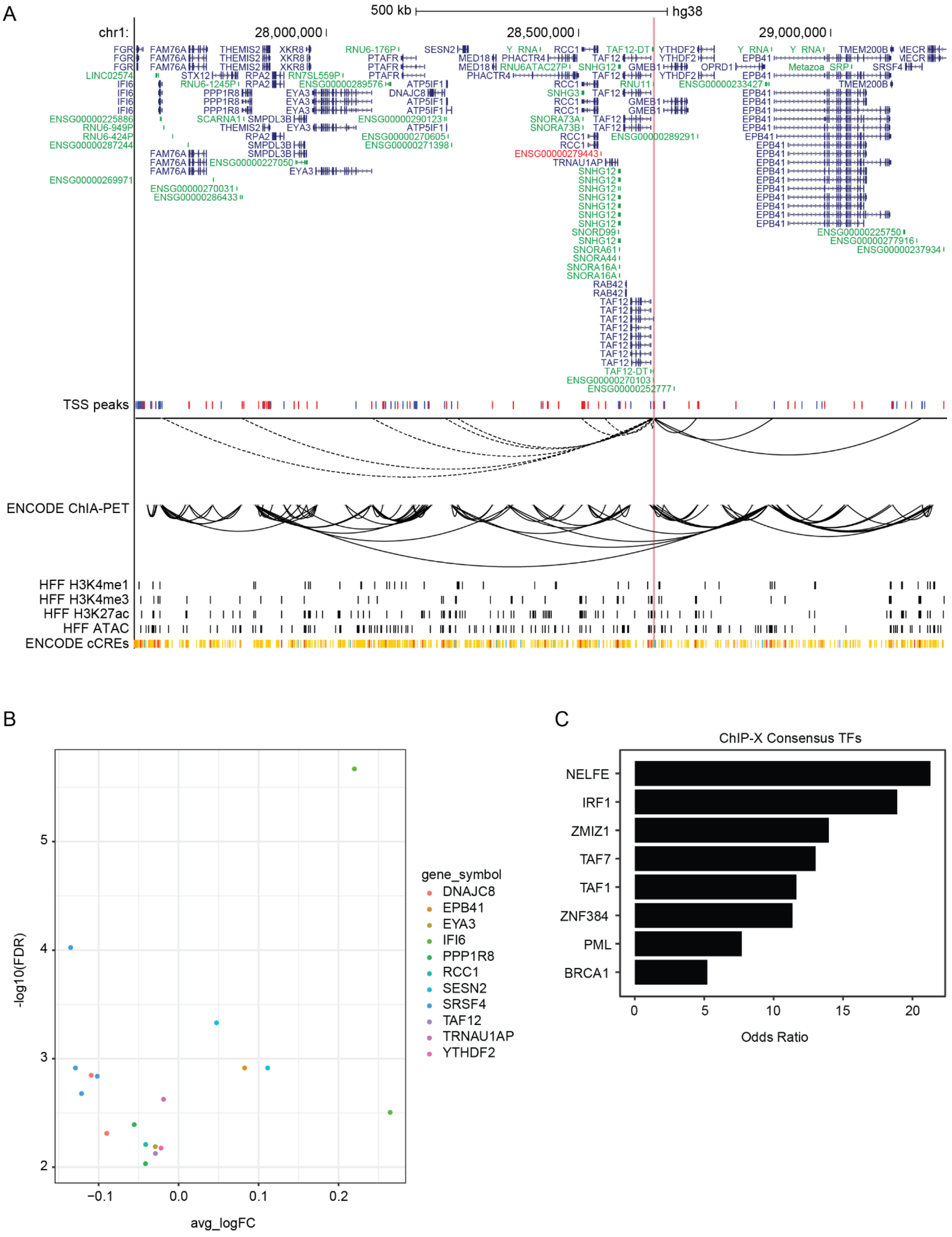


**Fig. S12. pREs with many gene connections. (A)** Browser track showing chr1:28648521−28649567. pRE−gene connections are shown below the 'TSS peaks' with dashed and solid lines indicating the gene is located to the left or right of the pRE, respectively. **(B)** pRE-gene connections for pREs linked to 11 genes (**Methods**). Each point represents an individual gRNA-gene pair and the coloring corresponds to the connected gene. **(C)** Odds ratios for enrichment of transcription factor motifs in differentially expressed genes upon perturbation of chr1:28648521-28649567 (adj. p-value < 0.1).

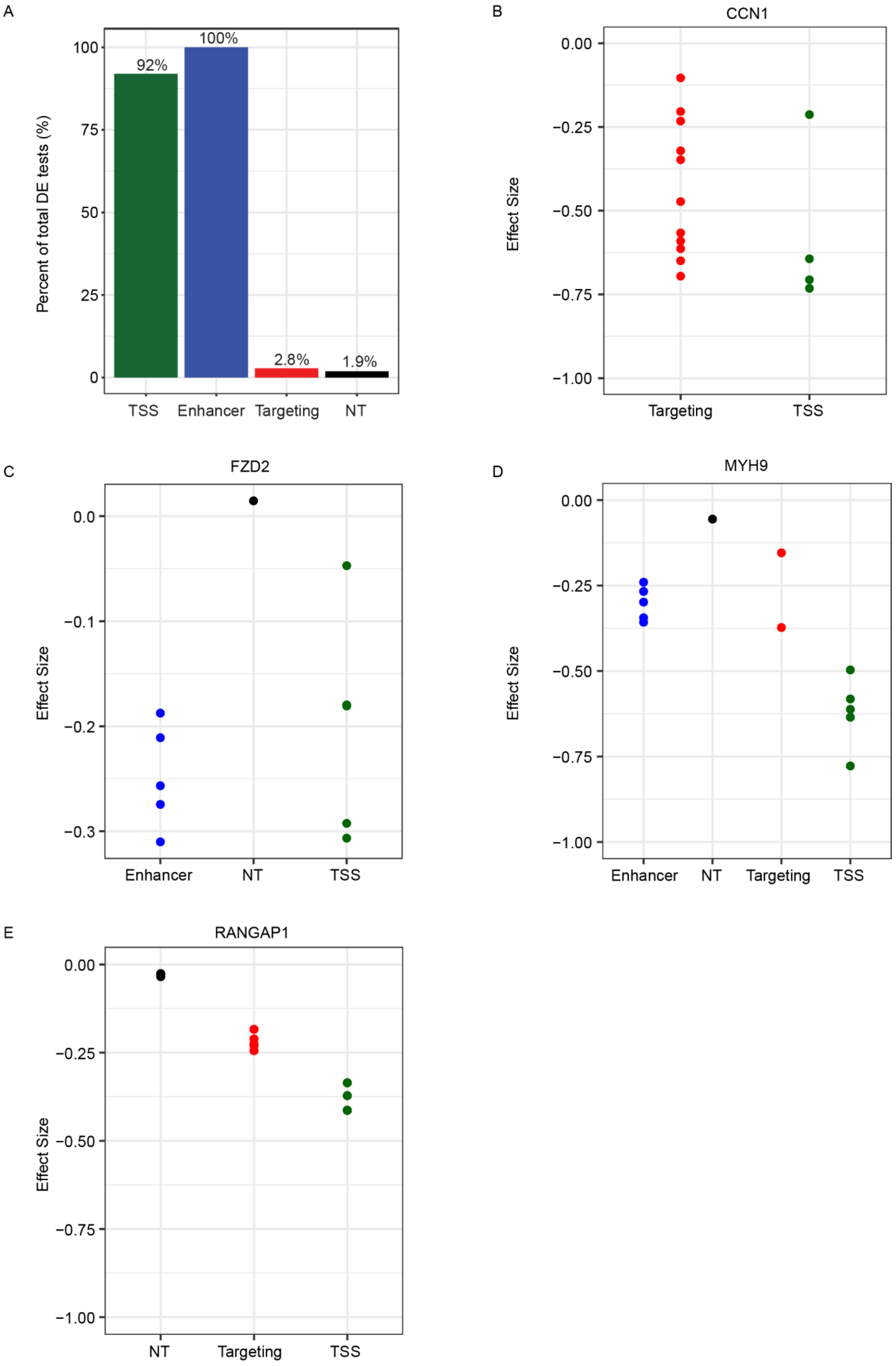
A
Percent of total DE tests (%)
100
75
50
25
0
92%
100%
2.8%
1.9%
TSS
Enhancer
Targeting
NT
B
CCN1
Effect Size
0.00
−0.25
−0.50
−0.75
−1.00
Targeting
TSS
C
FZD2
Effect Size
0.0
−0.1
−0.2
−0.3
Enhancer
NT
TSS
D
MYH9
Effect Size
0.00
−0.25
−0.50
−0.75
−1.00
Enhancer
NT
Targeting
TSS
E
RANGAP1
Effect Size
0.00
−0.25
−0.50
−0.75
−1.00
NT
Targeting
TSS

**Fig. S13. Comparison of perturbation effects by gRNA type. (A-E)** Results for non-targeting ('NT'), pRE-targeting ('Targeting'), TSS-targeting positive controls ('TSS'), and enhancer-targeting positive control ('Enhancer') gRNAs, colored by black, red, green, and blue, respectively. **(A)** Percent of total gRNA-gene differential expression tests with FDR < 0.01. Percent noted above each bar. **(B-E)** Comparison of effect size (Methods) for significant gRNA-gene connections between gRNA types. **(C-E)** Adjusted p-value from Tukey's post-hoc test noted following each comparison. **(B)** No significant difference in effect size for CCN1 expression for TSS vs Targeting (t-test, p-value = 0.3623). **(C)** Significant difference in effect size for FZD2 expression for Enhancer vs NT (adj. p-value = 0.0453). No significant difference for TSS vs NT (adj. P = 0.0972) and TSS vs Enhancer (adj. p-value = 0.6556). **(D)** Significant difference effect in effect size for MYH9 mRNA expression for TSS vs Enhancer (adj. p-value = 0.0017), TSS vs NT (adj. p-value = 0.0015), and TSS vs Targeting (adj. p-value = 0.0053). No significant difference for Targeting vs Enhancer (adj. p-value = 0.9582), NT vs Enhancer (adj. p-value = 0.1366), and Targeting vs NT (adj. p-value = 0.3121). **(E)** Significant difference in effect size for RANGAP1 mRNA expression for Targeting vs NT (adj. p-value = 1.18e-05), TSS vs NT (adj. p-value = 1.00e-07), and TSS vs Targeting (adj. p-value = 1.86e-05).

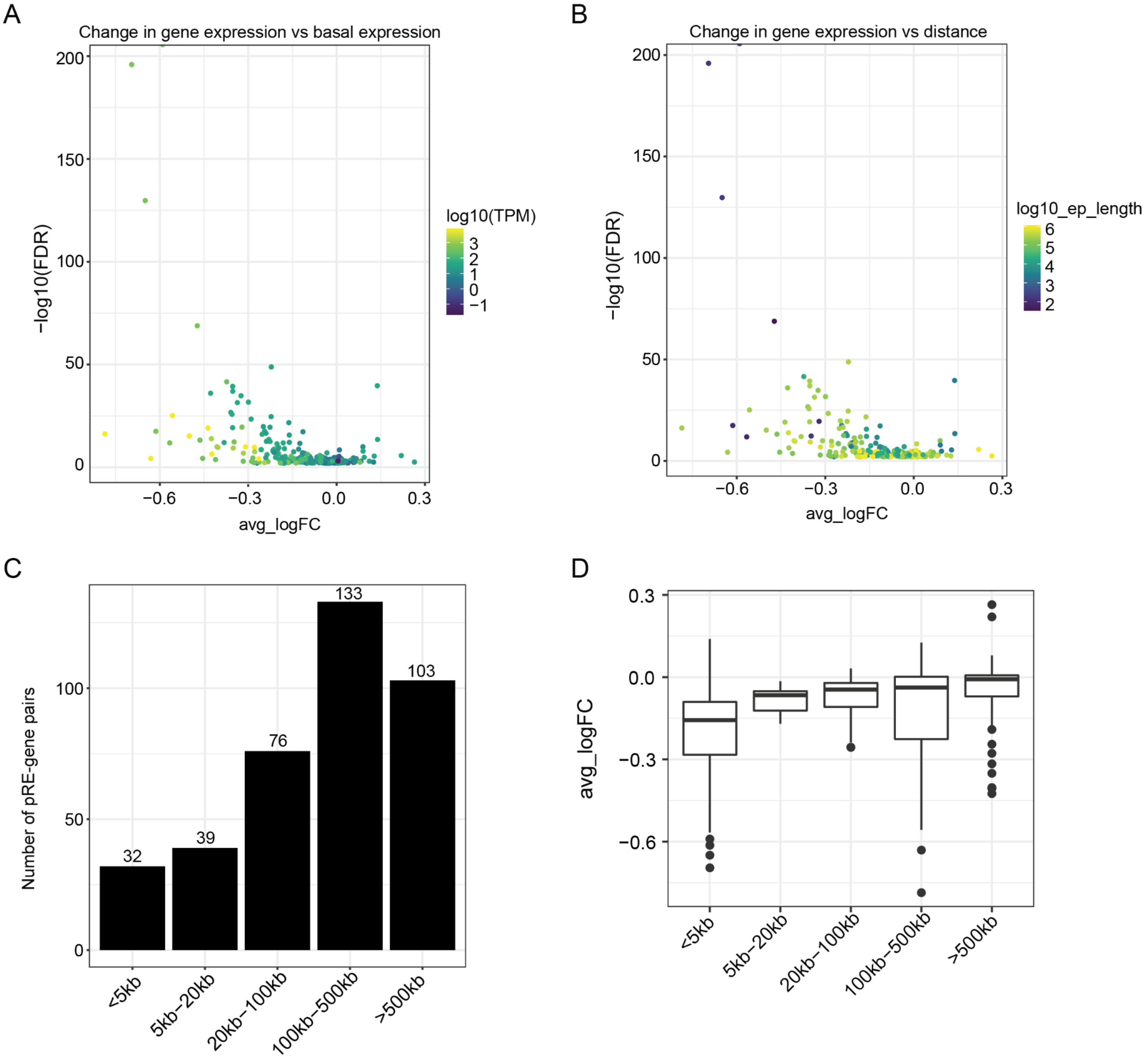


**Fig. S14. Characterization of interaction distance and basal expression levels for DHS-gene connections. (A)** Volcano plot of single cell screen results comparing the change in mRNA expression (avg_logFC) versus the significance (-log10(FDR)) for significant gRNA-gene connections (FDR < 0.01) and colored by basal level of mRNA expression in HFF cells. Note, mRNA expression values were log10-transformed prior to visualization. **(B)** Volcano plot of single cell screen results comparing the change in mRNA expression (avg_logFC) versus the significance (-log10(FDR)) for significant gRNA-gene connections (FDR < 0.01) and colored by the absolute distance between each pRE and gene. Note, the absolute distance was log-10 transformed prior to visualization. **(C)** Total count of gRNA-gene connections grouped by absolute distance between pRE and target gene (<5kb, 5kb-20kb, 20kb-100kb, 100kb-500kb, >500kb). **(D)** Comparison of avg_logFC of gene expression for gRNA-gene connections between groups in **(C)**. Boxes show the quartiles with a line at the median. Lines extend to 1.5 times the interquartile range, and dots show outliers. All comparisons versus '<5kb' are significant (adj. p-value < 0.05). Sample sizes for each group: <5kb = 32, 5kb-20kb = 103, 20kb-100kb = 76, 100kb-500kb = 133, >500kb = 103.

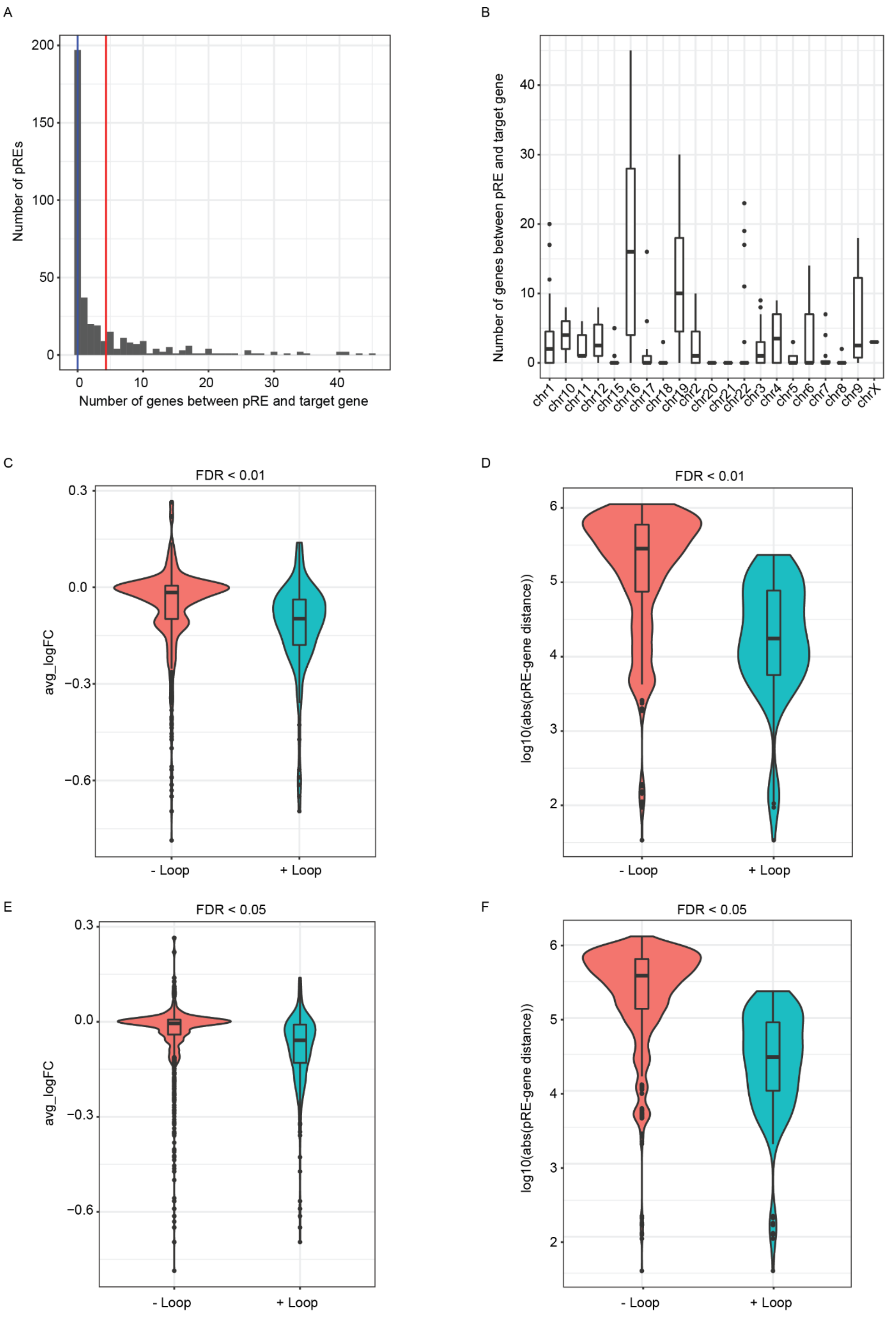
A
Number of pREs
Number of genes between pRE and target gene
B
Number of genes between pRE and target gene
chr1
chr10
chr11
chr12
chr15
chr16
chr17
chr18
chr19
chr2
chr20
chr21
chr22
chr3
chr4
chr5
chr6
chr7
chr8
chr9
chrX
C
FDR < 0.01
avg_logFC
- Loop
+ Loop
D
FDR < 0.01
log10(abs(pRE-gene distance))
- Loop
+ Loop
E
FDR < 0.05
avg_logFC
- Loop
+ Loop
F
FDR < 0.05
log10(abs(pRE-gene distance))
- Loop
+ Loop

**Fig. S15. Comparison between functional characterization DHS-gene connections and other prediction methods. (A)** Histogram of the number of genes located between the pRE and connected gene in the scRNA-seq screen. Blue line indicates mean, red line indicates median. **(B)** Box plot showing the number of genes located between the pRE and connected gene in the scRNA-seq screen for each chromosome. Boxes show the quartiles with a line at the median. Lines extend to 1.5 times the interquartile range, and dots show outliers. Sample sizes for chromosomes in order of x-axis from left to right: chr1, N=33; chr10, N=2; chr11, N=5; chr12, N=6; chr15, N=9; chr16, N=31; chr17, N=7; chr18, N=3; chr19, N=15; chr2, N=18; chr20, N=2; chr21, N=1; chr22, N=9; chr3, N=12; chr4, N=8; chr5, N=10; chr6, N=12; chr7, N=7; chr8, N=4; chr9, N=7; chrX, N=1. **(C-F)** Comparison of scRNA-seq links between pRE-gene pairs with chromatin loop observed in microC (+Loop) versus pRE-gene pairs without a chromatin loop (-Loop). The change in gene expression **(C)** and the distance between the pRE and linked gene **(D)** for pRE-gene pairs with FDR < 0.01 (N_+loop = 144, N_-loop = 885; gene expression: t-test, p-value = 1.195e-07; pRE-gene distance: t-test, p-value < 2.2E-16). The change in gene expression **(E)** and the distance between the pRE and linked gene **(F)** for pRE-gene pairs with FDR < 0.05 (N_+loop = 217, N_-loop = 2029; gene expression: t-test, p-value = 1.391e-12; pRE-gene distance: t-test, p-value < 2.2e-16). For **(D)** and **(F)**, the distance was log10-transformed prior to statistical test and for visualization. **(C-F)** For box plots within each violin plot, boxes show the quartiles with a line at the median. Lines extend to 1.5 times the interquartile range, and dots show outliers.

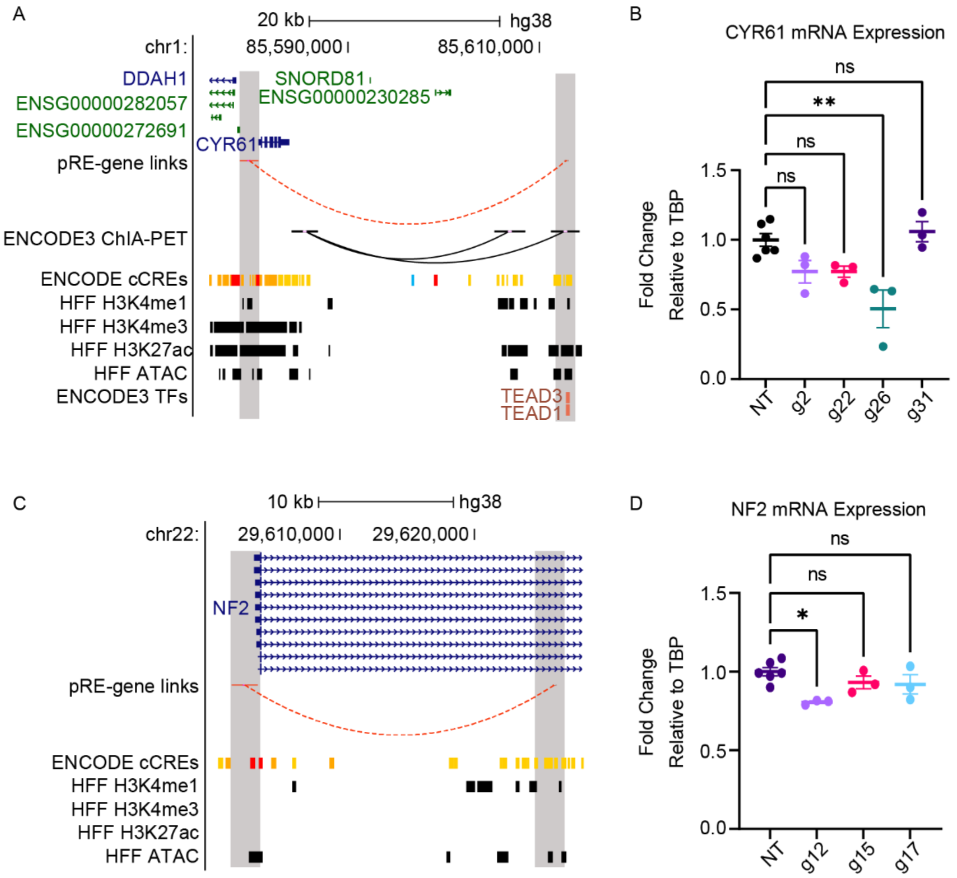


**Fig. S16. Close-range pRE-gene validation examples. (A)** Browser track of the region containing the gene CCN1 and the linked pRE with epigenomic and genomic annotations in HFF cells and across ENCODE3 biosamples. **(B)** CCN1 mRNA expression measured via RT-qPCR following individual delivery of gRNAs targeting the connected pRE in CRISPRi HFF cells (N=6 for NT, N=3 for all other gRNAs; 'NT' indicates non-targeting; 'ns' indicates not significant, adj. p-value >= 0.05; ** denotes adj. p-value < 0.01). Error bars indicate +/- 1 SEM, horizontal line indicates mean. **(C)** Browser track of the region containing the gene NF2 and the linked pRE with epigenomic and genomic annotations in HFF cells and across ENCODE3 biosamples. **(D)** NF2 mRNA expression measured via RT-qPCR following individual delivery of gRNAs targeting the connected pRE in CRISPRi HFF cells (N=6 for NT, N=3 for all other gRNAs; 'NT' indicates non-targeting; 'ns' indicates not significant, adj. p-value >= 0.05; * denotes adj. p-value < 0.05). Error bars indicate +/- 1 SEM, horizontal line indicates mean.

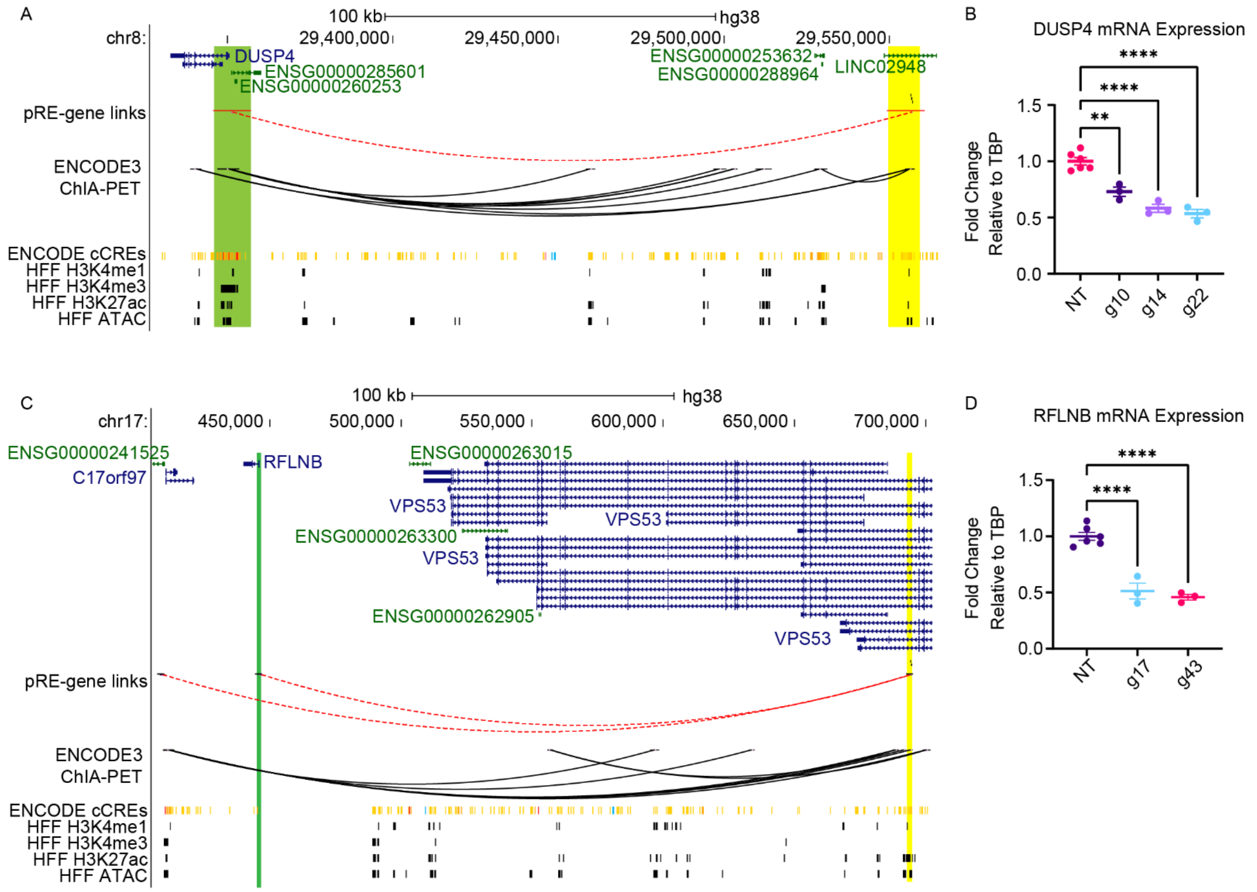


**Fig. S17. Long-range pRE-gene validation examples. (A)** Browser track of the region containing the lncRNA LINC02948, the gene DUSP4, and the linked pRE with epigenomic and genomic annotations in HFF cells and across ENCODE3 biosamples. **(B)** DUSP4 mRNA expression measured via RT-qPCR following individual delivery of gRNAs targeting the connected pRE in (A) in CRISPRi HFF cells (N=6 for NT, N=3 for all other gRNAs; 'NT' indicates non-targeting; 'ns' indicates not significant, adj. p-value >= 0.05; ** denotes adj. p-value < 0.01). Error bars indicate +/- 1 SEM, horizontal line indicates mean. **(C)** Browser track of the region containing the gene RFLNB and the linked pRE with epigenomic and genomic annotations in HFF cells and across ENCODE3 biosamples. **(D)** RFLNB mRNA expression measured via RT-qPCR following individual delivery of gRNAs targeting the connected pRE in (C) in CRISPRi HFF cells (N=6 for NT, N=3 for all other gRNAs; 'NT' indicates non-targeting; 'ns' indicates not significant, adj. p-value >= 0.05; * denotes adj. p-value < 0.05). Error bars indicate +/- 1 SEM, horizontal line indicates mean.

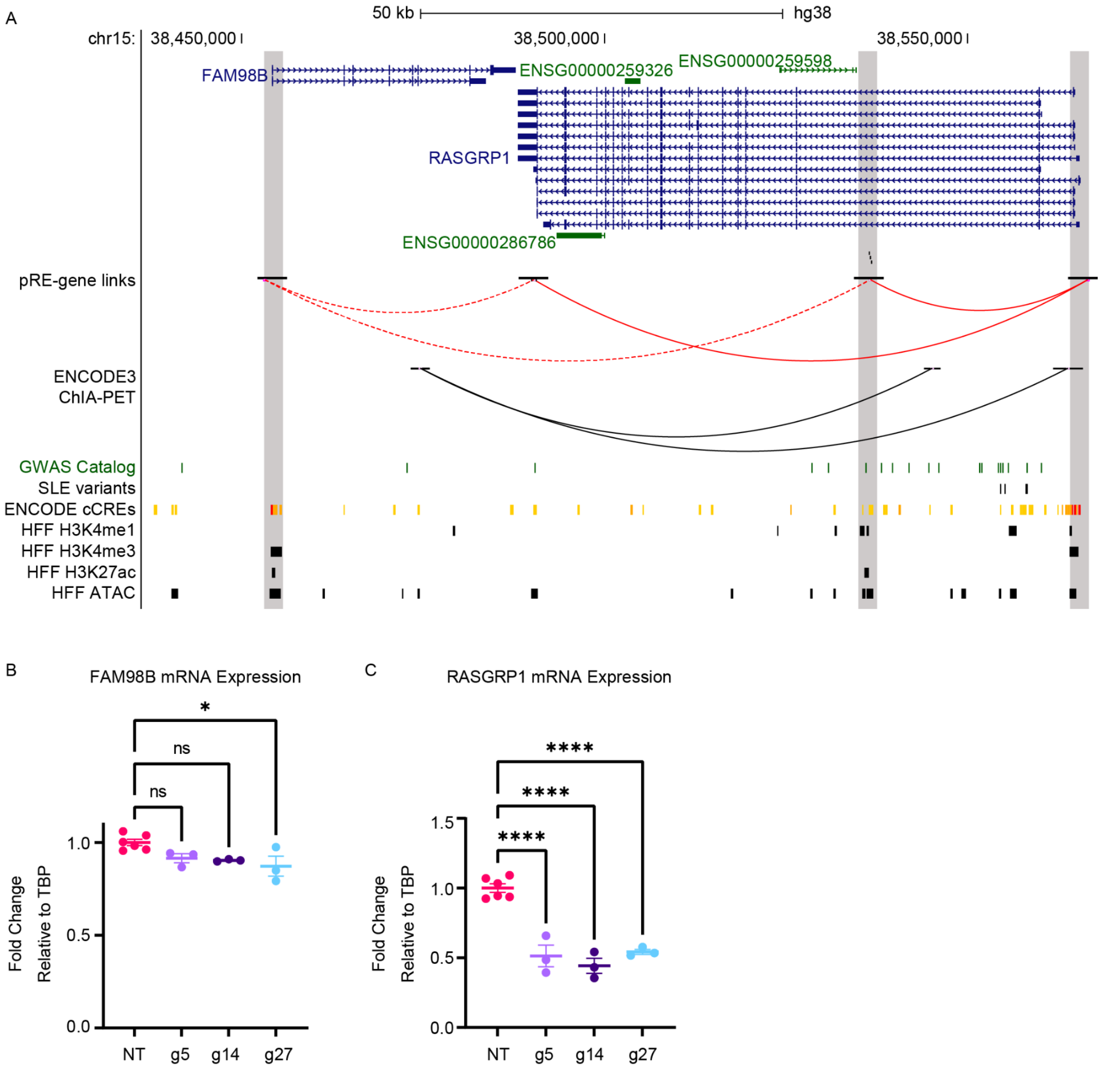


**Fig. S18. Intronic pRE regulates two genes and contains functional SLE variants. (A)** Browser track of the region containing the genes, FAM98B and RASGRP1, and the linked pRE with epigenomic and genomic annotations in HFF cells and across ENCODE3 biosamples. Single nucleotide polymorphisms from the GWAS Catalog (*47*) and SLE variants shown (*42*, *43*). Grey shading indicates promoter of FAM98B, pRE identified in scRNA-seq screen, and promoter of RASGRP1, from left to right, respectively. **(B)** FAM98B mRNA expression measured via RT-qPCR following individual delivery of gRNAs targeting the connected pRE in (A) in CRISPRi HFF cells (N=6 for NT, N=3 for all other gRNAs; 'NT' indicates non-targeting; 'ns' indicates not significant, adj. p-value >= 0.05; * denotes adj. p-value < 0.05). Error bars indicate +/- 1 SEM, horizontal line indicates mean. **(C)** RASGRP1 mRNA expression measured via RT-qPCR following individual delivery of gRNAs targeting the connected pRE in (B) in CRISPRi HFF cells (N=6 for NT, N=3 for all other gRNAs; 'NT' indicates non-targeting; 'ns' indicates not significant, adj. p-value >= 0.05; **** denotes adj. p-value < 0.0001). Error bars indicate +/- 1 SEM, horizontal line indicates mean.

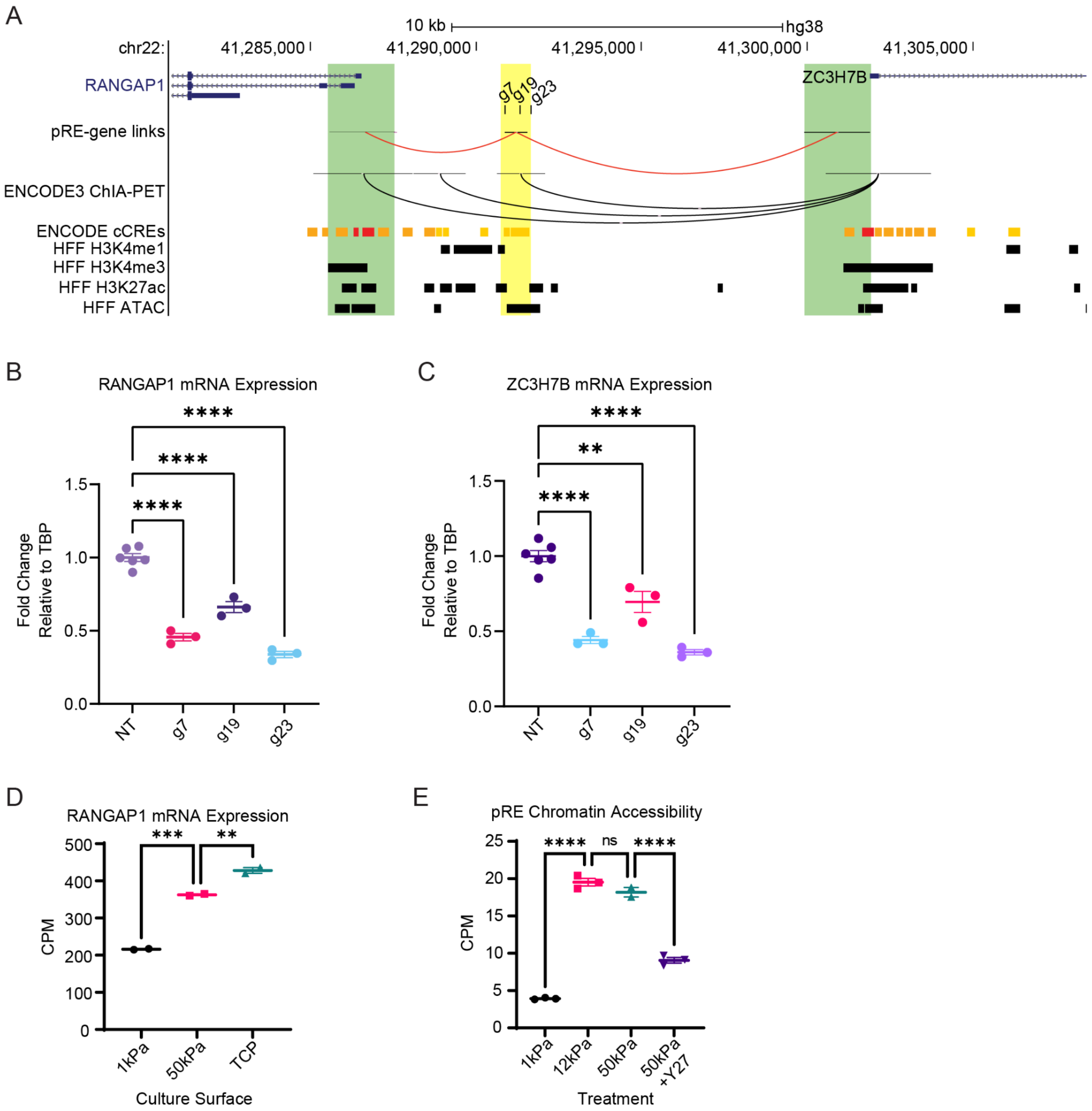


**Fig. S19. Characterization of RANGAP1 enhancer element. (A)** Browser track showing gene connections for pRE proximal to RANGAP1 and ZC3H7B. Yellow shading indicates region containing pRE and green shading highlights promoters of differentially expressed genes. ChIA-PET loops for RAD21, SCREEN cCREs, and H3K4me1, H3K4me3, H3K27ac, and ATAC-seq peaks, are also shown. Significant pRE-gene connections are drawn from the pRE (yellow shaded region) to the promoters of RANGAP1 and ZC3H7B with individual gRNA validations denoted by 'g7', 'g19', and 'g23'. **(B)** RANGAP1 mRNA expression measured via RT-qPCR for individual gRNA validations (non-targeting ('NT'), N=6; N=3 for all other gRNAs). **(C)** ZC3H7B mRNA expression measured via RT-qPCR for individual gRNA validations (non-targeting ('NT'), N=6; N=3 for all other gRNAs). **(D)** CPM values for RANGAP1 mRNA expression from bulk RNA-sequencing of HFF cells cultured on 1kPa (N=2), 10kPa (N=2), or TCP (N=2) surfaces. Individual points represent biological replicates. Error bars represent mean +/- 1 SEM. **** indicates p-value < 0.0001, *** indicates p-value < 0.001, ** indicates p-value < 0.01, * indicates p-value < 0.05.

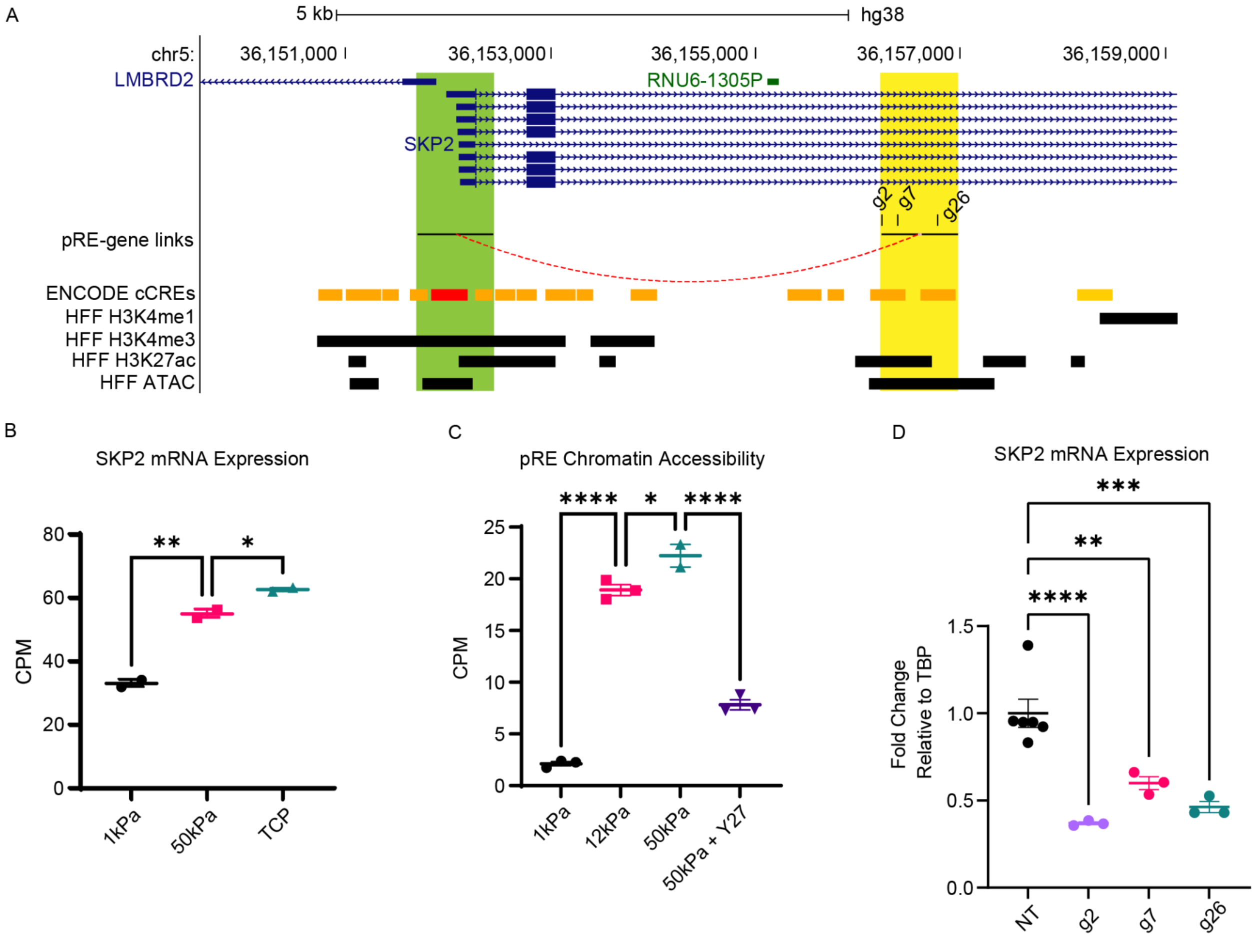


**Fig. S20. Characterization of SKP2 enhancer element. (A)** Browser track showing gene connections for SKP2 intronic pRE. Yellow shading indicates region containing pRE and green shading highlights SKP2 promoter. ChIA-PET loops for RAD21, SCREEN cCREs, and H3K4me1, H3K4me3, H3K27ac, and ATAC-seq peaks, are also shown. Significant pRE-gene connections are drawn from the pRE (yellow shaded region) to the SKP2 promoter with individual gRNA validations denoted by ‘g2’, ‘g7’, and ‘g26’. **(B)** CPM values for chromatin accessibility of SKP2 enhancer in HFF cells cultured on 10ka (N=3), 12kPa (N=3), or 50kPa (N=2), surfaces or cultured on 50kPa surface and treated with Y27 (N=3). **(C)** CPM values for SKP2 mRNA expression from bulk RNA-sequencing of HFF cells cultured on 1kPa (N=2), 10kPa (N=2), or TCP (N=2) surfaces. **(D)** SKP2 mRNA expression measured via RT-qPCR for individual gRNA validations (non-targeting (‘NT’), N=6; N=3 for all other gRNAs). Individual points represent biological replicates. Error bars represent mean +/- 1 SEM. **** indicates p-value < 0.0001, *** indicates p-value < 0.001, ** indicates p-value < 0.01, * indicates p-value < 0.05.

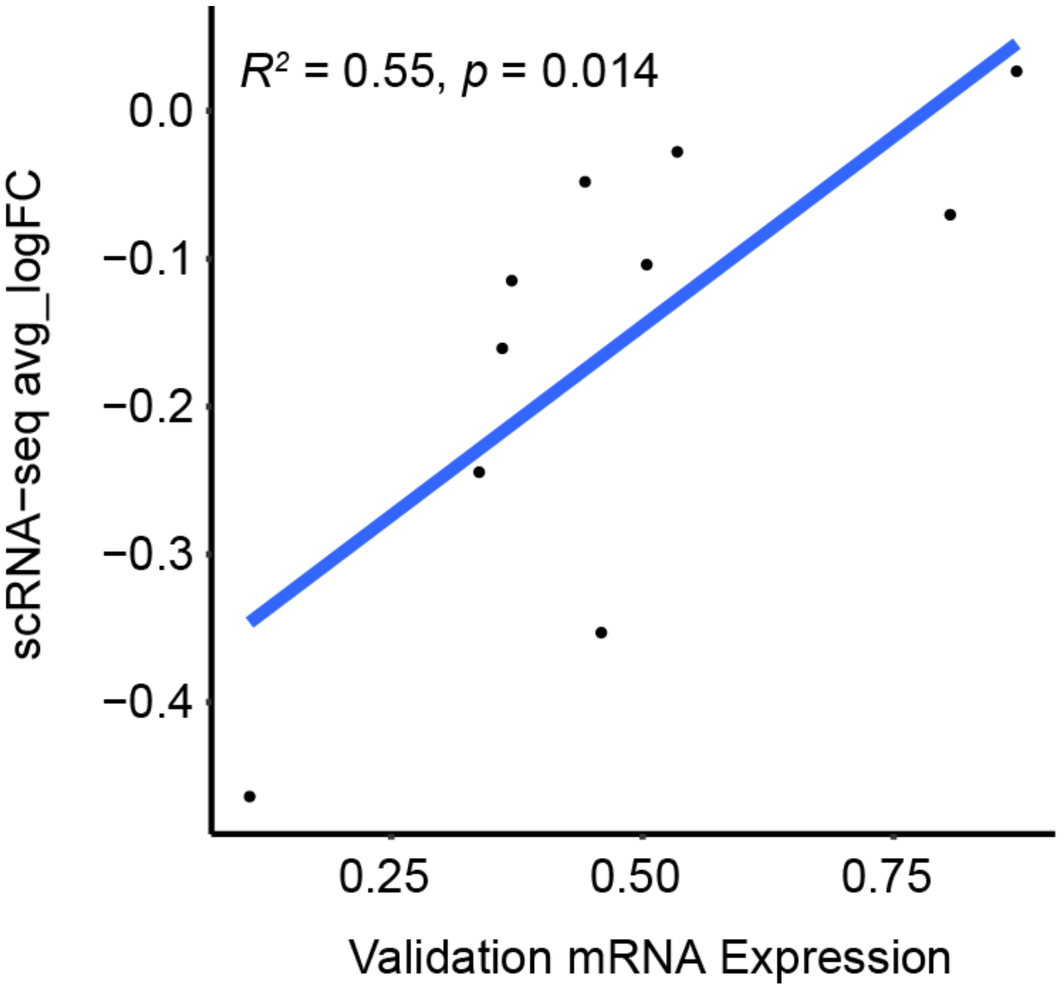


**Fig. S21. Changes in gene expression for confirmed pRE-gene connections are well correlated between single cell screen and individual gRNA validations.** Change in gene expression measured via single cell RNA-sequencing versus change in mRNA expression measured via RT-qPCR for pRE-gene connections that were confirmed with individual gRNA validations. The y-coordinate of each point is the most significant avg_logFC value for a given pRE-gene connection. Spearman correlation $R^2$ value and related significance are denoted in the upper left corner. Blue diagonal line represents the linear best-fit model.

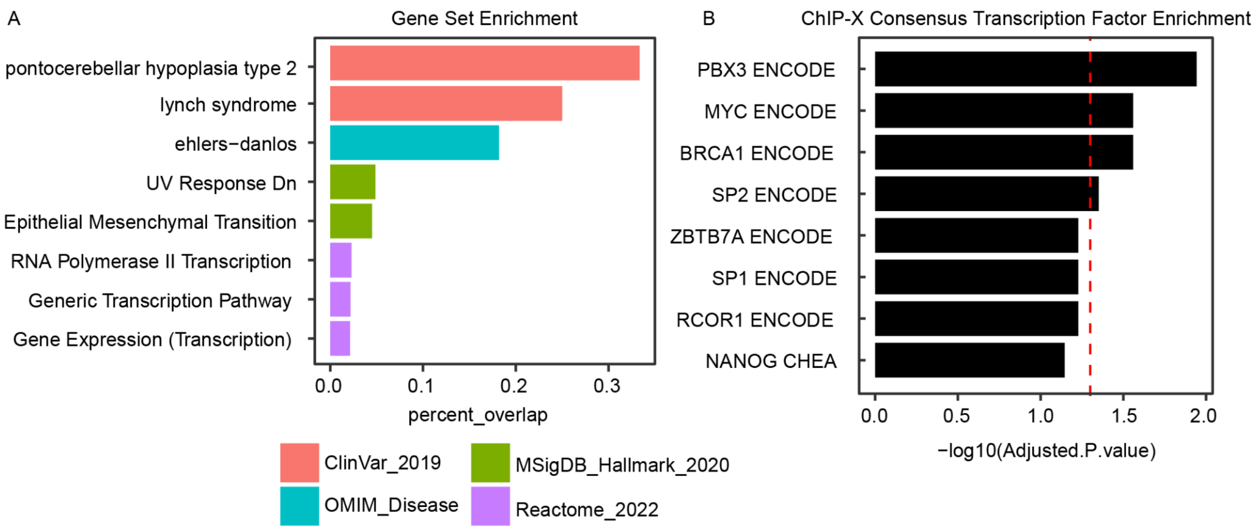


**Fig. S22. Gene set and transcription factor enrichment reveals shared networks regulated by mechanoresponsive regulatory elements. (A)** Pathway analysis using the union set of genes connected to at least one pRE link multiple mechanosensitive pathways (*48–50*). Proportion of overlap between pRE-connected genes and gene set shown on x-axis. Coloring indicates the database for each gene set. All pathways shown significant at adj. p-value < 0.05. **(B)** Transcription factor enrichment using the union set of genes connected to at least one pRE and the 'ChIP-X Consensus TF' gene set. -log10(adj. p-value) shown on x-axis. Red dashed indicates adj. p-value = 0.05. All sets shown significant at adj. p-value < 0.1.

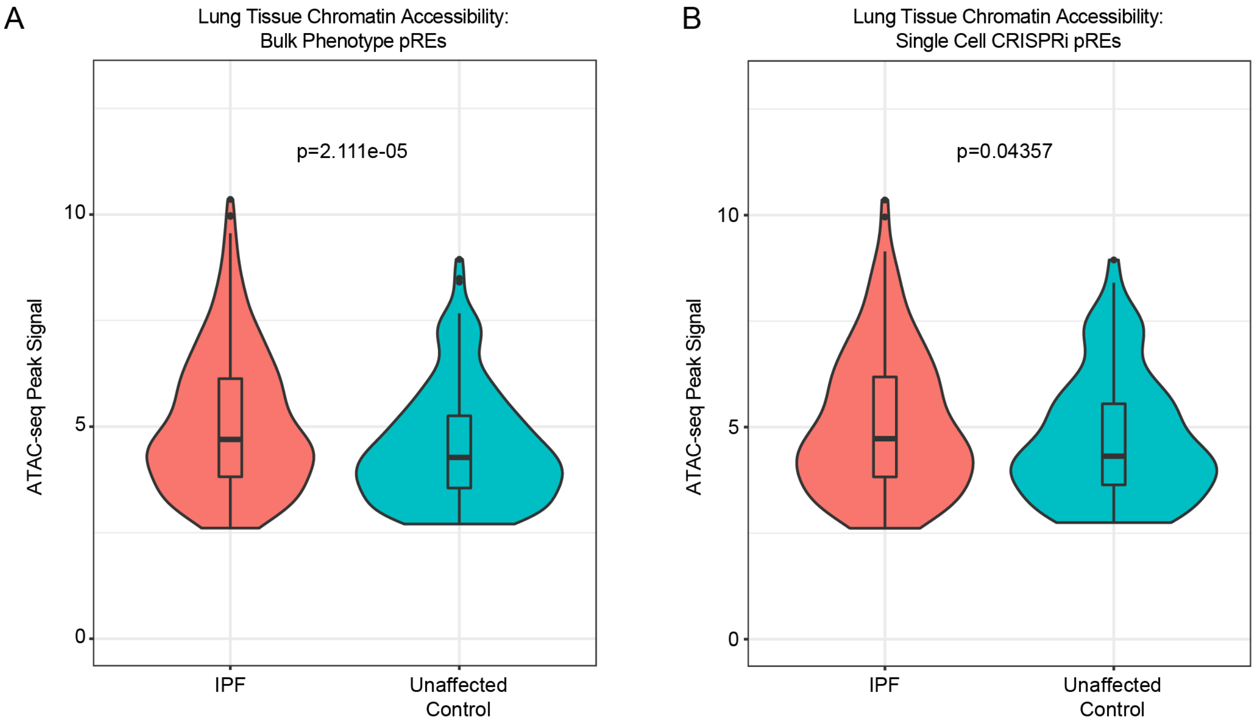


**Fig. S23. Genes regulated by mechanosensitive regulatory elements are differentially expressed in idiopathic pulmonary fibrosis. (A)** Comparison of the mean chromatin accessibility signal for ATAC-seq peaks that overlap a pRE identified in the bulk phenotype screens in lung tissue for either IPF (red) or unaffected controls (blue) (t-test, p-value = 2.111e-05; N_IPF = 303, N_Control = 196; mean_IPF = 5.084, mean_Control = 4.508). **(B)** Comparison of the mean chromatin accessibility signal for ATAC-seq peaks that overlap a pRE connected to a gene in the single cell CRISPRi screen in lung tissue for either IPF (red) or unaffected controls (blue) (t-test, p-value = 0.04357; N_IPF = 171, N_Control = 103; mean_IPF = 5.088, mean_Control = 4.699).

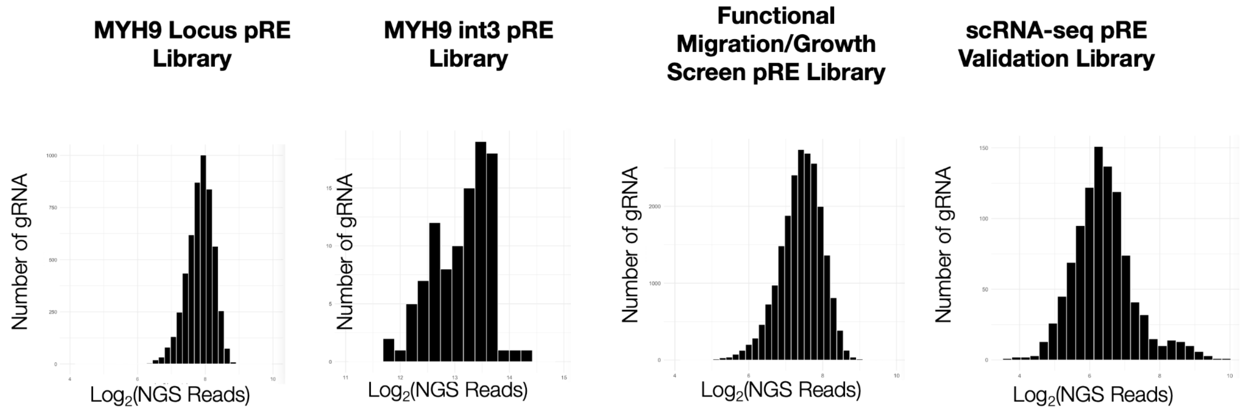


**Fig. S24. gRNA library plasmid pool verification by NGS sequencing.** For each library, a histogram is shown highlighting the log2 total reads from NGS sequencing and the number of gRNA from the library that fall in each bin. All plasmid pools showed full representation of every gRNA from the oligo pool.

**Tables S1-S19**

Table S1. ATAC-seq QC metrics.
Table S2. Differential expression results for stiff vs soft culture substrates.
Table S3. Differential accessibility results for 1kPa versus 50kPa.
Table S4. Differential accessibility results for 50kPa versus 50kPa + ROCKi.
Table S5. gRNA counts for MYH9 FACS screen.
Table S6. MYH9 FACS screen results.
Table S7. gRNA counts and results for MYH9 Cas9 tiling screen.
Table S8. gRNA counts for growth and migration screens.
Table S9. DHS-level results for growth and migration phenotype screens.
Table S10. Fisher's exact test results for overlap of significant screen regions in ENCODE biosamples.
Table S11. gRNA library for scRNA-seq screen.
Table S12. scRNA-seq screen differential expression results.
Table S13. scRNA-seq screen validation RT-qPCR results
Table S14. Pathway enrichment results for significant pRE-gene links.
Table S15. PCR primers for gDNA recovery and library preparation.
Table S16. Taqman probes used for RT-qPCR.
Table S17. Sequences for BMF luciferase assay.
Table S18. Primers used for CROP-seq recovery from 10X cDNA.
Table S19. Publicly available datasets used in this study.
